# Protocols for Multi-Scale Molecular Dynamics Simulations: A Comparative Study for Intrinsically Disordered Amyloid Beta in Amber & Gromacs on CPU & GPU

**DOI:** 10.1101/2023.10.24.563575

**Authors:** Pamela Smardz, Midhun Mohan Anila, Pawel Rogowski, Mai Suan Li, Bartosz Różycki, Pawel Krupa

## Abstract

Intrinsically disordered proteins (IDPs) present challenges to conventional experimental techniques due to their large-scale conformational fluctuations and the transient occurrence of structural elements. This work illustrates computational methods for studying IDPs at various levels of resolution. The included simulation protocol offers a step-by-step guide on how to conduct molecular dynamics (MD) simulations and analyze the results using the Amber and Gromacs packages, employing both all-atom and coarse-grained approaches. This protocol can be easily adapted to study other biomacromolecules, including folded and disordered proteins and peptides.

Furthermore, it is discussed in this work how to perform standard molecular modeling operations, such as amino-acid substitutions (mutagenesis) and insertions of residues missing in a protein structure, as well as how to incorporate post-translational modifications into the simulations, such as disulfide bonds, which are often crucial for proteins to attain their physiologically functional structure. In conventional MD studies, disulfide bonds are typically fixed at the preparation step and remain unchanged throughout the simulations, unable to break or reform. Here, in contrast, a dynamic approach is presented. It involves adequate distance restraints applied to the sulfur atoms of selected cysteine residues, allowing disulfide bonds to break and reform during the simulation.

The effectiveness of these methodologies is demonstrated by examining a model IDP, the monomeric form of 1-42 amyloid-β (Aβ42), both with and without disulfide bonds, at different levels of resolution. This study not only contributes to our understanding of the role of disulfide bonds but also provides detailed simulation protocols that can serve as a foundation for future investigations.

**SUMMARY:** Given the challenges of experimental studies on intrinsically disordered proteins, this manuscript demonstrates step-by-step protocols for conducting all-atom and coarse-grained molecular dynamics simulations using two widespread packages, Amber and Gromacs. The monomeric form of 1-42 amyloid-β (Aβ42) is used as an example, from which insights into the structure, dynamics and physicochemical properties of this protein can be obtained.

## INTRODUCTION

Proteins are one of the most important biological macromolecules, playing a variety of roles in organisms, such as serving as building blocks, catalyzing biochemical reactions as enzymes, facilitating molecular transport as transporters, and governing essential cellular processes as regulatory molecules^1^. For a considerable time, it was believed that every protein possessed a single unique structure, which was considered crucial for its functional role and believed to be deducible solely from its sequence (Anfinsen Dogma^2^). However, with advancements in science, it was discovered that many proteins at physiological conditions contain disordered regions or exist entirely in a disordered state^3^. Such systems are referred to as intrinsically disordered proteins (IDPs), although it may be more precise to classify them as ‘multi-structure proteins’ with low-energy conformational transitions or simply ‘intrinsically flexible systems’^4^. These systems are challenging to study due to their highly dynamic nature and their properties’ strong dependence on various environmental conditions, including pH, temperature, redox potential, and the presence of other molecular entities such as receptors or ligands^5^. They are also prone to oligomerization, aggregation or nanodroplet formation^6^. One example of such a system is the monomeric form of 1-42 amyloid-β (Aβ42), a fundamental component of toxic Aβ42 oligomers^7^ and fibrils associated with neurodegenerative diseases, including Alzheimer’s disease^8^. An interesting approach to expedite the study of Aβ fibril formation is the mutation of residues Leu17 and Leu34, which can be efficiently cross-linked in the fibril form, to cysteines with subsequent disulfide bond formation, forcing the peptide conformation to obtain a mature fibril structure without disturbing the fibril formation process ^9^.

Disulfide bonds are the most common post-translational modification of proteins, occurring in over 20% of proteins with known structures^10^. They fulfill a myriad of roles, including structural and enzymatic stabilization, cycle regulation, and response to stress conditions and are popularly used in industry^11^. Despite their high importance, disulfide bonds are often marginalized or even omitted in protein studies. In this work, methods for simulating the presence or absence of disulfide bonds in proteins are discussed. In addition, a novel approach allowing for dynamic formation and disruption of disulfide bonds during simulations based on finite distance restraints^12^ is presented.

Among numerous options and parameters available in molecular dynamics (MD) simulations, two of them hold paramount importance and can significantly impact the results: the choice of force field and the sampling method^13^. Together, they influence how efficiently the conformational space of the studied system is explored and whether the obtained results are reliable. While most force fields for proteins have been developed over many years primarily based on well-structured molecules, the study of IDPs necessitates the consideration of methods optimized and well-benchmarked for such systems^14, 15^. For instance, it has been demonstrated that older force fields, such as ff99SB^16^, ff14SB^17^, CHARMM22^18^, and CHARMM36^19^, yield results that more deviate from experimental data than newer parameter sets, such as ff19SB^20^ and CHARMM36m^21^. Notably, parameters for elements other than proteins, particularly water^22^ and ions^23^, are especially significant in the case of studies involving IDPs^24^. To achieve satisfactory sampling of the conformational space, more advanced techniques, such as replica exchange MD^25^ or conventional MD simulations consisting of multiple independent trajectories, each of reasonable duration, should be employed^26^. However, running simulations with more and longer trajectories can be very costly, especially for large systems comprising more than a single protein chain. Consequently, one can leverage powerful supercomputers or GPUs^27^ to accelerate classical all-atom simulations or opt for simplified models, including coarse-grained ones, in which groups of atoms are represented by single interaction centers or beads. Such simplifications not only expedite the calculation of a single MD step but also permit the use of longer time steps and, by reducing the number of degrees of freedom, smooth out the energy landscape, thereby accelerating processes within the simulation^28^. However, most coarse-grained force fields are based on statistical analyses of databases and are thus less suitable for studying IDPs, which are underrepresented in the training data^29^. One approach to address this issue is the utilization of a physics-based coarse-grained force field, such as UNRES^30^, a component of the UNICORN package^31^, or coarse-grained methods developed for IDPs, such as AWSEM-IDP^32^. Nevertheless, with the development of the Martini model^33^, the most popular coarse-grained force field to date^33^, it has been discovered that accurate scaling of interactions between solutes and solvents can be a key factor enabling MD studies of IDPs^34^. As a matter of fact, it has been recognized that MD simulations with the Martini force field typically generate IDP conformations which are more compact than those observed in experiments. Importantly, it is possible to diminish the discrepancies between simulations and experiments by carefully scaling the protein-water interactions in the Martini force field.

In this work, using the monomeric form of Aβ42 and its L17C/L34C mutant (Aβ42_disul_) as examples, it is demonstrated how to employ all-atom and coarse-grained force fields in two of the most popular simulational packages, Amber^35^ (with the all-atom ff19SB force field coupled with the OPC water model (which should perform well for both folded and disordered proteins and peptides) or with the coarse-grained SIRAH^36^ force field) and Gromacs^37^ (with the coarse-grained Martini 3 force field^33^ corrected for protein-water interactions^34^ and with the coarse-grained SIRAH^36^ force field), to study the dynamics of both folded and disordered proteins, with the option to incorporate static or dynamic treatments of disulfide bonds^38^.

## PROTOCOL

Note: For more details and explanations, the following protocol steps can be compared with descriptions in the Representative Results. Some additional explanations are also given in readme files and as comments in scripts provided in the Supplementary Materials.

This protocol is based on the use of the Ubuntu 22.04 LTS distribution of Linux, but analogous steps should be applicable using other UNIX-based operating systems. The protocol demonstrates how to efficiently run Amber MD calculations on a desktop computer equipped with a CUDA-compatible GPU as well as Gromacs MD simulations on a consumer-type CPU; however, similar calculations can be performed analogously using supercomputers or compute clusters.

The protocol consists of several options; options 1-4 are based on Amber software, while options 5-7 are based on Gromacs. Options 1-2 are for all-atom simulations, while options 3-7 demonstrate the use of software for SIRAH and Martini coarse-grained models. Options 2, 4, and 6 include mutagenesis of selected amino-acid residues to cysteines and the inclusion of a disulfide bond between them in the simulations. The application of the protocol to simulate other proteins or peptides is discussed in the Representative Results.

### 0. Prerequisites

0.1 Download the archive file from the Supplementary Materials and decompress it.
0.2 Ensure that all the required software is installed (in the case of a personal computer) or loaded in the form of modules (in the case of computer clusters or supercomputers). Some details about that are provided in the equipment table attached to this publication.
0.3 Make sure that software and packages are not only properly compiled but also installed, which may require adding some directories and libraries to operating system paths.
  0.3.1 In the case of Amber software, make sure that the “amber.sh” file is exported (by “export AMBERPATH/bash.sh” command or by adding this line permanently to the .bashrc file) before starting the procedure, as indicated in the Amber installation guide.

## Use of Amber as a Computational Engine

### Option 1

1. All-atom simulations of monomeric Aβ42 in ff19SB force field using Amber software
  1.1. Prepare the PDB input file to run MD simulations using the PDB2PQR server^39^: https://server.poissonboltzmann.org/pdb2pqr
    1.1.1. Click “Upload a PDB file” button
    1.1.2. Click “Select File” button and select the PDB file from the extracted directory
    1.1.3. Fill out the “pH field” with the requested value, in this case input “7”
    1.1.4. Select the default option “Use PROPKA to assign protonation states at provided pH“
    1.1.5. Select “AMBER” as a requested force field
    1.1.6. Select “AMBER” as a output naming scheme
    1.1.7. Select following additional options to ensure proper file processing:
      1.1.7.1. Ensure that new atoms are not rebuilt too close to existing atoms
      1.1.7.2. Optimize the hydrogen bonding network
      1.1.7.3. Add/keep chain IDs in the PQR file
      1.1.7.4. Remove the waters from the output file
    1.1.8. Click “Start Job” and wait for the completion (it usually takes less than a minute)
    1.1.9. Find a file with a “pqr” extension in “PDB2PQR Output Files” and click the “Download” button next to it to download the PQR file.
    1.1.10. Rename the downloaded “pqr” file to “***input.pqr***” and copy it to ***amber/tleap***.
1.2. Enter the directory ***Amber_all-atom_and_SIRAH***.
1.3. Execute the bash script ***amber/setup_amber.sh*** to perform system setup, energy minimization and equilibration (it can take up to one hour).
  1.3.1. Ensure that the script is working properly by checking output files at every step by manual inspection in any text editor
1.4. Execute the bash script ***amber/MD1/dyn1/start_dyn1.sh*** to start the production phase (it can take a few days).
1.5. Execute the bash script ***amber_analysis/run_analysis.sh*** to perform typical analysis of the trajectories with the use of cpptraj, a part of AmberTools package.
1.6. Execute the script ***amber_analysis/run_analysis_rg_snap.sh*** to find and extract snapshots for the all-atom simulations with the minimum and maximum *R*_g_ values
1.7. To visualize the snapshots, open PyMOL
  1.7.1. Click File from the top menu
  1.7.2. Click Open from the list
  1.7.3. Select the PDB file generated during the analysis
  1.7.4. Click A in the side menu on the right hand side of the object name, which, in this case, is the name of the PDB file without the extension
  1.7.5. Click “remove water” to make the visualization of the protein more clear
  1.7.6. Click S in the side menu on the right hand side of the object name
  1.7.7. Click cartoon to select cartoon representation of the protein
  1.7.8. Click C in the side menu on the right hand side of the object name
  1.7.9. Click “spectrum” and select the “rainbow” to use a rainbow representation to visualize the N-to C-termini of the polypeptide chain

### Option 2

2. All-atom simulations of monomeric Aβ42 with a disulfide bond in ff19SB force field using Amber software
  2.1. Enter the directory ***Amber_all-atom_and_SIRAH***
  2.2. Modify the PDB file using PyMOL software to mutate two amino-acid residues to cysteines:
    2.2.1. Open PyMOL by typing “pymol” in the terminal window.
    2.2.2. Click “S” button in the bottom right corner of the visualization window to show the protein sequence
    2.2.3. Click “Wizard” from the top window
    2.2.4. Click “Mutagenesis” button from the list
    2.2.5. Click on the amino-acid residue L17 from the displayed sequence
    2.2.6. Click on a “No Mutation” button in the right panel
      2.2.6.1. Select “CYS”
      2.2.6.2. Ensure that the “No Mutation” button changed to “Mutate to CYS”
      2.2.6.3. Click “Apply”
      2.2.6.4. Click “Done”
    2.2.7. Repeat steps 2.1.3 to 2.1.6 to mutate the amino-acid residue L34 to cysteine
    2.2.8. Click “File” from the top list
    2.2.9. Click “Save Molecule…”
      2.2.9.1. Click “OK”
      2.2.9.2. Change the directory and name the file ***mutated_disul.pdb***
      2.2.9.3. Click “Save”
    2.2.10. Close PyMOL
  2.3. Prepare the PDB file by the use of the PDB2PQR tool (i.e. execute step 1.1), copy it to folder ***amber_disul/smd_disul/tleap*** and change its name to ***input_smd.pqr***.
  2.4. Execute the bash script ***amber_disul/smd_disul/setup_smd_disul.sh*** to perform energy minimization, equilibration and steered molecular dynamics simulation to bring the sulfur atoms of the two cysteines close to each other.
  2.5. Execute the bash script ***amber_disul/tleap/cys2cyx.sh*** to convert cysteine residues (CYS to CYX) to include information about the disulfide bond between C17 and C34
  2.6. Execute step 1.1 to process the file ***amber_disul/tleap/output_smd_cyx.pdb*** using the PDB2PQR tool.
    2.6.1. After the pqr file is generated, copy it to ***amber_disul/tleap*** as **input_disul.pqr**.
  2.7. Execute ***amber_disul/setup_amber_disul.sh*** to perform system setup, energy minimization and equilibration, if one would like to use the static disulfide bond model, or ***amber_dyn_disul/setup_amber_dyn_disul.sh***, if one would like to use the dynamic disulfide model (it can take up to one hour).
    2.7.1. Ensure that the script is working properly by checking output files at every step by manual inspection in any text editor
  2.8. Execute the bash script ***amber/MD1/dyn1/start_dyn1.sh*** to start the production phase (it can take a few days).
  2.9. Run steps 1.4-1.6 to analyze and visualize the simulation results

### Option 3

3. Coarse-grained MD simulations of monomeric Aβ42 in SIRAH force field using Amber software
  3.1. Enter the directory ***Amber_all-atom_and_SIRAH***
  3.2. Perform step 1.1 to convert the PDB file to the PQR file
  3.3. Execute ***amber_sirah/setup_sirah.sh***, which converts the .pqr file using the bash script ***sirah_convert.sh*** to generate a coarse-grained structural model from the all-atom structure, and then performs system setup, energy minimization and equilibration.
  3.4. Execute the bash script ***amber_sirah/MD1/dyn1/start_dyn1.sh*** to start the production phase (it can take a few days).
  3.5. Execute the bash script ***amber_sirah_analysis/run_analysis.sh*** to perform typical analyses of the MD trajectories with the use of cpptraj, a part of AmberTools package.
  3.6. Execute the script ***amber_sirah_analysis/rg_min_max_cg_snap.sh*** to identify and extract simulation snapshots with the minimum and maximum *R*_g_ values
  3.7. Execute the VMD script ***sirah_vmdtk.tcl*** to map the simulation structures from the coarse-grained representation back to the all-atom representation. To do that perform following operations:
    3.7.1. Enter the directory ***amber_sirah/MD1/dyn1/*** for backmapping the whole trajectory or the directory ***ptraj_MD1*** (created in step 3.5 automatically) for backmapping a trajectory stripped from water and ions.
    3.7.2. Open VMD with ***sirah_vmdtk.tcl*** script using the command ***vmd -e ../../sirah_x2.2_20-08.amber/tools/sirah_vmdtk.tcl*** in the terminal
      3.7.2.1. Click “File” in the top left corner of the VMD Main window.
      3.7.2.2. Click “New Molecule”
      3.7.2.3. In the pop-up window named Molecule File Browser:
        3.7.2.3.1. Click “Browse…” and select file system.top
        3.7.2.3.2. Choose from Determine file type: AMBER7 Parm
        3.7.2.3.3. Click “Load”
        3.7.2.3.4. Click “Browse…” and select file traj.nc
        3.7.2.3.5. Choose from Determine file type: AMBER Coordinates with Periodic Box
        3.7.2.3.6. Click “Load”
        3.7.2.3.7. Choose the directory and file name, and click “OK”
      3.7.2.4. Click “Extensions” and select “Tk Console” to open a new window with VMD terminal
      3.7.2.5. Type ***sirah_backmap*** in the Tkconsolel to perform the backmapping of all snapshots in the trajectory (alternatively, type ***sirah_backmap now*** to perform the backmapping of only the currently displayed snapshot)
    3.7.3. Save the backmapped trajectory as a single pdb file.
      3.7.3.1. Click “File” in the top left corner of the VMD Main window.
      3.7.3.2. Click “Save Coordinates”
      3.7.3.3. In the newly pop-up window named Save Trajectory:
        3.7.3.3.1. Choose from the file type list: pdb
        3.7.3.3.2. Choose from Selected atoms: all
        3.7.3.3.3. Click “Save”
        3.7.3.3.4. Choose the directory and file name, and click “OK”
  3.8. Perform step 1.7 to visualize the snapshots in the all-atom representation generated in the above step

### Option 4

4. Coarse-grained MD simulations of monomeric Aβ42 with a disulfide bond in SIRAH force field using Amber software
  4.1. Perform steps 2.1 to 2.4 to mutate the selected amino-acid residues to cysteines and to convert the PDB file to the PQR format
  4.2. Execute ***amber_sirah_disul/setup_sirah_disul.sh*** to convert the .pqr file using the ***sirah_convert.sh*** script to obtain a coarse-grained structural model from the all-atom structure
  4.3. Execute the bash script ***amber_sirah/MD1/dyn1/start_dyn1.sh*** to start the production phase (it can take a few days).
  4.4. Perform steps 3.5 and 3.7 to analyze the simulation data and to convert coarse-grained structural models to all-atom structures
  4.5. Run step 1.7 to visualize the snapshots in the all-atom representation

## Use of Gromacs as a Computational Engine

### Option 5

5. Coarse-grained MD simulations of monomeric Aβ42 with solute-solvent interaction scaling in Martini force field using Gromacs software
  5.1. Enter the directory ***Gromacs_Martini/ABeta***
  5.2. Execute the bash script ***setup_scripts/setup_martini_AB.sh*** to convert the all-atom structure of Aβ42 to the corresponding Martini coarse-grained model, solvate it, and perform energy minimization (it can take up to an hour)
  5.3. Execute the bash script ***equilibration/run_equilibration.sh*** to perform equilibration simulations (it can take up to an hour)
  5.4. If using a computer cluster, execute 15 bash scripts ***production/qsub_096.sh*** to ***qsub_110.sh*** to run production simulations with the solute-solvent interaction rescaling parameter in the range from λ=0.96 to λ=1.1 (it can take up to days to complete). Alternatively, to run the simulations on a desktop computer, execute the bash script ***production/run_on_PC.sh***.
  5.5. Execute the bash script ***production/script_pbc.sh*** to perform the analysis of *R*_g_
  5.6. Execute the bash script ***production/validation.sh*** to calculate average *R*_g_ values for λ=0.96, …1.1
  5.7. Execute the bash script ***production/dmax.sh*** which calculates the maximum dimension of the Aβ42 molecule, *D*_max_, for each of the simulation structures in each of the trajectories
  5.8. Execute the bash script ***production/contactdata.sh*** which calculates contact frequency maps based on the trajectories with λ=0.96, … 1.1
  5.9. Open VMD by typing ***vmd peptide.gro run_pbc_lambda1.00.xtc*** in the terminal to visualize the MARTINI trajectories
    5.9.1. Type ***pbc box*** in the command line of VMD to display the simulation box
    5.9.2. Click “Graphics”
    5.9.3. Click “Representations”
    5.9.4. Select VDW from the Drawing Method to visualize the system (by default, the backbone beads are depicted in pink whereas the side chain beads are shown in yellow)
    5.9.5. Use the slider in the main window to move between frames
    5.9.6. Use the play button to visualize the whole trajectory
  5.10. Execute the script ***production/backmapping/backmapping.sh*** to convert the coarse-grained structures generated in the Martini simulations to all-atom structures
  5.11. Open VMD by typing ***vmd Ab42_input.pdb merged.xtc*** to display the trajectory of Aβ42 in the all-atom representation centered in the simulation box
    5.11.1. Use steps 5.9.1-5.9.6 to visualize the trajectory in the all-atom representation
    5.11.2. Click “Graphics” to change background color
    5.11.3. Click “Colors”
    5.11.4. Choose “Display” and then “Background**”** from “Categories” and set the color to white
    5.11.5. Click **File** → **Render** and choose **Start Rendering** to visualize and save a selected snapshot
  5.12. Open ChimeraX to visualize the system
    5.12.1. Click **File** → **Open** and choose one of the backmapped structures (e.g. ***production/backmapping/splitted_traj/lambda_1.03/temp/frame_100.pdb)***
    5.12.2. Go to **Favorites** → **Settings** and in the first tab **Background** change the color in **Background color** option
    5.12.3. Click **Select** → **Chains** → **chain ?** to select a whole polypeptide chain
    5.12.4. Click **Select** → **Chains** → **chain ?** and then select **Actions** → **Atoms/Bonds** → **Show** to show side-chain atoms
    5.12.5. Click **Actions** → **Atoms/Bonds** → **Atom Style** → **Ball & Stick** to show atoms of the side-chain in a different style
    5.12.6. Click **Actions** → **Cartoon** → **Hide** to hide the ribbon representation and display backbone atoms
    5.12.7. Click **File**→ **Save…** and in **Files of type** menu choose the **PNG image (*.png)**, next specify the file name in **File name** and, after setting the resolution in the **Size** option, use the **Save** button to save the image with the visualization

### Option 6

6. Coarse-grained MD simulations of monomeric Aβ42 with a disulfide bond in Martini force field using Gromacs software
  6.1. Enter the directory ***Gromacs_Martini/ABeta_CC***
  6.2. Execute the bash script ***setup_scripts/setup_martini_AB.sh*** to convert the all-atom model of Aβ42 to the Martini coarse-grained model, solvate it, and perform energy minimization with an addition of the disulfide bond (it can take up to an hour)
  6.3. Perform steps 5.3-5.12 to simulate, analyze and visualize the system

### Option 7

7. Coarse-grained MD simulations of monomeric Aβ42 in SIRAH force field using Gromacs software
  7.1. Enter the directory ***Gromacs_Sirah***
  7.2. Execute the bash script ***setup_sirah.sh*** with parameter ***Ab42*** (by typing “bash setup_sirah.sh Ab42” in the terminal) to protonate the all-atom model of Aβ42, convert it to the SIRAH coarse-grained model, solvate it, and create a topology file of the whole simulation system
  7.3. Run the bash script ***run_min_eq.sh*** with parameter ***Ab42*** (i.e. type “./run_min_eq.sh Ab42” in the terminal) to perform energy minimization and equilibration simulations (it can take few minutes)
  7.4. Execute the bash script ***run_md.sh*** with parameter ***Ab42*** (by typing “./run_md.sh Ab42” in the terminal) to run the production simulation on a desktop computer
  7.5. Run the script ***vis_trajectory.sh*** with parameter ***Ab42*** (i.e type “./vis_trajectoty.sh Ab42“ in the terminal) to visualize the SIRAH trajectory using the VMD program. NOTE: Steps analogous to those described in points 5.5-5.8 above can be performed to analyze the SIRAH simulation results.

## Use of CHARMM-GUI to Generate Input Files for the MD Simulations

### Option 8

8. Go to the official website of CHARMM-GUI: https://www.charmm-gui.org/
  8.1. Click “Input Generator” from the list on the left hand side of the website.
  8.2. Register as a non-profit user or log in with your credentials if the account is already registered.
  8.3. Click “Solution Builder” from the expanded list on the left hand side of the website.
  8.4. Click “Browse…” in the “Protein Solution System” menu and choose the file ***Amber_all-atom_and_SIRAH/amber/tleap/input.pdb***.
    8.4.1. Select the option “Check/Correct PDB Format” to ensure that CHARMM-GUI will fully process the PDB file.
    8.4.2. Write down the “JOB ID” from the top right corner of the website in case the procedure is not completed in a single use. This number can be used with the Job Retriever in the top left corner of the Input Generator menu to recover the job.
  8.5. Click “Next Step: Select Model/Chain” in the bottom right corner of the website to proceed (each step may take a few seconds to a few minutes, depending on the PDB file and the current server load).
  8.6. Check if the system has properly recognized all amino acid residues and click “Next Step: Manipulate PDB” in the bottom right corner of the website to proceed further.
  8.7. Check the option list and click “Next Step: Generate PDB” in the bottom right corner of the website to proceed further. Note: Mutations or adding disulfide bonds can be performed at this step, but it is not possible to perform both operations simultaneously. To do both, first mutate L17 to C17 and L34 to C34, then generate and download the resulting PDB in the next step, and then repeat the procedure points from 8.4 to 8.7 with the newly generated PDB file, selecting the formation of a disulfide bond.
  8.8. In the “Fit Waterbox Size to Protein Size” box, select the thickness of the water layer around the protein to 23 Å.
  8.9. Click “Next Step: Solvate Molecule” in the bottom right corner of the website to proceed further.
  8.10. Click “Next Step: Setup Periodic Boundary Condition” in the bottom right corner of the website to proceed further.
    8.10.1. Download file ***step3_pbcsetup.pdb*** file and visualize it to check if the protein structure is fine. NOTE: This PDB file can be used in the standard Amber or Gromacs procedure (options 1-7) after removing water molecules.
  8.11. Select “Amber” in the “Force Field Options:” field.
    8.11.1. Ensure that “FF19SB” is selected as the force field for proteins.
  8.12. Select “Amber” in the “Input Generation Options:” field. NOTE: Alternatively, “Gromacs” or another computational engine may be selected.
  8.13. Select “NVT Ensemble” in the “Dynamics Input Generation Options:” field.
    8.13.1. Ensure that “Temperature:” is set to 300K.
  8.14. Download the package of the generated input files by clicking “download.tgz” in the top right corner of the website.
  8.15. Extract the downloaded file.
  8.16. ***Copy*** amber/step3_input.rst7 ***as*** system.crd ***to the*** Amber_all-atom_and_SIRAH/amber/tleap directory.
  8.17. Copy ***amber/step3_input.parm7*** as ***system.top*** to the ***Amber_all-atom_and_SIRAH/amber/tleap*** directory.
  8.18. Execute the bash script ***Amber_all-atom_and_SIRAH/amber/setup_amber_ch-gui.sh*** to perform system setup, energy minimization, and equilibration (it can take up to one hour).
  8.19. Ensure that the script is working properly by checking output files at every step through manual inspection in any text editor.
  8.20. Execute the bash script ***amber/MD1/dyn1/start_dyn1.sh*** to start the production phase (it can take a few days).
  8.21. Perform steps 1.5-1.7 to analyze and visualize the simulation results.

## REPRESENTATIVE RESULTS

When following any tutorial on molecular dynamics, it is important to always consult appropriate manuals, in this protocol for Amber (https://ambermd.org/Manuals.php) or GROMACS (https://manual.gromacs.org/) as they thoroughly explain all program syntax, flags and scope of use. Users should be aware that it is often required to process or prepare the PDB file before it can be used as input for MD simulations. Although the presented results are based on a typical situation when a high-quality PDB file is used, one should take in consideration the following steps before starting MD simulations:

- An initial conformation of a peptide or protein can be downloaded as a PDB file from the PDB (https://www.rcsb.org/), PMDB (http://srv00.recas.ba.infn.it/PMDB/) or other database or source or predicted using modeling software or servers.
- All of the molecules, atoms and ions that should not be included in the simulation system need to be removed from the PDB file.
- It is usually safer to remove hydrogen atoms (by e.g. using reduce -Trim *.pdb) or to use an external tool to predict the correct protonation state for the system (PDB2PQR^39^: https://server.poissonboltzmann.org/pdb2pqr) as described in Protocol step 1.1.
- For convenience, external tools and server, such as CHARMM-GUI^40^ (https://www.charmm-gui.org/) can be used for the system generation, especially if one would like to include other molecules, such as lipids (micelles, lipid bilayers), or post-translational modifications.
- It may be necessary to rebuild missing fragments of the structure: if they are small (a couple of amino-acid residues missing), the task is trivial and can be done manually (approximation of the CL position in text editor) or using a visualization tool such as PyMOL or using any software for structure prediction (Modeller^41^, I-TASSER^42^, AlphaFold^43^); however, if a larger fragment is missing in the structure, its modeling may significantly impact the simulation results.
- If a study includes mutagenesis, like in the case of a disulfide bond addition to the monomeric Aβ42, it can be performed manually, or using a visualization tool like PyMOL (step 2.2 in Protocol), or using other modeling software.
- In same cases, non-standard amino acids could be converted to the standard ones; however, if they are necessary in the system under study or if other molecules which are supposed to be studied are not present in the force field, they have to be manually parameterized in the case of all-atom force fields or approximated by existing bead types in coarse-grained representations; for all-atom Amber force fields, AnteChamber, which is a part of Amber package, or external server, such as PyRed^44^ (https://upjv.q4md-forcefieldtools.org/REDServer-Development/) can be used.

Users must ensure that sufficient simulation time is achieved to observe the intended effects of investigation. Achieving this often necessitates the use of simplifications in both the system and its representation. To ensure proper sampling, it is advisable to routinely simulate multiple trajectories instead of relying on a single one. In some cases, the use of multiple initial conformations can expedite simulations and provide a more comprehensive scan of the conformational space. Moreover, if trajectories converge starting from various points, this convergence can serve as a check for the reliability of the simulation results. In the case of monomeric Aβ42, any method suitable for studies of IDPs should be able to reach convergence within several microseconds of the MD simulation. However, for the examination of oligomeric Aβ42, enhanced sampling methods with multiple starting points may be necessary to obtain reliable results.

It should be noted that in the case of IDPs, the convergence of simulation results cannot be defined simply as the stability of a chosen parameter (such as RMSD or R_g_) over time. Instead, it is better characterized as the system’s ability to traverse between similar conformational states multiple times, reflecting the inherently disordered nature of IDPs.

### Simulations in Amber Software

#### Software Tips

- The latest version of the proprietary NVIDIA graphics card driver is required. Alternatively, AMD graphics cards can be used if a special version of Amber is utilized (https://www.amd.com/en/technologies/infinity-hub/amber).
- CUDA Toolkit without the graphic card CUDA Driver (note: installation of drivers outside the official repository may cause problems with future upgrades of the operating system). (https://developer.nvidia.com/cuda-downloads)
- Latest version of Amber, a software package that facilitates the setup of simulation systems and enables the execution and analysis of MD simulations (https://ambermd.org/GetAmber.php) (license for academic use of both Amber and AmberTools is currently free).
  - It is important to verify the installation of all software and ensure that their versions are compatible with the current version of Amber, especially cmake, gcc and gfortran compilers, CUDA toolkit, and GPU driver. For more information, please refer to: https://stackoverflow.com/questions/6622454/cuda-incompatible-with-my-gcc-version).
  - After downloading and unpacking AmberTools and Amber files, remember to run update_amber.sh script and the adjust **build/run_cmake** script, e.g. change the flag “DCUDA=TRUE” to enable CMake to install pmemd.cuda, a program that allows running programs on the GPU. Since not all programs are GPU-compatible, it is recommended to first compile the code on the CPU and then on the GPU.
  - If one would like to use multiple CPU cores for MD simulations in the Amber package, another compilation with the flag “DMPI=TRUE” should be performed, and pmemd.MPI used instead of pmemd to run the simulations. In this case, ensure that MPI or OpenMP is installed.

#### Representative Results of Option 1: All-Atom MD Simulations of A**β**42

The first step of every simulation procedure presented in this publication is to process the input PDB file with the use of the PDB2PQR server^39^ (https://server.poissonboltzmann.org/pdb2pqr) as presented in Protocol step 1.1. Note that a standalone version of this software can be used instead (https://github.com/Electrostatics/pdb2pqr).

The second step (Protocol step 1.2) is to generate the topology and coordinate files based on the selected force field:

- The initial structure of Aβ42, as given in the PDB file ***Ab42_input.pdb*** (due to the disordered character of Aβ42, there are no experimentally determined structures of its monomeric form in solution; here, one of the conformations obtained in previous MD studies is used as the initial model^13^) is converted to Amber topology and coordinate files using the **tLeap** program with the ff19SB force field for proteins.
- The protein is solvated with the OPC water and co-optimized ions (sodium and chloride) are added to neutralize the system and to reach the physiological salt concentration of about 0.15 M. Counter-ions are calculated automatically if *-1* number is specified in the tLeap input file (e.g. “addions2 system Na+ -1”), but salt concentration needs to be inserted manually. For additional details on setting up the salt concentration, please refer to the Amber tutorial available at: (https://ambermd.org/tutorials/basic/tutorial8/index.php) or, alternatively, on the CHARMM-GUI site, the input generator tool allows one to calculate the salt concentration (https://www.charmm-gui.org/).
- The system is placed in a selected type of a periodic boundary condition (PBC) box (in this case: truncated octahedron), the size of which is determined based on the added water layer.

The provided scripts for all-atom simulations in Amber utilize a default set of the protein force field (ff19SB) coupled with the recent and most versatile water model (OPC). Despite the higher computational cost of using the four-point OPC water model, it is a recommended model to be used in Amber all-atom force fields, as it provides much better results if solute-solvent interactions are important. This is especially true for IDPs, such as the monomeric Aβ42, when compared to classical three-point models, such as TIP3P. It should be noted that users can easily switch between various force field versions and water models by simply modifying the top lines in the leap.in file according to the official Amber website and a manual or publications of unofficial force field versions, such as, for example, ff14IDPs ^45^, designed specifically for IDPs.

Once the simulation system is set up, the energy minimization is performed. Parameters of the energy minimization procedure are specified in the input file ***md_min.in***, and the bash script ***start_min.sh*** is used to copy the generated topology and coordinates files and to run the minimization. During the energy minimization, the optimization method is switched from the steepest descent to the conjugate gradient, combining stability with efficiency. Restraints are typically imposed to maintain the initial positions of solute heavy atoms.

If the energy minimization cannot be executed successfully, one should check for any warnings or issues in the tLeap log file (leap.log) and consider increasing the number of minimization steps in both algorithms. In the rare event that the steepest descent calculation gets stuck in a high-energy local minimum, the subsequent conjugate-gradient calculation may fail. In such a situation, one may try to perform only the steepest descent minimization followed by a very short equilibration run (below ps) with a significantly reduced time step (several orders of magnitude shorter than usual) to move the system out from this conformation. It is advisable to repeat the standard minimization procedure afterward.

The system is equilibrated in two successive steps. In the first one, equilibration is performed using the CPU to increase stability in the NPT ensemble. The second step utilizes the GPU to expedite the process without risking instabilities. For the first equilibration run (on CPU), all simulation parameters are specified in the input file ***md_eq1.in***. The bash script ***start_eq1.sh*** is used to initiate the first equilibration run. For the second equilibration run (on GPU), the input file ***md_eq2.in*** and the bash file ***start_eq2.sh*** are used. The simulation time step is set to 2 fs, and the total equilibration time is 1 ns, which should be adequate for a typical system without a lipid bilayer to reach the proper temperature and density. This time can be increased if necessary. During the equilibration process, initial velocities are generated, and the system is heated up to the production temperature (300K) to prevent potential steric clashes. In systems with multiple components, such as ligands or lipids, restraints are applied to groups of atoms to ensure a proper system preparation and to prevent component dissociation resulting from minor structural imperfections that could lead to localized high energy. Equilibration in the NPT ensemble establishes the correct system density, allowing the production phase of the simulation to proceed in the NVT ensemble. A cutoff distance of 0.9 nm is set, and the simulation uses a Langevin thermostat with a collision frequency of 2 ps^-1^ and a Berendsen barostat with isotropic position scaling. To calculate the long-ranged electrostatic interactions, the Particle-Mesh Ewald (PME) method is used, which gives a speed-up to the calculations. For GPU calculations on NVIDIA GPUs, one can use the ‘CUDA_VISIBLE_DEVICES’ flag to select which GPU unit to use when multiple GPUs are available. Additional equilibration with the use of GPU can be performed. To do this, users need to create a directory named ‘**eq3**’, copy the contents of the ‘**eq2**’ directory into ‘**eq3**’, and rename the bash script ‘**start_eq2.sh**’ to ‘**start_eq3.sh**’. Then modify the script to initiate the simulation from the restart file from the second equilibration run: ‘**-c system_eq2.crd**’, and generate a third restart file: ‘**-r system_eq3.crd**.’ Finally, execute the bash script ‘**start_eq3.sh**’ to run the third equilibration simulation.

The scripts ***start_min.sh***, ***start_eq1.sh*** and ***start_eq2.sh*** copy the necessary input coordinate and topology files from the respective subdirectories of the previous steps. They then initiate simulations that generate trajectory files, which contain coordinates of the system (snapshots) saved at specified intervals, a ‘restart’ file with coordinates and velocities (after equilibration), output with simulation information, and details regarding the simulation’s progress.

The final simulational step (see Protocol step 1.4) is to run production MD simulations. All the parameters for the MD simulation are specified in the input file ***md_dyn1.in***. In this phase, the simulation time step is set to 2 fs for all-atom simulations, and the production time is 1 μs. For the coarse-grained simulations, the simulation time step is set to 20 fs, and the production time is 20 μs. To enhance the simulation speed and stability, the NVT ensemble is typically used for production trajectories (https://ambermd.org/GPULogistics.php). The cut-off distance remains at 0.9 and 1.2 nm for all-atom and coarse-grained simulations, respectively, and Langevin dynamics with a collision frequency of 2 psL¹ at a temperature of 300K are applied. The only difference compared to the equilibration phase is the absence of pressure control since a constant volume, not pressure, is maintained. The SHAKE algorithm is used to constrain hydrogen bond lengths to allow for a longer time step and higher stability of the simulation.

If one needs to extend the simulations, navigate to the folder ***MD1/dyn2***, set the desired production steps using the ‘nstlim’ flag, and execute the bash script ***start_dyn2.sh***. This extension can be performed multiple times, but make sure to use appropriate file naming. To run multiple trajectories, duplicate the input folder ***MD1*** (i.e ***MD2***, ***MD3***, etc.) as many times as needed (a reasonable minimum for averaging is 3 trajectories) and follow analogous steps as for MD1.

During the simulations, the following files are generated:

- **traj.nc** file containing atom/bead coordinates,
- **system_dyn*.nc** restart file with coordinates and velocities, which allows to restart the simulation,
- **min.out** file with simulation information output,
- **mdinfo** file with information about the simulation progress, speed of the simulation and the estimated time to finish the production run.

After the simulations are finished, there are several analyses that allow one to study structural properties of the system based on the MD trajectories. The bash script ***amber_analysis/run_analysis.sh*** performs the following operations (see Protocol step 1.5):

- Creates folders for analysis for each trajectory listed in the script, and runs the ***cpptraj*** program with commands provided in the input file ***ptraj_traj.in***. This process generates topology and coordinate files in the **ptraj_MD\*** folders after centering the solute in the periodic boundary condition box (using the ‘autoimage’ command), followed by the removal of water molecules and ions. Caution while using the ‘autoimage’ command in more complex systems is recommended, as it may cause unwanted shifts of solute molecules. This step is important for speeding up all the following analyses. (Note: if there is any analysis that needs to incorporate initial coordinates or water molecules and ions, it should be run on the initial coordinate and topology files).
  - Be careful not to run any further analysis in a single script if the RMSD calculation has been performed as it may impact the results. Conversely, be cautious about the orientation of the solute in the PBC box when calculating RMSF, or other analyses requiring superpositioning with a reference structure.
- Runs **cpptraj** program with the commands included in the input files **ptraj_rmsd_rg.in**, ***ptraj_pdb.in*** and ***ptraj_image_distance.in***. ***Ptraj_rmsd_rg.in*** calculates RMSD, *R*_g_ and RMSF; ***ptraj_pdb.in*** creates PDB files from Amber topology and trajectory files, and ***ptraj_image_distance.in*** calculates the minimum non-self imaged distance, which is used to determine if the periodic box is of sufficient size to prohibit interactions between periodic images.
- Copies all the resulting text and PDB files with overwritten names of each trajectory to the corresponding folders (rg, rmsd, rmsf, pdb, mindist).

To visualize snapshots for the all-atom simulations with the minimum and maximum *R*_g_ values, the batch script ***amber_analysis/rg_min_max_snap.sh*** perfors following operations (see Protocol step 1.6): is searches for the minimum and maximum values of *R*_g_ and identifies the corresponding structures; then it extracts those structures from the trajectory file and saves them in the pdb format. Visualization is presented in Protocol step 1.7.

#### Representative Results for Option 2: All-Atom MD Simulations of A**β**42 with an Artificially Added Disulfide Bond

The second set of the MD simulations in Amber is performed with the disulfide bond present in the monomeric Aβ42. However, as this system does not include cysteines that can form disulfide bonds in the native state, two amino-acid residues have to be mutated to cysteines. Note that in Amber software when simulating a protein that contains disulfide bonds in its wild type form, the amino-acid three-letter code in the PDB file must be changed from CYS to CYX to reflect these bonds. Additionally, the tLeap input file should include a line specifying which sulfur atoms are bonded together. If the simulation aims to reduce or remove these disulfide bonds, the amino-acid code should remain as CYS.

Mutations in the PDB file can be performed manually with a text editor or using visualization, modeling, or structure prediction tools. If a single point mutation is planned, then the fastest approach should be to use a manual method, however, if more extensive modifications are planned, it may be necessary to use more advanced methods such as I-TASSER or AlphaFold, as described briefly below.

##### • Manual mutation

To manually mutate any amino acid residue in the PDB file, except for Gly, remove all atoms from the side chain except for the CB atom and change the three-letter code of the residue to the desired amino acid. For Gly, change the amino-acid code for the CA atom only. During system preparation, tLeap will automatically assign any missing atoms. Be cautious as this process may lead to steric clashes; therefore, perform the minimization and equilibration steps with extra care.

##### • Visualization-tool mutation (PyMol)

PyMOL Mutagenesis Wizard tool can be used to change any of the amino-acid residues (Protocol step 2.1). The tool also allows the user to choose a rotamer, but it’s important to note that they may still result in steric clashes (for more information see: https://pymolwiki.org/index.php/Mutagenesis).

##### • Use of structure prediction tool

If extensive mutations or deletions are planned, using advanced software like I-TASSER^46^ or AlphaFold^43^, which predicts structures based on provided sequence and known protein structures, may be advisable. Such tools not only have extensive checks to avoid steric clashes caused by the mutation but, to some extent, can also incorporate local or even global conformational changes upon mutation; therefore, they should provide more reliable initial structures than manual modifications.

#### Insertion of a disulfide bond

##### • Selection of the conformation with cysteine residues in the proximity

In some cases, there may be a conformation in the trajectory of the wild-type protein where amino acid side-chains that need to be mutated are in close proximity. To check this, one can use cpptraj with the ‘distance’ command to measure the distance between the desired CB or CG atoms. If a suitable conformation is found, it can be extracted and then mutated as described above to create the initial conformation.

##### • SMD of a structure to bring cysteines together

If the previous method fails or if there is no trajectory without mutation available, another option is to run a short steered MD (SMD) simulation (Protocol step 2.3). After mutating the chosen amino-acid residues to cysteines, the initial steps (system setup, minimization, and equilibration) should follow the same procedure as for the wild-type version. Then, run a short production run of SMD with distance restraints imposed on the sulfur atoms in cysteine side-chains, reducing the distance between them to 0.2-0.3 nm. Convert the last snapshot to a pdb file and, after verifying the structure with PyMOL or other molecular visualization software, use it as the initial conformation. The corresponding scripts and input files can be found in the directory ***amber_disul/disul_smd***.

##### • Selection of a disulfide-bond model

There are two options for applying disulfide bonds in MD simulations using the Amber software: one can simply use covalent bonds, which can be present or absent throughout the entire simulation (static treatment), or one can use a pseudopotential that allows for breaking and reforming disulfide bonds during the simulation (dynamic treatment).

The static disulfide-bond treatment is performed by the bash script ***amber_disul/setup_amber_disul.sh*.** This script performs operations analogous to those described for simulations in Amber without disulfide bonds with two modifications: (i) The initial structure in the PDB file ***amyloidB_withSS.pdb*** is used as input. This structure of Aβ42 contains a disulfide bond between Cys17 and Cys34, indicated as CYX in the PDB file. (ii) The tleap program has an additional command ***bond*** to set disulfide bonds between the sulfur atoms. Tleap detects and incorporates the disulfide bonds into the topology file.

The dynamic disulfide-bond treatment is performed by the bash script ***amber_dyn_disul/setup_amber_dyn_disul.sh.*** This script performs operations analogous to those described for simulations in Amber without disulfide bonds with three modifications: (i) The initial structure in the PDB file ***amyloidB_dynSS.pdb*** is used as input. This structure of Aβ42 contains a disulfide bond between Cys17 and Cys34 which is indicated as ‘CYS’ in this PDB file, in contrast to the static disulfide bond ‘CYX’ version. (ii) There is a third equilibration step with restraints placed on the sulfur atoms to mimic a breakable bond. (iii) Both the third equilibration and production steps have an additional input file that indicates the sulfur atoms that are restrained.

It is important to note that this approach represents a heavily simplified description of disulfide bond formation and disruption, and thus should be used with caution. Classical all-atom force fields cannot capture chemical reactions and, therefore, the hydrogen atom in the thiol group of cysteine remains present throughout the simulation. Nevertheless, we have found this approach to be effective in both the bound and unbound states of cysteines, exemplified by bovine ribonuclease A^12^.

#### Representative Results of Options 3 and 4: SIRAH Coarse-Grained MD Simulations of A**β**42 without and with Disulfide Bonds

The cash script ***amber_sirah/setup_sirah.sh*** (see Protocol step 3.2) converts the .pqr file using ***sirah_convert.sh***. The .pqr file can be prepared with charge assignment using the PROPKA tool, for example, *via* the PDB2PQR server (protocol step 1.1). The pqr files are ready to be used as input for tLeap, with syntax analogous to that of all-atom simulations. The only changes required are in the names of the protein and water parameters, which are also explained in the ***README*** file provided by the SIRAH developers.

Similarly, the bash script ***amber_sirah_disul/setup_sirah_disul.sh*** (see Protocol step 4.2) uses the coarse-grained SIRAH force field and model with the Aβ42 double Cys mutant based on the PQR file. In the case of the coarse-grained simulations using ***amber_sirah_disul***, the tleap input includes an additional command to set a disulfide bond between the appropriate coarse-grained beads.

The bash script ***amber_sirah_analysis/rg_min_max_cg_snap.sh*** (see Protocol step 3.5) is used to perform the analysis for the visualization of the SIRAH coarse-grained simulation results. This script performs operations analogous to those from the all-atom procedure with two modifications: (i) The extracted structures are saved in Amber topology and coordinate files. (ii) The VMD script ***sirah_vmdtk.tcl*** is used to map the simulation structures from the coarse-grained representation back to the all-atom representation. This script is provided by SIRAH developers, and is included in the SI file in the Amber_all-atom_and_SIRAH/sirah_x2.2_20-08.amber/tools directory. After opening VMD with the trajectory file, the backmapping is performed with the command ***sirah_backmap***, which backmaps every snapshot in the trajectory, or with the command ***sirah_backmap now***, which backmaps only the current snapshot.

### Simulations in Gromacs Software

Access to a computer cluster is useful but not necessary to perform the Martini simulations. The MD simulations can be run on a desktop computer with a multi-core CPU. However, the main advantage of using a computer cluster is that many simulations can be run simultaneously on separate compute nodes. Here, we provide bash scripts to execute the Martini simulations on a computer cluster equipped with the Slurm queue system and for simulations run on a desktop computer. On a supercomputer, or computer cluster, the installation of necessary software is typically performed by the system administrator and can be loaded using the “module load” command.

#### Representative Results of Option 5: Martini Coarse-Grained MD Simulations of A**β**42

General steps to run simulations in Gromacs are similar to the ones in Amber. The bash script ***setup_scripts/setup_martini_AB.sh*** (see point 5.2 in the protocol) performs the following operations:

- The initial structure of Aβ42, as given in the PDB file ***Ab42_input.pdb***, is converted to a coarse-grained representation using the martinize2 program. Martini topology and structure files are generated based on the information from the provided PDB file. It is important to note at this point that if one would like to use another version of the Martini force field, one simply has to copy the force-field parameter files to the directory ***Gromacs_Martini/ABeta/setup_scripts*** and update the ***setup_martini_AB.sh*** script accordingly.
- The coarse-grained protein structure is placed in a cubic box of side length of 9 nm using the Gromacs editconf tool.
- Protein is solvated using the Python script ***insane.py***. Sodium and chloride ions are added to neutralize the system and to reach the physiological salt concentration of 0.15 M. Note that flag “-salt” applied to the ***insane.py*** script indicates the salt concentration in molar units.
- Energy minimization is performed. Parameters of the energy minimization procedure are specified in the Gromacs input file ***minimization.mdp***.
- The force-field parameters describing non-bonded interactions between the protein and water beads are re-scaled by a factor λ using the bash script ***rescaling.sh***. The values of parameter λ are set here to 0.96, 0.97 … 1.10.

The goal of performing a series of MD simulations with different λ-values is to determine an optimal value of parameter λ which yields the best agreement between the simulation and experiment^34^. The range of values of parameter λ can be changed in the bash script ***rescaling.sh***. Note that λ = 1 corresponds to unscaled solute-solvent interactions, i.e. to the original Martini 3 force field.

The bash script ***equilibration/run_equilibration.sh*** (see point 5.3 in the protocol) performs equilibration simulations. The same range of λ-values as in the system setup step described above (i.e. λ = 0.96, 0.97 1.10) is used here. All the parameters of the equilibration simulations are specified in the Gromacs input file ***equilibration.mdp***. Importantly, the integration time step is set to 2 fs to allow for slow relaxation of the system. The equilibration time is set to 10 ns, which should be enough for typical protein systems to reach the specified temperature and density.

In the first step, the script ***run_equilibration.sh*** generates a binary tpr file that contains information about all the simulation parameters (including force field parameters), about the system topology, and about the starting coordinates and velocities of all the beads forming the simulation system. Here, the starting velocities are generated on the basis of the Maxwell-Boltzmann distribution with the specified temperature. In the second step, the MD simulation is started using the tpr file as input. The simulation box size after the equilibration can be checked at the end of the ***eqb1.00.gro*** file. Here, the simulation box is a cube of the side length of 9.23 nm.

Production runs are performed by executing the bash scripts ***qsub_096.sh*** … ***qsub_110.sh*** (see point 5.4 in the protocol). Each of the scripts starts one simulation with a specified λ-value, ranging from λ=0.96 to λ=1.1. All the simulation parameters are given in the Gromacs input file ***dynamic.mdp***. Here, the integration time step is set to 20 fs and the total simulation time is 20 μs. The temperature and pressure are kept constant at T = 300 K and p = 1 bar, respectively, using the velocity-rescaling thermostat and the Parrinello-Rahman barostat. Nonbonded interactions are treated with the Verlet cutoff scheme. The cutoff for van der Waals interactions is set to 1.1 nm. Coulomb interactions are treated using the reaction-field method with a cutoff of 1.1 nm and dielectric constant of 15. Frames are saved every 1 ns.

Similarly as in the equilibration procedure, the scripts ***qsub_096.sh*** … ***qsub_110.sh*** firstly generate binary tpr files containing all the information needed to start the simulations (i.e. system topology, MD simulation parameters, and starting coordinates and velocities of the beads). Then the MD simulations are started using the tpr files as input. Each of the production simulations takes about two days on a 12-core compute node.

The following files are generated in the course of the simulations:

- ***run_pbc_lambda*.gro*** files containing the Cartesian coordinates of the beads at the final step of the simulation.
- *run_pbc_lambda*.xtc* trajectory files.
- ***run_pbc_lambda*.edr*** binary files that contain time-series non-trajectory data such as energy breakdowns, temperature and pressure of the simulation system.
- ***run_pbc_lambda*.log*** files containing information about the overall performance of the simulation. These files also include energy breakdowns as well as the instantaneous values of the pressure and temperature of the simulation system.
- ***run_pbc_lambda*.cpt*** binary files required for restarting the simulation. The complete information about the system state is stored in these checkpoint files.

After the production simulations are completed, bash the script ***production/script_pbc.sh*** (see point 5.5 in the protocol) performs the following tasks: Firstly, the Gromacs trjconv tool with the flags “-pbc mol -center” makes the Aβ42 peptide centered inside the simulation box, taking into account the periodic boundary conditions. The resulting trajectory files are named ***run_pbc_lambda*.xtc*** and moved to directories ***AB_\**** for all the simulated values of parameter λ. Secondly, the radius of gyration of the Aβ42 molecule, Rg, is calculated for all of the simulation structures and recorded in output files rg_run_pbc_lambda*.xvg, which are moved to folder***/analysis_results/RG***. It should be noted that the radius of gyration is calculated here using the coordinates of all the beads forming Aβ42.

Next, the bash script ***production/validation.sh*** (see point 5.6 in the protocol) calculates the average radius of gyration for each of the trajectories with λ = 0.96, …, 1.1. The statistical errors are estimated using block averaging. Namely, a given trajectory is divided into 10 blocks of 2 μs each, and the average value of *R*_g_ is calculated for each of the blocks. The standard deviation of these black-averaged *R*_g_ values estimates the statistical error of the average value of *R*_g_ in the whole trajectory.

Then the bash script ***production/dmax.sh*** (see point 5.7 in the protocol) calculates the maximum dimension of the Aβ42 molecule, *D*_max_, for each of the simulation structures in each of the trajectories. *D*_max_ is defined as the distance between two farthest beads of Aβ42. The *D*_max_ values are recorded in files ***dmax*.xvg*** and moved to folder ***dmax***.

Finally, the bash script ***production/contactdata.sh*** (see point 5.8 in the protocol) calculates contact frequency maps based on the ***run_pbc_lambda*.xtc*** trajectory files. The contact frequency maps are recorded in files ***contact*.xvg*** and moved to folder ***contactmap***. The only parameter in the contact map calculation is the cutoff distance below which a contact is assumed to exist between a pair of beads. Here, the default value of 0.7 nm is used (flag cutoff=0.7 in the Python script ***contactdata.py***). Thus, if the distance between any bead of a given amino-acid residue and any bead of another amino-acid residue is smaller than 0.7 nm, then this residue pair is deemed to be in contact.

The Aβ42 conformations sampled in the Martini simulations and saved in the trajectory files ***run_pbc_lambda*.xtc*** can be conveniently visualized using the VMD program (see point 5.9 in the protocol). In addition, the bash script ***production/backmapping/backmapping.sh*** (see point 5.10 in the protocol) converts the coarse-grained structures generated in the Martini simulations to all-atom structures. This script is based on the backmapping algorithm introduced by Wassenaal et al.^47^ and performs the following operations:

- An all-atom topology file is created using the PDB file ***Ab42_input.pdb*** and the CHARMM36 force field.
- A single coarse-grained structure is extracted from a given trajectory file run_pbc_lambda*.xtc.
- The coarse-grained structure is mapped back to an all-atom structure using the ***backward.py*** program. This backmapped structure is relaxed in several short cycles of energy minimization and position-restrained MD simulations with the CHARMM36 force field.
- The two points above are iterated over all the coarse-grained structures obtained from the Martini simulations, combining the resulting all-atom structures into a single trajectory file.

The backward program consists of several scripts and mapping definition files, including the Python script ***backward.py*** which conducts the actual backmapping; the bash wrapper ***initral-v5.sh*** that performs the energy minimization and position-restraint MD simulations using Gromacs with the CHARMM36 force field; files ***\*.map*** containing definitions of the mapping between the all-atom and coarse-grained representations; and the Python script ***init .py*** to interpret the map files.

It should be noted that the all-atom structures can be used not only to visualize the simulation trajectory (see points 5.11 and 5.12 in the protocol) but also to perform analyses analogous to those in points 5.5, 5.6, 5.7 and 5.8 of the protocol.

#### Representative Results of Option 6: Martini Coarse-Grained MD Simulations of A**β**42 with an Artificially Added Disulfide Bond

The bash script ***ABeta_CC/setup_scripts/setup_martini_AB.sh*** (see point 6.2 in the protocol) sets up the Martini simulations of the Aβ42 Leu17Cys/Leu34Cys mutant, denoted here by Aβ42_disul_. This script performs operations analogous to those described in the previous subsection for the Aβ42 variant without the disulfide bond, with two modifications: (i) The PDB file ***amyloidB_withSS.pdb*** with a Aβ42_disul_ structure is used as input. This structure contains a disulfide bond between Cys17 and Cys34. (ii) The martinize2 program is called with the flag “-cys auto”. With this flag, martinize2 automatically detects and incorporates disulfide bonds into the topology file as constraints on the distance between the cysteine side-chain SC1 beads. Here, the distance between the SC1 beads of Cys17 and Cys34 is constrained at 0.24 nm using the LINCS algorithm^48^. Importantly, accurately including disulfide bonds in the input structure may require a high-resolution initial structure or preprocessing, such as all-atom energy minimization.

#### Application of Martini Coarse-Grained MD Simulations to Structured and Mix-Folded Proteins

It should be noted that also proteins with well-defined structures can be simulated using the Martini coarse-grained model with just a few modifications to the protocol presented in this work. To set up Martini simulations of a protein with a well-defined structure, it is necessary to use an elastic network^49^, i.e. a set of harmonic restraints (or so-called “rubber bands”) that ensure that the secondary and tertiary structure of the protein is preserved in the course of the simulation. To this end, the martinize2 program should be called with additional flags, e.g. “-elastic -ef 700.0 -el 0.5 -eu 0.9 -ea 0 -ep 0 -scfix”. This imposes harmonic restraints between backbone beads within the lower and upper distance cut-off of 0.5 and 0.9 nm, respectively. The spring constant of the harmonic restraints is set to 700 kJ mol^-1^ nm^-2^. In addition, the flag “-scfix” adds extra torsional potentials between backbone beads and sidechain beads^50^. These potentials are included in the force field to prevent unphysical flipping of side chains in β-strands.

It is important to note that the elastic network is created on the basis of a reference secondary structure that needs to be indicated to the marinize2 program, which can be done by adding the flag “-dssp mkdssp”. With this flag, the DSSP program (https://github.com/cmbi/dssp) is used to determine the secondary structure elements based on the input structure. DSSP recognizes secondary structure based on the positions of both heavy atoms and hydrogens, therefore, it is important to use a high-resolution initial structure or perform preprocessing, such as all-atom energy minimization, to generate accurate secondary-structure restraints.

To set up Martini simulations of mix-folded proteins, in which folded protein domains are linked by intrinsically disordered regions, it is recommended to follow the method and procedures introduced by the Lindorff-Larsen group^34^.

#### Representative Results of Option 7: SIRAH Coarse-Grained MD Simulations of A**β**42

Although SHRAH and Martini are quite different in various aspects, the general protocol of running MD simulations with these two force fields is actually the same when using Gromacs. The bash script ***setup_sirah.sh*** called with parameter ***Ab42*** (see point 7.2 in the protocol) performs the following operations:

- The initial structure of Aβ42, as given in the PDB file ***Ab42.pdb***, is protonated at ph=7.0 using the PDB2PQR software (with --with-ph=7.0 flag) and converted to a coarse-grained representation using the cgconv.pl program. The SIRAH topology and structure files are generated based on the information from the provided PDB file.
- The coarse-grained protein structure is placed in a cubic box of side length of 9 nm using the Gromacs editconf tool (with -box 9 9 9 flag) and solvated using the Gromacs solvate tool.
- Sodium and chloride ions are added into the simulation box in order to neutralize the system and to reach the physiological salt concentration of 0.15 M. Note that the flags “-np 66 -pname NaW -nn 63 - nname ClW” define the numbers of ions in the system. Users must manually recalculate the numbers of ions if they want to use a different ionic strength in their simulations.

The bash script ***run_min_eq.sh*** called with parameter ***Ab42*** (see point 7.3 in the protocol) performs energy minimization and equilibration simulations. The energy minimization is performed in two subsequent steps with parameters given in files ***md_sim_parameters/em1_CGPROT.mdp*** and ***md_sim_parameters/em2_CGPROT.mdp***. The equilibration simulations are conducted at constant volume (NVT ensemble) in two subsequent steps, respectively, with hard and soft restraints on the coordinates of the backbone beads. The parameters of these equilibration simulations are specified in the Gromacs input files ***md_sim_parameters/eq1_CGPROT.mdp*** and ***md_sim_parameters/eq2_CGPROT.mdp***. Analogously, the parameters of the production simulation are given in the input file ***md_CGPROT.mdp***.

The bash script ***run_md.sh*** called with parameter ***Ab42*** (see point 7.4 in the protocol) starts the production simulation at constant pressure (NPT ensemble). Next, when this simulation is completed, it centers the protein in the simulation box taking into account the periodic boundary conditions. The resulting trajectory file is named ***Ab42_cg_md_pbc.xtc*** and placed in the directory ***run_md***. The final configuration of the simulation system is saved in the ***Ab42_cg_md.gro*** file. With these two files, one can analyze the SIRAH simulation trajectory in the same way as the Martini simulation trajectories are analyzed here (see points from 5.5 to 5.8 in the protocol).

It should be noted that in order to perform SIRAH simulations of a different protein using Gromacs, users can simply copy a PDB file of their protein (e.g. ***myprotein.pdb***) to the ***GROMACS_SIRAH*** directory and perform steps from 7.1 to 7.5 of the protocol by successively executing the four scripts with the input parameter being the name of the PDB file (e.g. ***myprotein***).

#### Representative Results of Option 8: Use of CHARMM-GUI for MD Simulations of A**β**42

CHARMM-GUI stands as a versatile tool, facilitating a range of essential functions for manipulations of PDB files and generation of initial files for simulating various biological systems. Its capabilities span through the generation of lipid bilayers and micelles, addition of various ions, and more. For comprehensive tutorials and practical demonstrations, users can visit the official CHARMM-GUI website at https://www.charmm-gui.org/?doc=demo.

### Representative Results of the Simulations in Amber and Gromacs

Execution of the example scripts provided in the Supporting Materials generates a complete workflow for preparing input files for MD simulation, performing minimization, equilibration, and production runs, and conducting basic result analysis. Figure 1 illustrates differences between the all-atom and coarse-grained simulation boxes in the Amber and Gromacs packages. In Amber, a truncated octahedron periodic box is used to save computational time by omitting calculations for the water molecules placed in the corners of the box. In contrast, Gromacs coupled with the Martini 3 force field uses a more expensive but simpler cubic box in terms of symmetry operations and further analysis.

**Figure 1:**
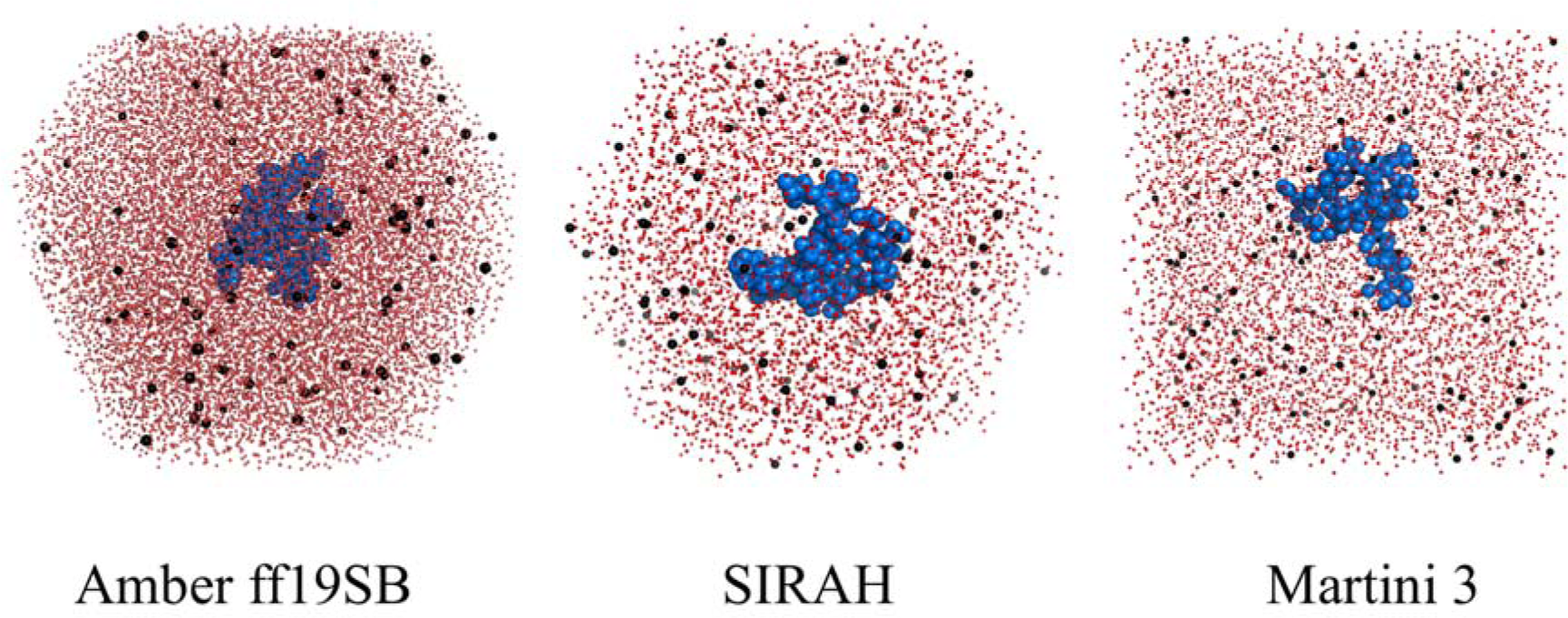
Simulation boxes. Visualization of the simulation boxes using both all-atom and coarse-grained methods. Atoms or interaction centers are represented as spheres in red (water), black (ions) and blue (Aβ42 peptide). To save computational time, the MD simulations in the Amber package employed a truncated octahedron box, while the Martini simulations used a standard cubic box.

Since the monomeric Aβ42 is intrinsically disordered and lacks a stable structure for reference in RMSD calculations, RMSD plots are not presented here. Nevertheless, RMSD can be easily calculated with minor modifications of the analysis scripts. The relevant properties that can be showcased here include the radius of gyration and the maximum dimension (Figure 2 for the results obtained using the Amber package). The presented results show that in the all-atom simulations with the ff19SB force field, Aβ42 tends to adopt a rather compact state with an average *R*_g_ of 1.29 ± 0.14 nm, and it becomes even more compact if the disulfide bond is present (Table 1). In the SIRAH coarse-grained model, Aβ42 exhibits a similar tendency, but the peptide conformations are even more compact, with an average *R*_g_ of 1.13 ± 0.11 nm without the disulfide bond present. To verify whether there are any contacts between periodic images (in other words, to check if the simulation box is sufficiently large), one can calculate the minimum-image distance (Figure 3). In our case, it is clear that there are no interactions between periodic images within the cut-off range (0.9 and 1.2 nm for ff19SB and SIRAH models, respectively), and the lowest observed values are above 1.6 and 2.6 nm for ff19SB and SIRAH models, respectively. These values are safely above the threshold, indicating that the periodic box was sufficiently large.

**Figure 2:**
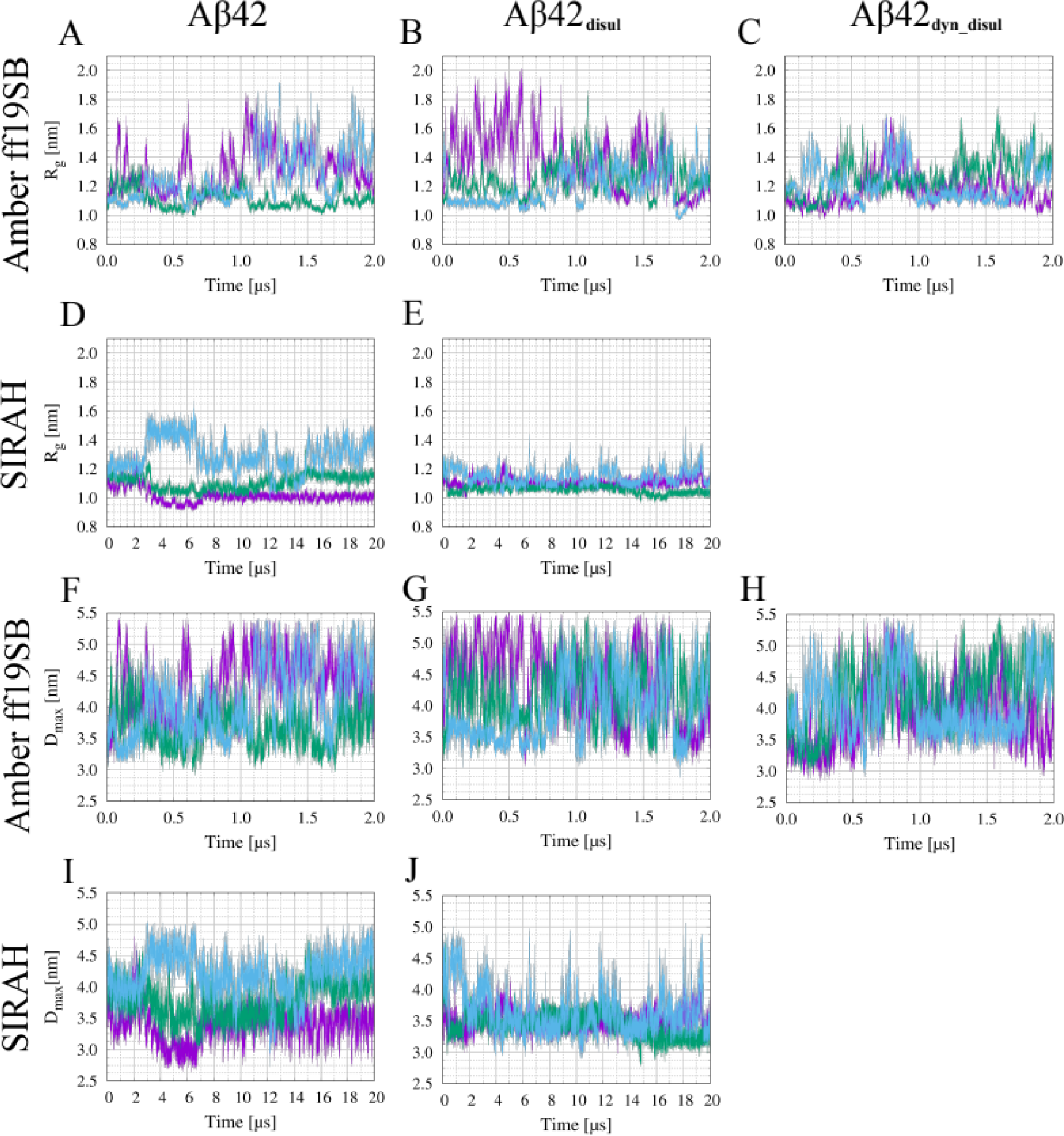
LJPlot of R_g_ and D_max_ in Amber and SIRAH. Time evolution of the radius of gyration (A-E) and of the maximum dimension (F-J) obtained from the MD simulations employing the all-atom Amber force field ff19SB (A-C and F-H) and SIRAH coarse-grained force field (DE and IJ) without the disulfide bond (A, D, F, I), with the static disulfide bond (B, E, G, J) and with the dynamic treatment of the disulfide bond (C and H).

**Figure 3:**
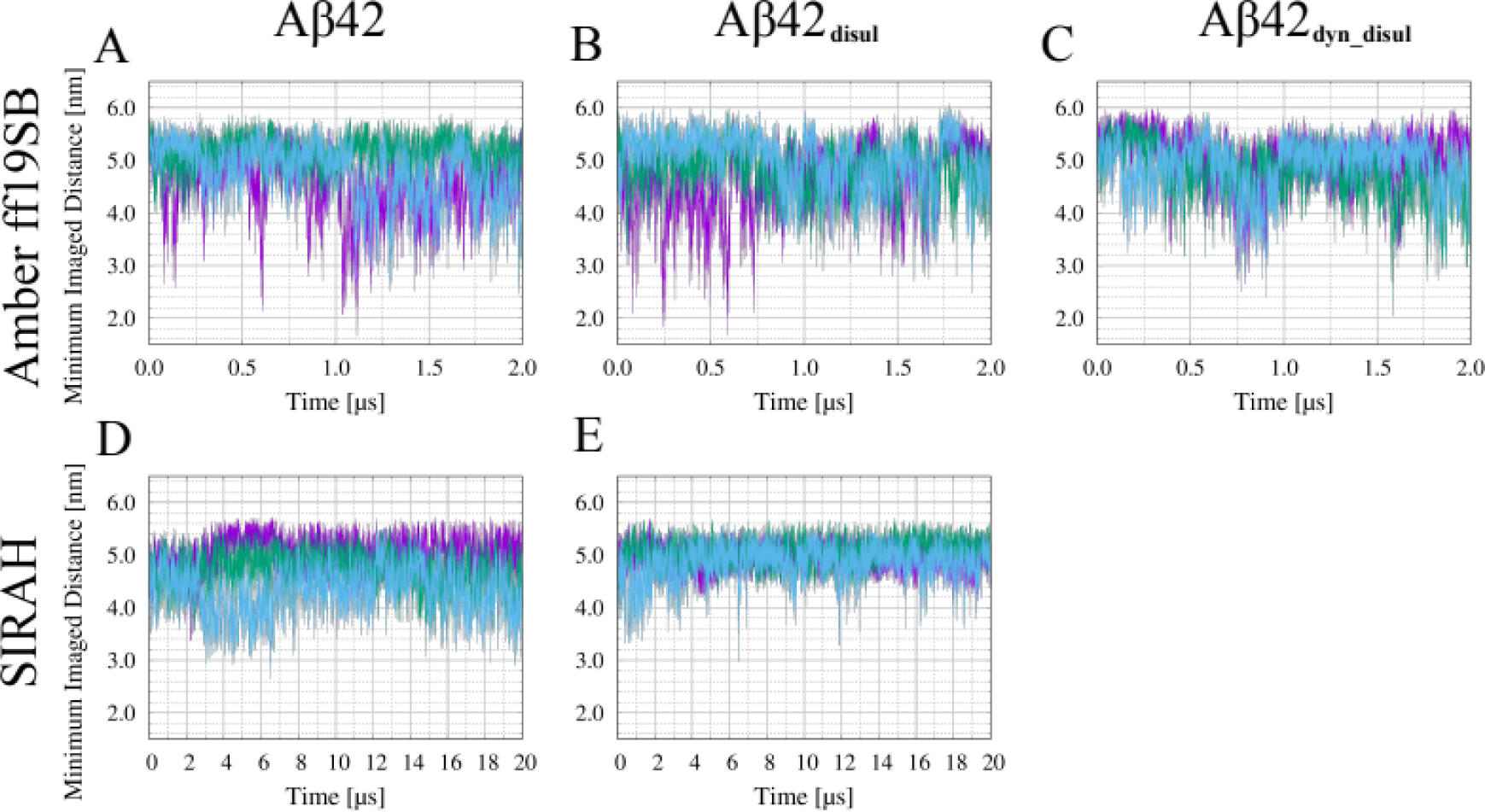
Periodic image distance. Time evolution of the minimum non-self image distance in the all-atom Amber ff19SB simulations of Aβ42 (A), of Aβ42 with the static disulfide bond (B), and of Aβ42 with the dynamic disulfide bond (C), as well as in the SIRAH coarse-grained simulations of Aβ42 (D) and of Aβ42 with the static disulfide bond (E).

**Table 1.** R_g_ and D_max_ values. Average R_g_ and D_max_ with SD (for the Amber and SIRAH force fields, the averages and SDs are calculated over three trajectories, while for Martini it is calculated over a single trajectory with block averaging).

| | Amber<br>ff19SB<br>Aβ42 | Amber<br>ff19SB<br>Aβ42<sub>disul</sub> | Amber<br>ff19SB<br>Aβ42<sub>dyn-disul</sub> | SIRAH<br>Aβ42 | SIRAH<br>Aβ42<sub>disul</sub> | Martini Aβ42<br>with $\lambda=1$ | Martini<br>Aβ42<sub>disul</sub> with<br>$\lambda=1$ |
| --- | --- | --- | --- | --- | --- | --- | --- |
| $R_g$ (nm) | 1.29±0.14 | 1.27±0.01 | 1.22±0.07 | 1.13±0.11 | 1.10±0.04 | 1.32±0.01 | 1.19±0.01 |
| $D_{max}$<br>(nm) | 4.22±0.42 | 4.16±0.06 | 4.09±0.21 | 3.83±0.31 | 3.48±0.09 | 4.20±0.04 | 3.83±0.03 |

Figure 4 shows results of the Martini simulations with λ=1, i.e. with no rescaling of the protein-water interactions. The radius of gyration as a function of time for Aβ42 and Aβ42_disul_ is shown, respectively, in panels A and B in Figure 4. Because of the constraint imposed by the covalent bond between Cys17 and Cys34, Aβ42_disul_ exhibits smaller *R*_g_ values than Aβ42. The abrupt fluctuations of *R*_g_ in time indicate that the simulation system is well equilibrated on the time scale of several microseconds. The maximum dimension as a function of time for Aβ42 and Aβ42_disul_ is shown, respectively, in panels C and D in Figure 4. It should be noted that *D*_max_ values do not exceed 7 nm, which is clearly smaller than the simulation box side length of about 9 nm. Thus, the simulation box is large enough to ensure that the peptide does not interact with its periodic image during the Martini simulations.

**Figure 4:**
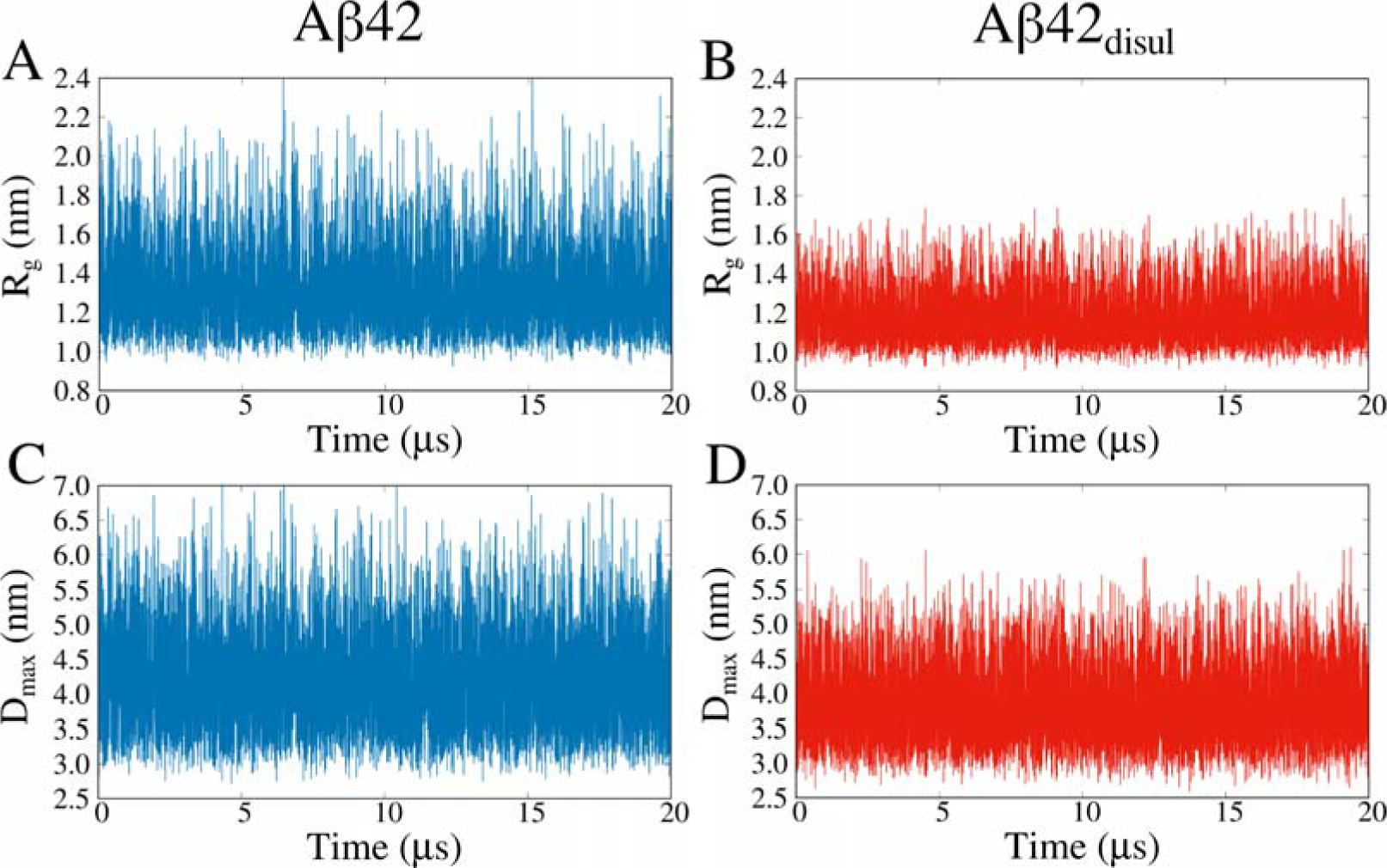
LJPlot of R_g_ and D_max_ in Martini. Time evolution of the radius of gyration (A,B) and of the maximum dimension (C,D) obtained from the Martini simulations of Aβ42 (A,C) and Aβ42_disul_ (B,D) with λ=1. Both Aβ42 and Aβ42_disul_ exhibit large conformational fluctuations at the thermodynamic equilibrium. Because of the constraint imposed by the covalent bond between Cys17 and Cys34, Aβ42_disul_ exhibits smaller values of *R*_g_ and *D*_max_ than Aβ42.

Figure 5 shows the average radius of gyration as a function of the water-protein interaction rescaling parameter λ used in the Martini simulations. As one would expect, the average *R*_g_ increases monotonically with λ, both in the case of Aβ42 and Aβ42_disul_. Indeed, increasing the protein-water interactions should result in expanded conformations of IDPs^34, 51^. Importantly, by comparing the average *R*_g_ from the simulations with the *R*_g_ from experimentals, one can determine an optimal value of parameter λ which yields the best agreement between the simulation and experiment^34^.

**Figure 5:**
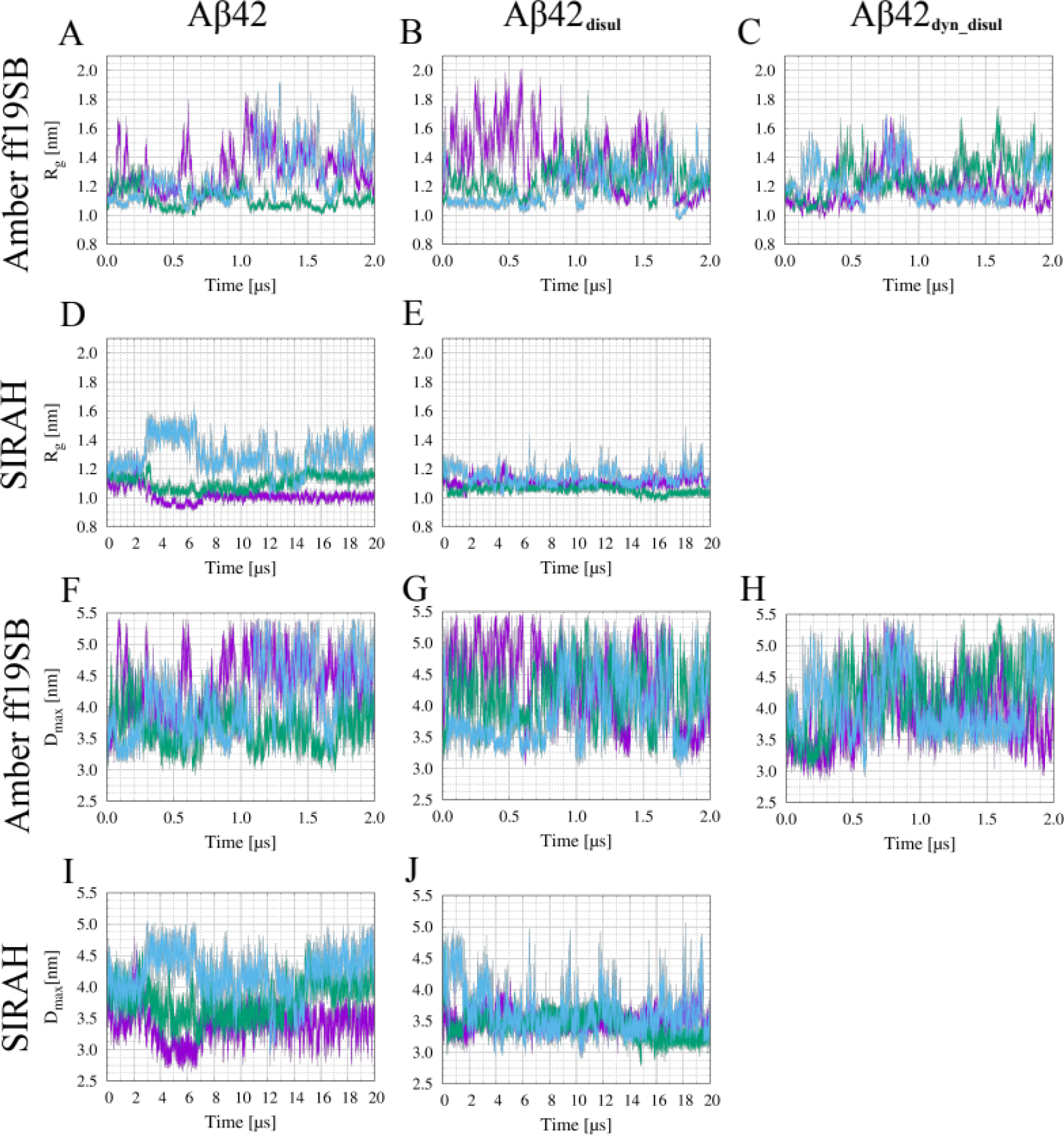
LJPlot of R_g_ vs. λ **in Martini.** Average radius of gyration as a function of the water-protein interaction rescaling parameter λ obtained from the Martini simulations of Aβ42 (blue) and Aβ42_disul_ (red). The error bars indicate the standard deviation calculated over a single trajectory using block averaging.

The contact maps of Aβ42 and Aβ42_disul_ obtained from the Martini trajectory with λ=1 are shown, respectively, in panels A and B in Figure 6. The color scale indicates the natural logarithm of contact frequency. Consequently, the points in red, yellow and blue represent frequent, transient and rare contacts, respectively.

**Figure 6:**
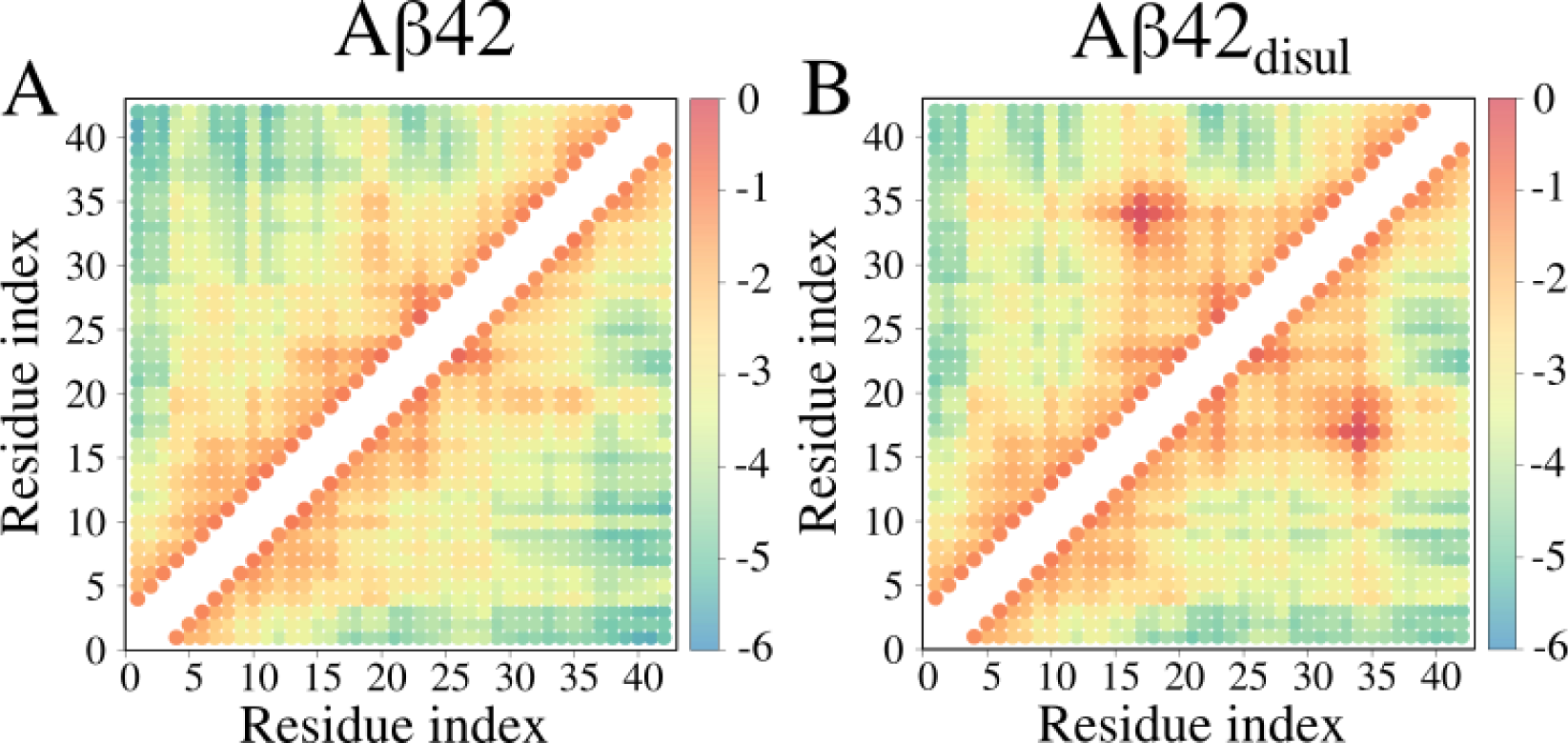
Contact map in Martini. Contact maps obtained from coarse-grained structures of Aβ42 (A) and Aβ42_disul_ (B) generated in the Martini simulations with λ=1 (no scaling of solute-solvent interactions). The color scale indicates the natural logarithm of contact frequency. The points in red, yellow and blue represent frequent, transient and rare contacts, respectively.

The contacts between amino acid residues that are close sequentially, namely contacts (i,i+1), (i,i+2), (i,i+3) and (i,i+4), are not shown. The presence of the disulfide bond between Cys17 and Cys34 is clearly reflected in the map shown in panel B. In general, contact map analysis can yield valuable insights into intramolecular interactions stabilizing protein conformations.

The result of mapping coarse-grained simulation structures back to the all-atom representation is illustrated in Figure 7. Specifically, the final structure obtained from the Martini simulation of Aβ42_disul_ with λ=1 is shown here in the van der Waals representation (panel A), in the stick-and-ball representation (panel B) and the ribbon representation (panel C). The all-atom structures obtained from the backmapping calculations can be used not only for visualization purposes but also for analysis and comparison with experimental data^34^.

**Figure 7:**
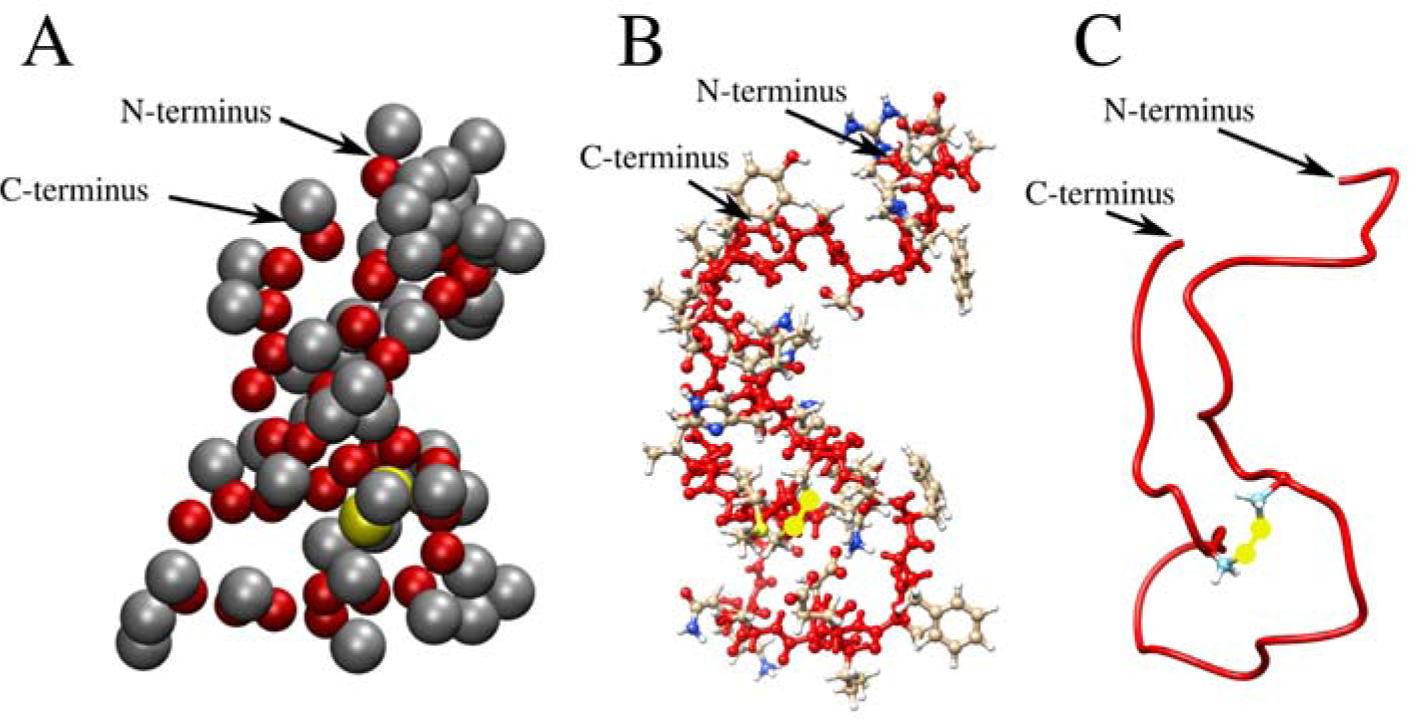
Representative structures. The final structure obtained from the Martini simulation of Aβ42_disul_ is shown in three different representations. (A) The coarse-grained structure, where the backbone and sidechain beads are colored in red and gray, respectively. The SC1 beads of Cys17 and Cys34 are highlighted in yellow. (B) The all-atom structure, as obtained from the backmapping calculation, is displayed in a stick-and-ball representation. The backbone atoms are shown in red. The sidechain carbon, nitrogen, and oxygen atoms are shown in gray, blue, and red, respectively. The sulfur atoms forming the covalent disulfide bridge are highlighted in yellow. (C) The all-atom structure is presented in a ribbon representation: the main chain is shown in red, while sidechains are not displayed, except for Cys17 and Cys34, which are shown in the stick representation.

Figure 8 compares the most extended and most compact structures obtained from the all-atom and coarse-grained simulations. Structures with and without the disulfide bond are depicted.

**Figure 8:**
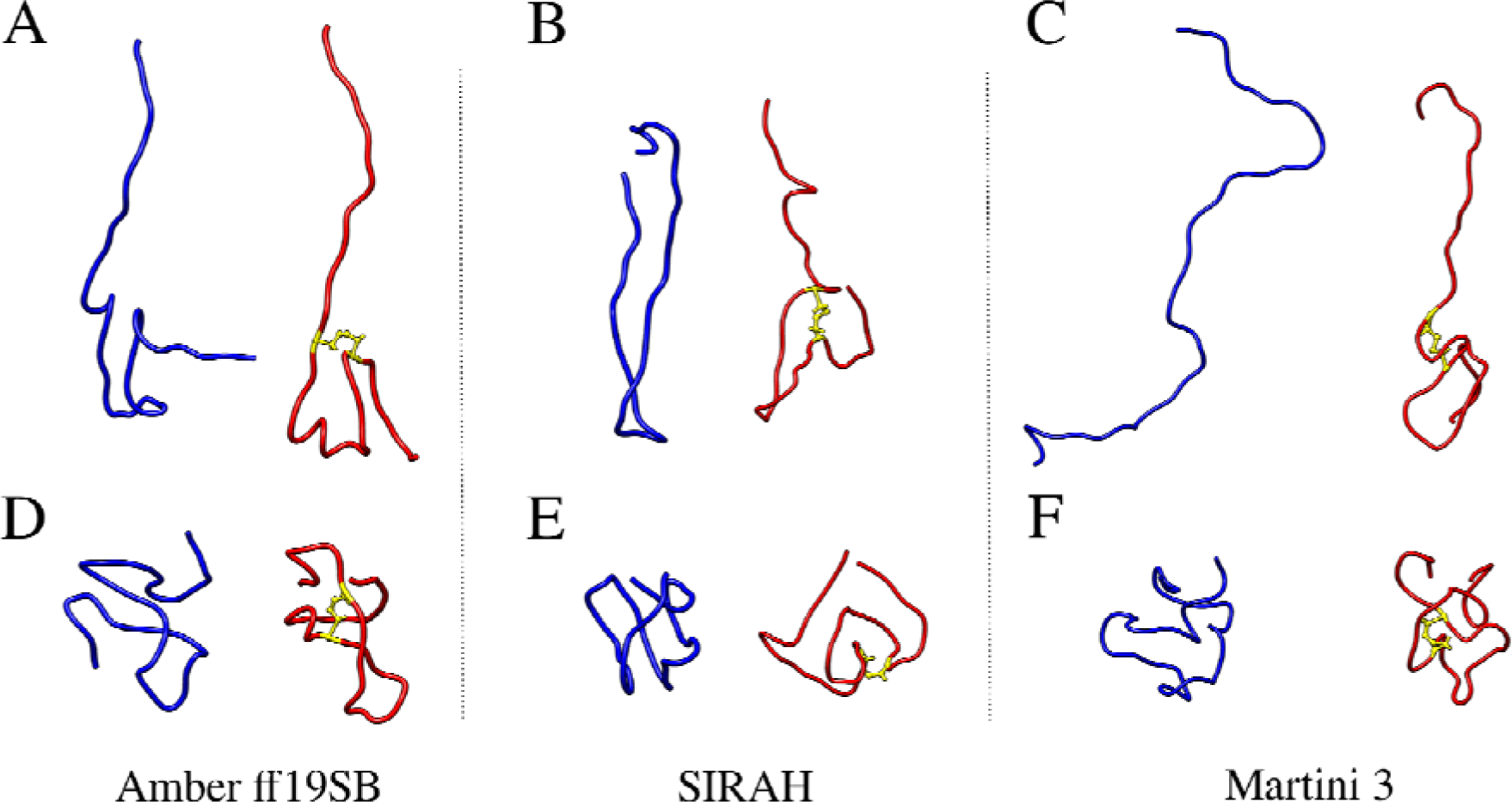
Most and least compact structures. Ribbon representation of the structures from the MD simulations with the all-atom Amber ff19SB (A, D), coarse-grained SIRAH (B, E) and coarse-grained Martini 3 (C, F) force fields with the maximum (A, B, C) and minimum (D, E, F) values of the radius of gyration in blue and red, respectively.

## DISCUSSION

In this work, we demonstrated how all-atom and coarse-grained force fields can be utilized to study a model IDP - monomeric Aβ42. To ensure the high reliability of the results, state-of-the-art methods were employed. Specifically, the all-atom Amber ff19SB force field for proteins was used, coupled with the four-point OPC water model^52^ and a co-optimized ions model^23^ to mimic physiological salt concentration, approximately 0.15M NaCl^53^. Additionally, the SIRAH coarse-grained force field within the Amber and Gromacs package and the Martini 3 coarse-grained model and force field within the Gromacs package were employed. In the latter approach, solute-solvent interactions were scaled up and down, and the results were compared to the radius of gyration values to ensure the correct compactness of the simulation structures. The use of a coarse-grained representation significantly accelerated the simulations by approximately 1-2 orders of magnitude, with no significant impact on the results. However, it is worth noting that the use of coarse-grained models can complicate the analysis, as it often requires reconstruction to all-atom models, and some atomistic details are lost due to simplifications of the protein representation. Still, coarse-grained models seem to be a method of choice when it comes to extensive simulations of large IDP complexes ^54^ and biomolecular condensates ^55, 56^.

Overall, the results obtained indicate that all the methods produced largely unstructured conformations, as expected for the monomeric Aβ42. However, this outcome is not always achieved, especially when older methods are employed ^13, 57^. Due to the much larger computational cost associated with the all-atom simulations, only three trajectories, each of 2 microseconds, were executed, which may not provide the highest level of robustness in the results. In contrast, with the coarse-grained SIRAH and MARTINI methods, simulations were 10 times longer, and convergence was reached more quickly due to the reduction in the number of degrees of freedom, resulting in a lower total computational cost. This approach of employing two entirely different computational methods, such as all-atom and coarse-grained force fields, is recommended for obtaining highly reliable results, especially in the absence of or with limited experimental data.

Earlier computational studies suggest that *R*_g_ values for Aβ42 are in range of 0.8 to 1.2nm^58–60^, while the experimental lowest found *R*_g_ values are about 1nm, which should correspond to the most compact conformations of the monomers. However, it should be noted that experimentally observed hydrodynamic radius strongly depends on the conditions and is found to be in the range of 0.8 to 1.5nm^60^, further confirming the molecular flexibility of Aβ42. More recent studies still present very different *R*_g_ values, ranging from 1.0^61^ to 1.6 nm^62^ obtained from SANS experiments. In the context of this evidence, all of the calculated values in this work fit into experimental and theoretical *R*_g_ ranges.

Point mutations are commonly introduced to proteins and peptides to either decrease^63^ or increase^9^ their aggregation properties, depending on the study’s objectives. In the case of Aβ42, a series of MD simulations were conducted on the L17C/L34C mutant with a disulfide bond acting as a cross-link between the amino-acid residues that come into contact in the fibrillar form. The designed mutant is known to efficiently form fibrils with both the wild-type and mutated Aβ42^9^, emphasizing some of the fibril-like contacts, including Q15-V36, L17-L34, and F19-I32. Interestingly, our all-atom and coarse-grained simulations, even without the disulfide bond, exhibit a significant number of conformations forming latter two contacts, and these conformations were further stabilized when a disulfide bond was incorporated (see Figure 6 and Table 2).

**Table 2.**
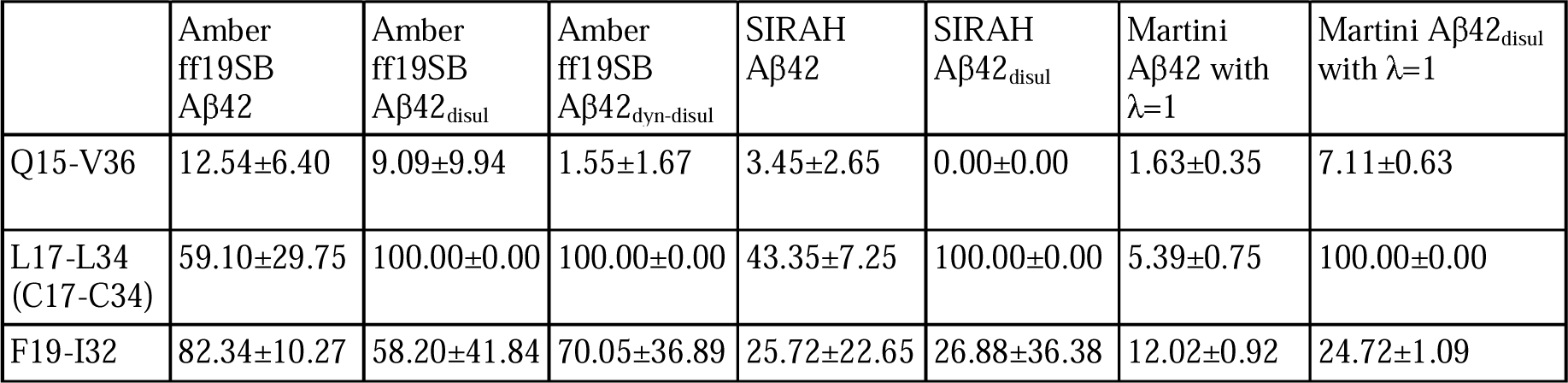
Frequency (in %) of contacts with SD in the simulations. A contact between atoms in a pair is counted as ‘1’ in a given snapshot if the CB atoms in all-atom representation or centers of interactions are within a 0.7 nm range.

The use of the coarse-grained SIRAH model enabled to obtain approximately a 6-fold decrease in the computational time needed to perform 1,000,000 steps compared to the all-atom ff19SB force field (Table 3). Taking into account a 10 times longer timestep value, this provided a 60-fold speed-up when using a low-end GPU, while the total number of interaction centers decreased by about 13 times. These values are in agreement with those obtained by SIRAH developers for a much larger system^36^. However, it should be noted that in the case of the monomeric Aβ4, the SIRAH simulations on a GPU did not scale at all, even with the use of a much more powerful GPU, while for the all-atom system, a 4-fold speed-up was observed, which most likely was caused by the small size of the studied system. An 80-fold speed-up was observed in the Amber all-atom force field when using a state-of-the-art GPU instead of a 20-core CPU node, whereas this difference was only about 10-fold if SIRAH is used in the Amber package. Interestingly, the SIRAH simulations in Gromacs are significantly faster then the SIRAH simulations in Amber when the CPU is used. This difference may be, at least partially, explained by the use of a simpler cutoff for the Van der Waals interactions in Gromacs (simple cutoff) than in Amber (PME). The dissimilarity disappears when the same GPU is used with PME treatment for all long-range interactions. While this observation may suggest that the CPU performance of Gromacs is superior to that of Amber, further investigations are needed to fully assess the impact of a simpler cutoff approach and to examine various systems on different hardware. It should be noted that use of Martini 3 model in the Gromacs package provides about a 15-fold speed-up compared to SIRAH in the Amber package when a similar CPU is used, which most likely comes from a much different coarse-grained model used (especially for water), different cutoff methods and ranges. Moreover, the performance of Martini in Gromacs scales well even when a high number of CPU cores is used (e.g. 3 times speed-up when 64 cores are used instead of 12). It should be noted that the Gromacs package^37^ is versatile in the sense that it can be used to perform MD simulations not only with such coarse-grained models as Martini^33^ and SIRAH^36^, but also within the all-atom model with various force fields, including GROMOS^64^, CHARMM^65^ and AMBER^20^. Similarly, the Amber package can be used with other than Amber force fields, including CHARMM for proteins and lipids, and GAFF^66^ for small organic molecules.

**Table 3.**
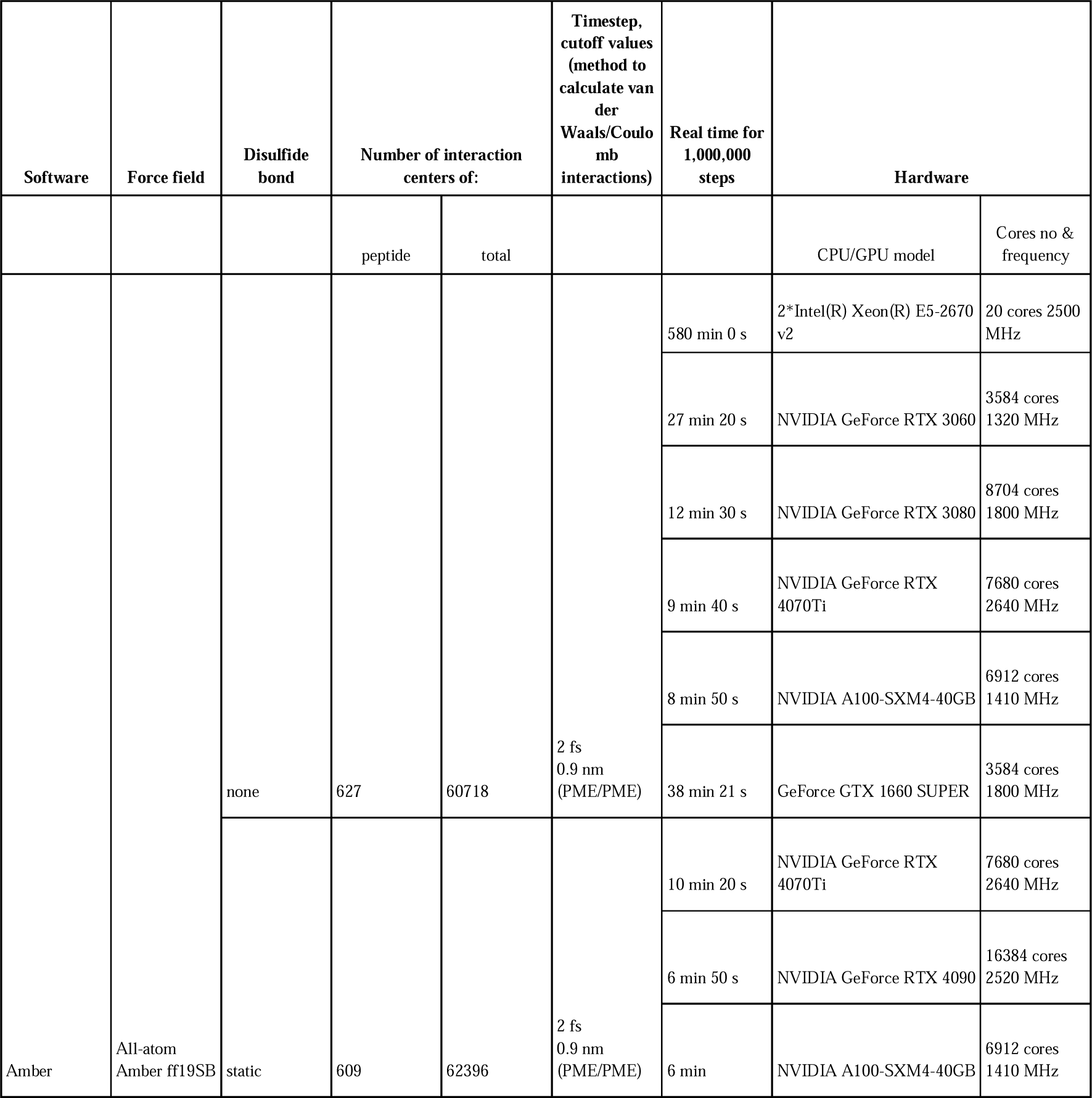
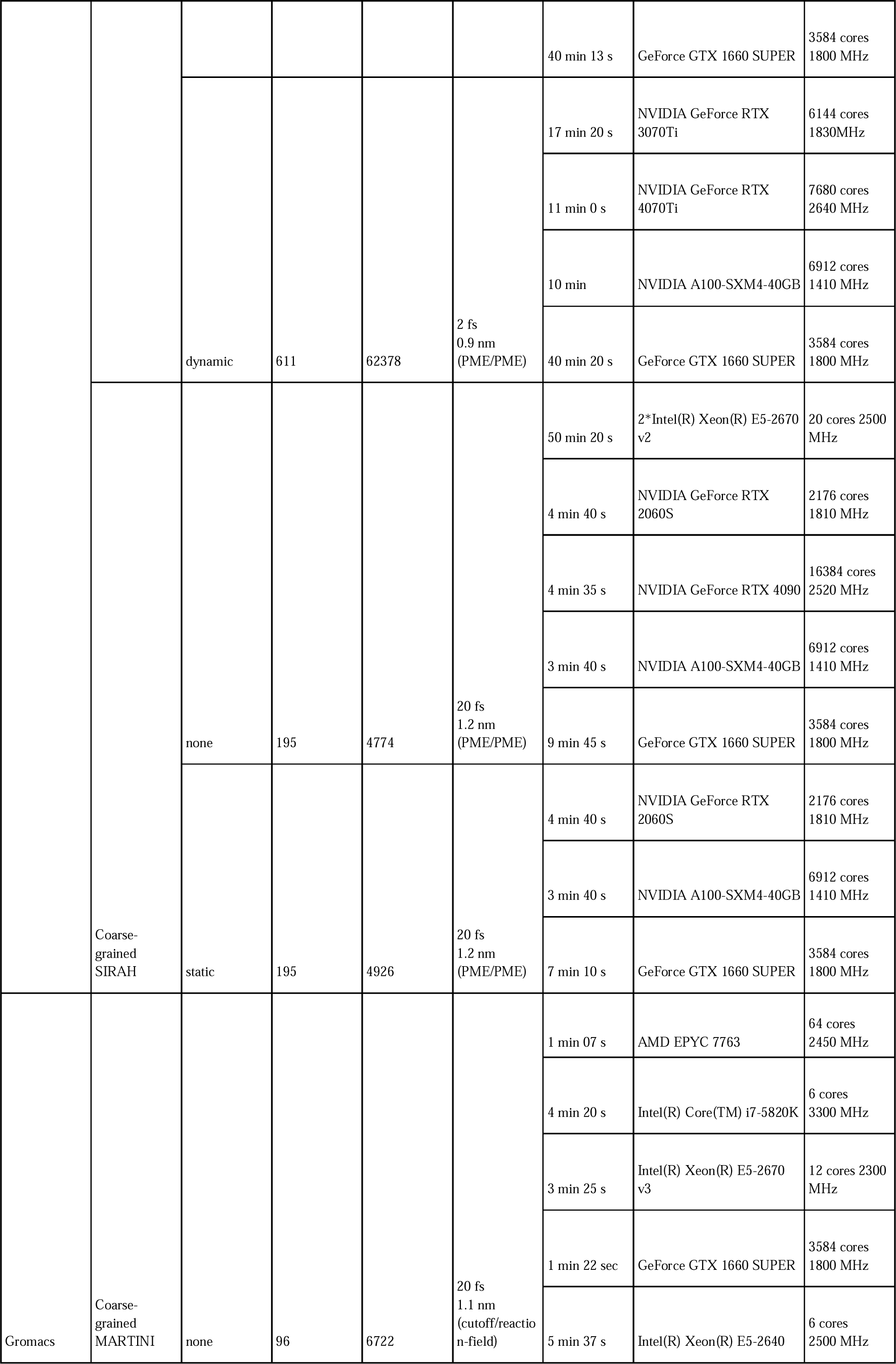
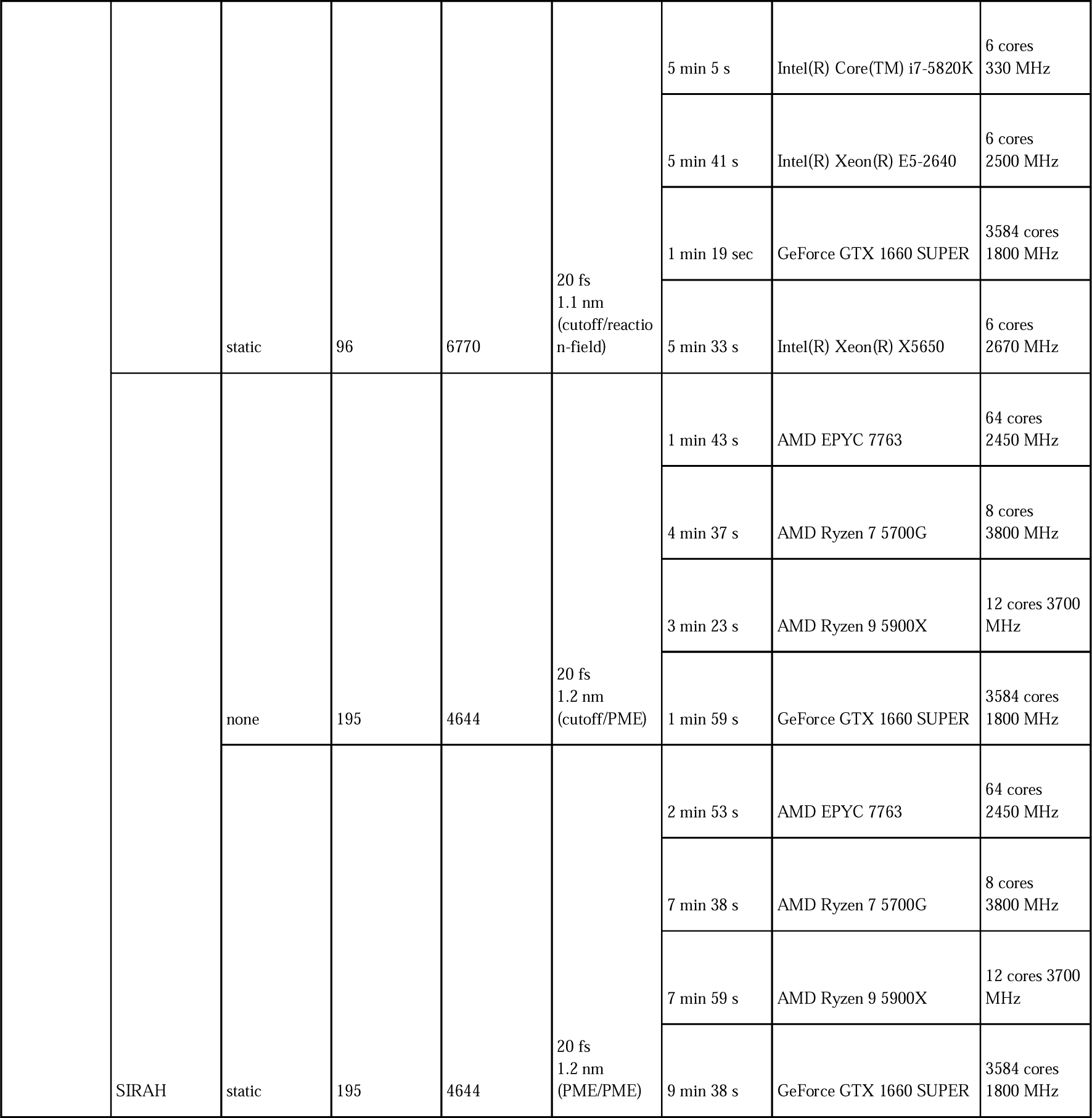
CPU and GPU performance. Computational times for various all-atom and coarse-grained approaches for the monomeric Aβ42 with and without disulfide bond treatments are shown with multiple configurations to demonstrate that these simulations can be run on a modern PC or a computer cluster.

In general, the provided protocols can be easily employed for routine simulations of most biomacromolecules, with the flexibility to change the force field, water model, and other components of the system. This adaptability can be managed even by less experienced users. While it is recommended to conduct classical MD simulations on supercomputers, access to which is often limited, the use of GPU computations and coarse-grained representations allows these simulations to be successfully run on modern desktop computers, which are usually equipped with up to 32-core CPUs. This democratizes access to scientific research by eliminating the need for substantial budgets, thereby promoting inclusivity in scientific studies.

## Supporting information

Supporting file - scripts

## ACKNOWLEDGMENTS

This work has received financial support from the National Science Centre, Poland, *via* grants SONATA No 2019/35/D/ST4/03156 (to PK) and OPUS No 2020/39/B/NZ1/00377 (to BR). PK and PS gratefully acknowledge Polish high-performance computing infrastructure PLGrid (HPC Centers: ACK Cyfronet AGH) for providing computer facilities and support within computational grant no. PLG/2023/016624. BR thanks the Centre of Informatics – Tricity Academic Supercomputer and networK (CI TASK) in Gdansk, Poland, for access to computer resources. PR, BR and MMA acknowledge Polish high-performance computing infrastructure PLGrid for awarding this project access to the LUMI supercomputer, owned by the EuroHPC Joint Undertaking, hosted by CSC (Finland) and the LUMI consortium through PLL/2023/04/016485.

## DISCLOSURES

Authors declare that there are no competing interests.

## Notes

### Competing Interest Statement

The authors have declared no competing interest.

### Summary of Updates

Click-and-go protocol was added in addition to the previous description, as well as, example how to perform SIRAH calculations in Gromacs and use of CHARMM-GUI is added making protocols more robust and versatile. Other minor improvements were also added.

