## Supporting file - scripts for "Protocols for Multi-Scale Molecular Dynamics Simulations: A Comparative Study for Intrinsically Disordered Amyloid Beta in Amber & Gromacs on CPU & GPU": setup_amber.sh

```
#!/usr/bin/bash

#Enter the directory and execute system preparation
cd tleap
echo "Running preparation in directory:"
pwd
tleap -f tleap.in
#Enter the directory and run minimalization
cd ../min
echo "Running minimalization in directory:"
pwd
bash start_min.sh
#Enter the directory and run equilibration on CPU
cd ../eq1
echo "Running equilibration on CPU in directory:"
pwd
bash start_eq1.sh
#Enter the directory and run equilibration on GPU
cd ../eq2
echo "Running equilibration on GPU in directory:"
pwd
bash start_eq2.sh
cd ../
echo "System setup, minimization and equilibration finished"

echo "Start production for first trajectory in directory /MD1/dyn1"

#cd MD1/dyn1
#pwd
#bash start_dyn1.sh
#cd ../../
```
