## Supporting file - scripts for "Protocols for Multi-Scale Molecular Dynamics Simulations: A Comparative Study for Intrinsically Disordered Amyloid Beta in Amber & Gromacs on CPU & GPU": run_analysis.sh

```
#!/usr/bin/bash
dir=`pwd`
echo $dir

mkdir rg  rmsd  rmsf pdb mindist dmax

cd ..

#names of different trajectories
for md in MD1 # MD2 MD3 
do

#names of simulations	
for file in amber amber_disul amber_dyn_disul 

do
cd $file
mkdir ptraj_${md}
cd ptraj_${md}

echo ${file}_${md}

cp $dir/ptraj_traj_${md}.in ./ptraj_traj.in
cp $dir/ptraj_rmsd_rg.in ./
cp $dir/ptraj_pdb.in ./
cp $dir/ptraj_image_distance.in ./


cpptraj -i ptraj_traj.in 
cpptraj -i ptraj_rmsd_rg.in 
cpptraj -i ptraj_pdb.in
cpptraj -i ptraj_image_distance.in

cp rg.txt ${dir}/rg/${file}_${md}_rg.txt
cp rmsf.txt ${dir}/rmsf/${file}_${md}_rmsf.txt
cp rmsd.txt ${dir}/rmsd/${file}_${md}_rmsd.txt
mv traj.pdb ${dir}/pdb/${file}_${md}.pdb
cp mindist.txt ${dir}/mindist/${file}_${md}_mindist.txt 
cp dmax.txt ${dir}/dmax/${file}_${md}_dmax.txt

cd ../../
done
done
```
