## Supporting file - scripts for "Protocols for Multi-Scale Molecular Dynamics Simulations: A Comparative Study for Intrinsically Disordered Amyloid Beta in Amber & Gromacs on CPU & GPU": run_analysis_rg_snap.sh

```
#!/usr/bin/bash
dir=`pwd`
echo $dir

rm rg_minmax.txt 
echo "title snap-max rg-max snap-min rg-min" >>  rg_minmax.txt
mkdir rg_snaps

cd ..

for file in amber amber_disul amber_dyn_disul 
do

for md in MD1 MD2 MD3
do

cd $file
mkdir ptraj_${md}
cd ptraj_${md}

rg_min=`cat rg.txt |grep -v Frame| sort -r -nk2,2| tail -1|awk '{print $2}'`
snap_min=`cat rg.txt |grep -v Frame| sort -r -nk2,2| tail -1|awk '{print $1}'`

rg_max=`cat rg.txt |grep -v Frame| sort -r -nk2,2| head -1|awk '{print $2}'`
snap_max=`cat rg.txt |grep -v Frame| sort -r -nk2,2| head -1|awk '{print $1}'`

echo ${file}_${md} ${snap_max} ${rg_max} ${snap_min} ${rg_min} >> ${dir}/rg_minmax.txt

cp $dir/ptraj_pdb_snap.in ./

sed -i "/s/MAX/${snap_max}/" ptraj_pdb_snap.in
sed -i "/s/MIN/${snap_min}/" ptraj_pdb_snap.in

mv traj_rg_min.pdb ${dir}/rg_snaps/${file}_${md}_rg_min.pdb 
mv traj_rg_max.pdb ${dir}/rg_snaps/${file}_${md}_rg_max.pdb

cd ../../
done
done
```
