## Supporting file - scripts for "Protocols for Multi-Scale Molecular Dynamics Simulations: A Comparative Study for Intrinsically Disordered Amyloid Beta in Amber & Gromacs on CPU & GPU": start_dyn1.sh

```
#!/bin/bash
export CUDA_VISIBLE_DEVICES=0

cp ../../eq2/*_eq2.crd ./
cp ../../min/*.top ./

pmemd.cuda -O -i md_smd.in -c system_eq2.crd -p system.top -o min.out -r system_smd.crd -x traj.nc 

ambpdb -p system.top -c system_smd.crd > output_smd.pdb
```
