## Supporting file - scripts for "Protocols for Multi-Scale Molecular Dynamics Simulations: A Comparative Study for Intrinsically Disordered Amyloid Beta in Amber & Gromacs on CPU & GPU": start_eq2.sh

```
#!/bin/bash

#export CUDA_VISIBLE_DEVICES=0

#echo "Change paths in file as apropriate"
#exit 0
#hash above after checking next two lines
#export CUDA_HOME=/usr/local/cuda-11.8
#source /home/../amber22/amber.sh

cp ../eq1/*_eq1.crd ./
cp ../eq1/*.top ./

pmemd.cuda -O -i md_eq2.in -c system_eq1.crd -p system.top -o min.out -r system_eq2.crd -x traj.nc
```
