## Supporting file - scripts for "Protocols for Multi-Scale Molecular Dynamics Simulations: A Comparative Study for Intrinsically Disordered Amyloid Beta in Amber & Gromacs on CPU & GPU": README_AMBER.txt

### Simulation Instructions

1. Before running the simulation, ensure that your login resource file (e.g., `~/.bashrc`, `~/.zshrc`, etc.) contains the appropriate sourcing:

    export CUDA_HOME=/usr/local/cuda-X.X # Replace X.X with the correct CUDA version
    source /home/xxxx/amber22/amber.sh

    Make sure to replace `X.X` with the specific CUDA version you are using and `xxxx` with the appropriate file path.

2. Verify if CUDA is functioning correctly by entering the following command in the terminal:

    nvidia-smi

    This command will display information about your NVIDIA GPU(s) and their current usage.

3. For the Steered Molecular Dynamics (SMD) process aimed at bringing sulfur atoms from cysteines close to each other, modify or verify the parameters in the `dist.RST` file:

    - Ensure the following are accurate in the `dist.RST` file:
    - The distance between sulfur atoms (located after the `r2=` flag)
    - The indices or numbers identifying the specific sulfur atoms (found after the `iat=` flag)

    Verify and/or adjust these values as necessary to accurately represent the intended simulation configuration for the SMD process.

    It's essential to double-check the `dist.RST` file to accurately define the distance and atom indices before initiating the SMD simulation.
