## Supporting file - scripts for "Protocols for Multi-Scale Molecular Dynamics Simulations: A Comparative Study for Intrinsically Disordered Amyloid Beta in Amber & Gromacs on CPU & GPU": 0README

#----------------------------------------------------------------------#
# SIRAH Tools                                                          #
#----------------------------------------------------------------------#
# A set of utilities for mapping, backmapping and visualizing          #
# coarse-grained models distributed within the SIRAH package (c)       #
#                                                                      #
# AUTHOR:    Matias Machado                                            #
#                                                                      #
# REFERENCE: Machado M, Pantano S. Bioinformatics (2016)               #
#            DOI: 10.1093/bioinformatics/btw020                        #
#                                                                      #
# This program is free software; you can redistribute it and/or modify #
# it under the terms of the GNU General Public License as published by #
# the Free Software Foundation; either version 2 of the License, or    #
# (at your option) any later version.                                  #
#                                                                      #
# This program is distributed in the hope that it will be useful,      #
# but WITHOUT ANY WARRANTY; without even the implied warranty of       #
# MERCHANTABILITY or FITNESS FOR A PARTICULAR PURPOSE. See the         #
# GNU General Public License for more details.                         #
#----------------------------------------------------------------------#
