## Supporting file - scripts for "Protocols for Multi-Scale Molecular Dynamics Simulations: A Comparative Study for Intrinsically Disordered Amyloid Beta in Amber & Gromacs on CPU & GPU": cgconv.pl

```
#!/usr/bin/perl

#----------------------------------------------------------------------#
# CGCONV                                                               #
#----------------------------------------------------------------------#
# Program to convert all atom structures in PDB format to coarse       #
# grained models.                                                      #
#                                                                      #
# USAGE:   cgconv.pl [-i] [-o] [-R] [-aPLhv] [MAP]... <STDIN> STDOUT   #
#                                                                      #
# AUTHOR:  Matias Machado                                              #
# E-MAIL:                                      #
# VERSION: 15, April 2020                                              #
#                                                                      #
# This program is free software; you can redistribute it and/or modify #
# it under the terms of the GNU General Public License as published by #
# the Free Software Foundation; either version 2 of the License, or    #
# (at your option) any later version.                                  #
#                                                                      #
# This program is distributed in the hope that it will be useful,      #
# but WITHOUT ANY WARRANTY; without even the implied warranty of       #
# MERCHANTABILITY or FITNESS FOR A PARTICULAR PURPOSE. See the         #
# GNU General Public License for more details.                         #
#----------------------------------------------------------------------#

  %opts=();
  use Getopt::Std;
  getopts("i:o:R:avhLP",\%opts);

############ Default variables ######################

  $INPUT  = "STDIN";
  $OUTPUT = "STDOUT";

# Script directory Path
  use File::Basename;
  $cgconv_path = dirname($0);

########## Program run starts here ##################

# Checking opts arguments
  if (  $opts{"v"} ) { print STDERR "version 15 [Apr 2020]\n"; exit 1 }
  if ( ($opts{"h"} == 1) || (!(exists $opts{"i"}) && (-t STDIN )) ) {&Usage;}

# Residue selection
  %selection=();

  if ( $opts{"R"} ) {

     while ( $opts{"R"} =~/([A-Za-z]*)\[([0-9][0-9,-]*)\]([A-Za-z]*)|([A-Za-z]+)|([0-9][0-9,-]*)/g){
            
           if ($1 | $3) {
                       @chain = split('',($1 . $3));

                       foreach $chain (@chain) {
                                push ( @{$selection{"$chain"}}, &GetResid($2) );
                         
                       }
           }

           elsif   ($4) {
                         @chain=split('',$4);
                        
                         foreach $chain (@chain) { $selection{"$chain"}=""; }

           }
             
           elsif  ($5 | $2) { push (@{$selection{'*'}}, &GetResid($5 . $2)); }

     }
   
     $opts{"P"} = 1; # Set flag -P on
  }

########## Load data from MAP file(s) ###############

# Set sirah_*.map files
  opendir(DIR, "$cgconv_path/maps/") or die "Can't open dir $cgconv_path/maps/ \n";

  while (my $file = readdir(DIR)) {

        if ($file =~ m/^sirah\_.+\.map$/) { push (@SIRAHTOP, "$cgconv_path/maps/$file"); }
  }

  closedir(DIR);

  if (!(@ARGV[0])) { # If no MAP file is set then use sirah_*.map files

        @CGTOP = @SIRAHTOP;
  }

# Else, use custom MAP file(s) or append them to sirah
  else { @CGTOP = @ARGV[0 .. $#ARGV]; if ($opts{"a"}) { @CGTOP = (@SIRAHTOP, @CGTOP) } }


# Open log file
  if ($opts{L} == 1) {open (LOG ,"> cgconv.log") or die "Can't open LOG file cgconv.log for writting\n";}

  foreach $cgtop (@CGTOP){

        # Data structure: hash %map_db
        #
        #                ELEMENT                  FUNCTION         DATA
        # ---------------------------------------------------------------------------------
        #
        # ( AAResName = (
        #                CGRES = CGResName,
        #
        #                ATOMS = (
        #                          AAAtomName = (
        #                                         MAP   =                     CGAtomName,
        #                                         GROUP = (
        #                                                  ndx = ( 
        #                                                          TYPE     = CGAtomName,
        #                                                          FUNCTION = CMASS/CGEOM,
        #                                                        )
        #                                                  ...
        #                                                 )
        #                                       )
        #                          ...
        #                        )
        #               )
        #   ...
        # )

          open (CGTOP ,"<$cgtop") or open (CGTOP ,"<$cgconv_path/maps/" . basename ($cgtop)) or die "Can't open MAP file $cgtop nor $cgconv_path/maps/" . basename($cgtop) . " for reading\n";
       
          if ($opts{L} == 1) {print LOG "Reading MAP file $cgtop\nChecking for syntaxis errors.\n";}
       
          $CGRES_ndx = 0;
       
          while (<CGTOP>) {
               
              # Ignore empty lines or lines starting with '#'
                if ( $_ =~ /^[\s]*[#]|^[\s]+$/ ) {next;}

                if ($_ =~ /^[\s]*[\>][\s]*CGRES[\s]*$/){
               
                   if (%CGRES) { 
                  
                      if ( !($CGRES{"CGNAME"}) || !($CGRES{"ALLNAME"}) || !($CGRES{"ATOMS"}) ) {&Error_2;}
                     
                      &AddMap(\%CGRES); %CGRES=();
                   }
                  
                   if ($opts{L} == 1) {++$CGRES_ndx; print LOG "\n>> CGRES $CGRES_ndx <<\n";}
                  
                   $gr_ndx=1;
                   next;
                }

                $_ =~ s/^\s+//; # remove leading whitespace
                $_ =~ s/\s+$//; # remove trailing whitespace return
                $_ =~ s/(.*)=>(.*)/\1 \2/;
               
                @data = split (/\s+/, $_);
               
              # Data structure
              # %CGRES = CGNAME  => CGResName,
              #          ALLNAME => [AAResName ...],
              #          ATOMS   => (AAAtomName => ( FUNCTION => data ))
               
                if    ($data[0] eq "CGNAME") { if ($#data != 1) {&Error_1;} $CGRES{"CGNAME"}  = $data[1]; }
                
                elsif ($data[0] eq "ALLNAME"){ if ($#data <  1) {&Error_1;} $CGRES{"ALLNAME"} = [ @data[1..$#data] ]; }
               
                elsif ($data[0] eq "MAP") { 
                
                      if ($#data < 2) {&Error_1;}
                      
                      for $attype (@data[1..($#data -1)]) { $CGRES{"ATOMS"}{"$attype"}{"MAP"} = $data[$#data]; } }
                
                elsif ($data[0] =~ /^(CMASS|CGEOM)$/) { 
                      
                      if ($#data < 2) {&Error_1;}
                      
                      for $attype (@data[1..($#data -1)]) {
                      
                          %{$CGRES{"ATOMS"}{"$attype"}{"GROUP"}{"$gr_ndx"}} = (
                                                                               "TYPE"     => $data[$#data],
                                                                               "FUNCTION" => $data[0]     ,
                                                                              );
                      }
                      
                      ++$gr_ndx;
                      
                      if ($data[0] eq "CMASS") {$usemass = 1;}
                }
                
                else {&Error_1;}
                
                if ($opts{L} ==  1) {print LOG "$data[0]....OK\n";}
          }
       
       # EOF
       if (%CGRES){ &AddMap(\%CGRES); $gr_ndx=1; %CGRES=(); }

  close CGTOP;                        
                         
  }


########## Load data from Atom Type MW file #########

# Load file only if it will be used
  if ($usemass) {&LoadAtMW;}

########## Load data from PDB file ##################

# Open input file
  if ($opts{i}) {$INPUT = "FILEIN"; open ($INPUT ,"<$opts{i}") or die "Can't open input file $opts{i} for reading\n";}

# Open output file
  if ($opts{o}) {$OUTPUT = "FILEOUT"; open ($OUTPUT ,">$opts{o}") or die "Can't open output file $opts{o} for writting\n";}

  my %no_map   = (); # resnames not found in map files
  my %res      = (); # group information
  my $resid    = ''; # residue number
  my $chain    = ''; # chain ID
  my $achar    = ''; # Code for insertion of residues
  my $atom_ndx =  0; # atom index

  while (<$INPUT>) {

        $_ =~ s/^\s+|\s+$//g; # remove leading and trailing whitespace & return
 
        if ( $_ =~ /^(TER|END)/ ) {
   
           if (%res) { &PrintGroups(\%res); %res = (); } # Print stored data

           print $OUTPUT "$1\n";
        
           $resid = '';
           $chain = '';
           $achar = '';
           next;
        } 
       
        elsif (!($_ =~ /^ATOM|^HETATM/)) {next;}
   
      # http://deposit.rcsb.org/adit/docs/pdb_atom_format.html
      # Allow (nonstandard) use of four-character residue names occupying an additional column (18-21)
   
      #           [0]       [1]      [2]   [3]        [4]   [5][6][7]
      # @atom = ( atom_type res_name chain res_number achar  x  y  z )
      #                  1               2                3            4           5          6         7         8
      #         123456789012 3456  7   8901  2  3456  7 890 12345678  90123456  78901234 56789012345678901234567890 
        @atom = /^............(....)[A ](....)(.)(....)(.)...(........)(........)(........).*/ or next;

        for (@atom) {s/^\s{0,}([^\s]+)\s{0,}$/\1/g;} # Remove leading and trailing whitespaces from array's elements

        if ( (%res) && (($resid != $atom[3]) || ($chain ne $atom[2]) || ($achar ne $atom[4])) ) { # If next residue and exists stored data

           &PrintGroups(\%res); %res = ();
           $resid = $atom[3];
           $chain = $atom[2];
           $achar = $atom[4];
        }
   
      # Apply filter over residue selection
        if ( !(&InSelection($atom[2], $atom[3], \%selection)) ) {
      
           if ( $opts{"P"} == 1 ) {
           
              ++$atom_ndx; if ($atom_ndx > 99999) { $atom_ndx = 0; } ; # Reset counter
              printf $OUTPUT ("ATOM%7d",$atom_ndx); print $OUTPUT substr($_,11,44),"\n";
           }

           next;
        }
      # ------------------------------------
   
      # Evaluate mapping
   
      # No residue mapping
        if ( !(exists $map_db{"$atom[1]"}) ){
   
           if ( $opts{"P"} == 1 ) {
   
              ++$atom_ndx; if ($atom_ndx > 99999) { $atom_ndx = 0; } ; # Reset counter
              printf $OUTPUT ("ATOM%7d",$atom_ndx); print $OUTPUT substr($_,11,44),"\n";
           }
        
           $no_map{"$atom[1]"} = 1; # hash list of not mapped residues
           next;
        }
   
      # Evaluate mapping on current atom
      # The following code is compatible with old perl versions
        @function = keys %{$map_db{"$atom[1]"}{"ATOMS"}{"$atom[0]"}};

      # SKIP
        if (!@function) {delete $map_db{"$atom[1]"}{"ATOMS"}{"$atom[0]"}; next} # To avoid 'autovivification' of elements

      # MAP
        if ( $function[0] eq "MAP" || $function[1] eq "MAP" ) {
   
           ++$atom_ndx; if ($atom_ndx > 99999) { $atom_ndx = 0; } ; # Reset counter
           printf $OUTPUT ("ATOM%7d%5s %-4s",
                            $atom_ndx,
                            $map_db{"$atom[1]"}{"ATOMS"}{"$atom[0]"}{"MAP"},
                            $map_db{"$atom[1]"}{"CGRES"}
                          );
           
           print  $OUTPUT (substr($_, 21,34),"\n");
        }
   
      # CMASS/CGEOM
        if ( $function[0] eq "GROUP" || $function[1] eq "GROUP" ) {

              for $gr_ndx (keys %{$map_db{"$atom[1]"}{"ATOMS"}{"$atom[0]"}{"GROUP"}}) {
               
                  $mass  = $atomtypes{"$atom[0]"} ;
                  $apply = $map_db{"$atom[1]"}{"ATOMS"}{"$atom[0]"}{"GROUP"}{"$gr_ndx"}{"FUNCTION"}; # CMASS/CGEOM

                # Atom mass and total mass of the group
                  if ($apply eq "CGEOM") {$mass = 1;} elsif (!($mass)) {&Warning_1; $mass = 0;}

                  $res{"$gr_ndx"}{"mass"}   += $mass;
               
                # Compute and store center of mass/geom
                  $res{"$gr_ndx"}{"x"}      += ( $atom[5] * $mass );
                  $res{"$gr_ndx"}{"y"}      += ( $atom[6] * $mass );
                  $res{"$gr_ndx"}{"z"}      += ( $atom[7] * $mass );

                # Store Extra Data
                  $res{"$gr_ndx"}{"cgres"}   = $map_db{"$atom[1]"}{"CGRES"};
                  $res{"$gr_ndx"}{"cgattyp"} = $map_db{"$atom[1]"}{"ATOMS"}{"$atom[0]"}{"GROUP"}{"$gr_ndx"}{"TYPE"};
                  $res{"$gr_ndx"}{"chain"}   = $atom[2]; $chain = $atom[2];
                  $res{"$gr_ndx"}{"resid"}   = $atom[3]; $resid = $atom[3];
                  $res{"$gr_ndx"}{"achar"}   = $atom[4]; $achar = $atom[4];
              }
        }
  }

# EOF
  if (%res) {&PrintGroups(\%res); %res = ();}

  close $INPUT;
  close $OUTPUT;

  &Warning_2(\%no_map);

################## SUBROUTINES ######################

# Usage
  sub Usage {             
  print STDERR "
Program description:
 \tCGCONV converts all-atom structures (in PBD or PQR format) to coarse-grained or multiscale
 \trepresentation.

 \tThe mapping of all-atom (AA) coordinates into coarse-grained (CG) centroids is performed at 
 \tresidue level following definitions contained in specific mapping files (MAP, see 
 \tCGCONV/maps/example.map). Several MAP files can be used at the same time by concatenating them
 \tin the command line. In case the mapping transformation involves a center of mass calculation
 \tthe file CGCONV/parms/atwm.tbl is requested in order to define the atom mass weight for the AA
 \ttypes. The script can not check or add missing atoms, so the user is warned to use protonated input
 \tstructures to assure the correct mapping. In case of alternate atom locations the 'A' position
 \twill be chosen.

 \tA multiscale system can be generated by selecting the AA residues to be mapped into CG beads.
 \tThe selection is read though the option -R by using the following syntax:
 \tI  \tThe string must not contain blank characters.
 \tII \tColons work as logical 'or' functions.
 \tIII\tResidues can be selected by number, chain identifier (ID) or both.
 \tIV \tTo select particular residue numbers from a chain use the chain ID(s) followed by the
 \t\tresidues numbers in between square brackets. For example, A[1-10] selects residues 1
 \t\tto 10 of chain A, while AB[1-10] selects residues 1 to 10 from chains A and B. The
 \t\tsame rules apply to numbers within and outside brackets. Only one level of bracketing
 \t\tis supported.
 \tV  \tDistributive and commutative properties apply to selections. So selection AB[1-10]
 \t\tor A[1-10],B[1-10] or A[1-10]B renders the same result.
 \tVI \tResidues outside selection will remain unchanged.
 
 \t\tSome examples:
 \t\t1-10,15       \tSelects residues 1 to 10 and 15 from any chain.
 \t\tA,B,C (or ABC)\tSelects all residues from chain A, B and C.
 \t\t1-10,15,ABC   \tSelects residues 1 to 10 and 15 from any chain and all residues
 		   \t\tfrom chains A, B and C.


Usage:\tcgconv.pl [-i] [-o] [-R] [-aPLhv] [MAP]... < STDIN > STDOUT

Options: These are the optional arguments

 -i\tInput file in pdb format, default STDIN (<)
 -o\tOutput file name, default STDOUT (>)
 -R\tApply mapping to selected residues. Default apply to all. This option sets -P on.
 -a\tFlag to append custom MAP file(s) to SIRAH maps instead of replacing them.
 -P\tFlag to force the printing of residues not found in MAP file(s) or selection.
 -L\tFlag to generate the cgconv.log file. The LOG file is useful to check the correct reading of
 \tthe MAP files.
 -h\tFlag to print help and exit.
 -v\tFlag to print version and exit.
 [MAP]\tPath to mapping file(s). MAP file(s) must be the last defined option. Files are first searched
 \tat the specified path, if not present there they are looked at repository CGCONV/maps/.
 \tIf no MAP file is set, then the default is to apply the SIRAH mapping by using CGCONV/maps/sirah_*.map files.
 

Examples:

  1.\tBasic usage: SIRAH mapping
 \tcgconv.pl -i all-atom.pdb -o sirah-cg.pdb

  2.\tResidue selection: Map residues 20 to 30 and 40 to 50 from chains A and B into CG beads
 \tcgconv.pl -i all-atom.pdb -o sirah-cg.pdb -R AB[20-30,40-50]
   
  3.\tCustom MAP files:
 \tcgconv.pl -i all-atom.pdb -o user-cg.pdb \$PATH/MAP1 \$PATH/MAP2
   
  4.\tCGCONV in pipeline:
 \t[do something] all-atom.pdb | cgconv.pl [options] | [do something else] > output.pdb
\n";

  exit 1;
  }

# Load data from Atom Type MW file
  sub LoadAtMW {

      open (ATMW ,"<$cgconv_path/parms/atmw.tbl") or die "Can't open atmw.tbl file for reading\n";
              
      while (<ATMW>) {

            # Ignore empty lines or lines starting with '#'
            if ( $_ =~ /^[\s]*[#]|^[\s]+$/ ) {next;}
 
            $_ =~ s/^\s+//; # remove leading whitespace
            $_ =~ s/\s+$//; # remove trailing whitespace return
                                       
            @table_field = split(/\s+/,$_);
            if ( !($table_field[1]=~ /^[0-9]+\.{0,1}[0-9]+$/) ) {&Error_3;}
 
            @atomtypes = @table_field[2 .. $#table_field];
            foreach $atomtype (@atomtypes) { $atomtypes{"$atomtype"} = $table_field[1]; }
      }

      close ATMW;
  }

# GetResid
  sub GetResid { # [resid selection]

      my $selection = $_[0];
      my @resid     = split(',', $selection);
      my $count     = 0;

      foreach $num (@resid) { if ($num =~ /([0-9]+)\-([0-9]+)/) {$resid[$count] = $1; push(@resid,($1 + 1)..$2);} ++$count; }

      return @resid;
  }

# Filter selection
  sub InSelection { # [chain] [resid] [%selection]
    
      my $chain =   $_[0]  ; #
      my $resid =   $_[1]  ; #
      my %selec = %{$_[2]} ; # Hash of selection
      my $eval  =      0   ; # FALSE = 0; TRUE = 1

      if (!(%selec)) {$eval = 1; return $eval;}

      if ( exists $selection{"$chain"} ){

         if ( $selection{"$chain"} eq '' ) { $eval = 1; }
    
         elsif ( grep (/^$resid$/,  @{ $selection{"$chain"} }) ) { $eval = 1; }
      }
    
      elsif ( grep (/^$resid$/,  @{ $selection{"*"} }) ) { $eval = 1; }
    
      return $eval;
  }

# PrintGroups
  sub PrintGroups { # [hash of groups]

      my %gr_db = %{$_[0]} ; # group data
    
      for my $gr_ndx ( sort (keys %gr_db) ) {
         
          if ($gr_db{"$gr_ndx"}{"mass"} == 0) {print STDERR "Several atom types without mass definition\n"; exit 1;}
         
          ++$atom_ndx; if ($atom_ndx > 99999) { $atom_ndx = 0; } ; # Reset counter
          printf $OUTPUT ("ATOM%7d%5s %-4s%s%4s%s%11.3f%8.3f%8.3f\n",

                           $atom_ndx,
                           $gr_db{"$gr_ndx"}{"cgattyp"},
                           $gr_db{"$gr_ndx"}{"cgres"},
                           $gr_db{"$gr_ndx"}{"chain"},
                           $gr_db{"$gr_ndx"}{"resid"},
                           $gr_db{"$gr_ndx"}{"achar"},
                         ( $gr_db{"$gr_ndx"}{"x"} / $gr_db{"$gr_ndx"}{"mass"} ),
                         ( $gr_db{"$gr_ndx"}{"y"} / $gr_db{"$gr_ndx"}{"mass"} ),
                         ( $gr_db{"$gr_ndx"}{"z"} / $gr_db{"$gr_ndx"}{"mass"} )
          );
      }
  }

# Add MAP to database
  sub AddMap { # [input hash] [output hash]

      my %CGRES = %{$_[0]};

    # Build map hash 
      for my $res ( @{$CGRES{"ALLNAME"}} ){

          $map_db{"$res"}{"CGRES"} = $CGRES{"CGNAME"};
          $map_db{"$res"}{"ATOMS"} = \%{ $CGRES{"ATOMS"} };
      }
  }

# Error messages
  sub Error_1 { print STDERR "Syntaxis error in map file (check LOG).\n"; if ($opts{L} == 1) {print LOG "\nSyntaxis error in line $.\n";} exit 1;}

  sub Error_2 { print STDERR "Incomplete definition of CG residue in map file (check LOG).\n"; if ($opts{L} == 1) {print LOG "Missing options!\n";} exit 1;}

  sub Error_3 { print STDERR "Error reading line $. in table $cgconv_path/parms/atmw.tbl\n"; exit 1;}

# Warning messages
  sub Warning_1 { print STDERR "Warning! No MW defined for atom type $atom[0], it will not contribute to center of mass.\n";}

  sub Warning_2 { # [input hash]

      my %list  = %{$_[0]} ; # hash list of not mapped residues
      my $count = 0        ;
    
      if (%list) {
    
         print STDERR "Warning! Some residues were not found in MAP files:";
    
         foreach $resname (keys %list) {

                 if (($count % 10) == 0){ print STDERR "\n"; }
                 printf STDERR "%-5s", $resname;
                 ++$count;
         }
       
         print STDERR "\n";
      }
  }

################ END OF THE CODE ####################
```
