## Supporting file - scripts for "Protocols for Multi-Scale Molecular Dynamics Simulations: A Comparative Study for Intrinsically Disordered Amyloid Beta in Amber & Gromacs on CPU & GPU": COPYING

		    GNU GENERAL PUBLIC LICENSE
		       Version 2, June 1991

 Copyright (C) 1989, 1991 Free Software Foundation, Inc.
     51 Franklin Street, Fifth Floor, Boston, MA  02110-1301  USA
 Everyone is permitted to copy and distribute verbatim copies
 of this license document, but changing it is not allowed.

			    Preamble

  The licenses for most software are designed to take away your
freedom to share and change it.  By contrast, the GNU General Public
License is intended to guarantee your freedom to share and change free
software--to make sure the software is free for all its users.  This
General Public License applies to most of the Free Software
Foundation's software and to any other program whose authors commit to
using it.  (Some other Free Software Foundation software is covered by
the GNU Library General Public License instead.)  You can apply it to
your programs, too.

  When we speak of free software, we are referring to freedom, not
price.  Our General Public Licenses are designed to make sure that you
have the freedom to distribute copies of free software (and charge for
this service if you wish), that you receive source code or can get it
if you want it, that you can change the software or use pieces of it
in new free programs; and that you know you can do these things.

  To protect your rights, we need to make restrictions that forbid
anyone to deny you these rights or to ask you to surrender the rights.
These restrictions translate to certain responsibilities for you if you
distribute copies of the software, or if you modify it.

  For example, if you distribute copies of such a program, whether
gratis or for a fee, you must give the recipients all the rights that
you have.  You must make sure that they, too, receive or can get the
source code.  And you must show them these terms so they know their
rights.

  We protect your rights with two steps: (1) copyright the software, and
(2) offer you this license which gives you legal permission to copy,
distribute and/or modify the software.

  Also, for each author's protection and ours, we want to make certain
that everyone understands that there is no warranty for this free
software.  If the software is modified by someone else and passed on, we
want its recipients to know that what they have is not the original, so
that any problems introduced by others will not reflect on the original
authors' reputations.

  Finally, any free program is threatened constantly by software
patents.  We wish to avoid the danger that redistributors of a free
program will individually obtain patent licenses, in effect making the
program proprietary.  To prevent this, we have made it clear that any
patent must be licensed for everyone's free use or not licensed at all.

  The precise terms and conditions for copying, distribution and
modification follow.

		    GNU GENERAL PUBLIC LICENSE
   TERMS AND CONDITIONS FOR COPYING, DISTRIBUTION AND MODIFICATION

  0. This License applies to any program or other work which contains
a notice placed by the copyright holder saying it may be distributed
under the terms of this General Public License.  The "Program", below,
refers to any such program or work, and a "work based on the Program"
means either the Program or any derivative work under copyright law:
that is to say, a work containing the Program or a portion of it,
either verbatim or with modifications and/or translated into another
language.  (Hereinafter, translation is included without limitation in
the term "modification".)  Each licensee is addressed as "you".

Activities other than copying, distribution and modification are not
covered by this License; they are outside its scope.  The act of
running the Program is not restricted, and the output from the Program
is covered only if its contents constitute a work based on the
Program (independent of having been made by running the Program).
Whether that is true depends on what the Program does.

  1. You may copy and distribute verbatim copies of the Program's
source code as you receive it, in any medium, provided that you
conspicuously and appropriately publish on each copy an appropriate
copyright notice and disclaimer of warranty; keep intact all the
notices that refer to this License and to the absence of any warranty;
and give any other recipients of the Program a copy of this License
along with the Program.

You may charge a fee for the physical act of transferring a copy, and
you may at your option offer warranty protection in exchange for a fee.

  2. You may modify your copy or copies of the Program or any portion
of it, thus forming a work based on the Program, and copy and
distribute such modifications or work under the terms of Section 1
above, provided that you also meet all of these conditions:

    a) You must cause the modified files to carry prominent notices
    stating that you changed the files and the date of any change.

    b) You must cause any work that you distribute or publish, that in
    whole or in part contains or is derived from the Program or any
    part thereof, to be licensed as a whole at no charge to all third
    parties under the terms of this License.

    c) If the modified program normally reads commands interactively
    when run, you must cause it, when started running for such
    interactive use in the most ordinary way, to print or display an
    announcement including an appropriate copyright notice and a
    notice that there is no warranty (or else, saying that you provide
    a warranty) and that users may redistribute the program under
    these conditions, and telling the user how to view a copy of this
    License.  (Exception: if the Program itself is interactive but
    does not normally print such an announcement, your work based on
    the Program is not required to print an announcement.)

These requirements apply to the modified work as a whole.  If
identifiable sections of that work are not derived from the Program,
and can be reasonably considered independent and separate works in
themselves, then this License, and its terms, do not apply to those
sections when you distribute them as separate works.  But when you
distribute the same sections as part of a whole which is a work based
on the Program, the distribution of the whole must be on the terms of
this License, whose permissions for other licensees extend to the
entire whole, and thus to each and every part regardless of who wrote it.

Thus, it is not the intent of this section to claim rights or contest
your rights to work written entirely by you; rather, the intent is to
exercise the right to control the distribution of derivative or
collective works based on the Program.

In addition, mere aggregation of another work not based on the Program
with the Program (or with a work based on the Program) on a volume of
a storage or distribution medium does not bring the other work under
the scope of this License.

  3. You may copy and distribute the Program (or a work based on it,
under Section 2) in object code or executable form under the terms of
Sections 1 and 2 above provided that you also do one of the following:

    a) Accompany it with the complete corresponding machine-readable
    source code, which must be distributed under the terms of Sections
    1 and 2 above on a medium customarily used for software interchange; or,

    b) Accompany it with a written offer, valid for at least three
    years, to give any third party, for a charge no more than your
    cost of physically performing source distribution, a complete
    machine-readable copy of the corresponding source code, to be
    distributed under the terms of Sections 1 and 2 above on a medium
    customarily used for software interchange; or,

    c) Accompany it with the information you received as to the offer
    to distribute corresponding source code.  (This alternative is
    allowed only for noncommercial distribution and only if you
    received the program in object code or executable form with such
    an offer, in accord with Subsection b above.)

The source code for a work means the preferred form of the work for
making modifications to it.  For an executable work, complete source
code means all the source code for all modules it contains, plus any
associated interface definition files, plus the scripts used to
control compilation and installation of the executable.  However, as a
special exception, the source code distributed need not include
anything that is normally distributed (in either source or binary
form) with the major components (compiler, kernel, and so on) of the
operating system on which the executable runs, unless that component
itself accompanies the executable.

If distribution of executable or object code is made by offering
access to copy from a designated place, then offering equivalent
access to copy the source code from the same place counts as
distribution of the source code, even though third parties are not
compelled to copy the source along with the object code.

  4. You may not copy, modify, sublicense, or distribute the Program
except as expressly provided under this License.  Any attempt
otherwise to copy, modify, sublicense or distribute the Program is
void, and will automatically terminate your rights under this License.
However, parties who have received copies, or rights, from you under
this License will not have their licenses terminated so long as such
parties remain in full compliance.

  5. You are not required to accept this License, since you have not
signed it.  However, nothing else grants you permission to modify or
distribute the Program or its derivative works.  These actions are
prohibited by law if you do not accept this License.  Therefore, by
modifying or distributing the Program (or any work based on the
Program), you indicate your acceptance of this License to do so, and
all its terms and conditions for copying, distributing or modifying
the Program or works based on it.

  6. Each time you redistribute the Program (or any work based on the
Program), the recipient automatically receives a license from the
original licensor to copy, distribute or modify the Program subject to
these terms and conditions.  You may not impose any further
restrictions on the recipients' exercise of the rights granted herein.
You are not responsible for enforcing compliance by third parties to
this License.

  7. If, as a consequence of a court judgment or allegation of patent
infringement or for any other reason (not limited to patent issues),
conditions are imposed on you (whether by court order, agreement or
otherwise) that contradict the conditions of this License, they do not
excuse you from the conditions of this License.  If you cannot
distribute so as to satisfy simultaneously your obligations under this
License and any other pertinent obligations, then as a consequence you
may not distribute the Program at all.  For example, if a patent
license would not permit royalty-free redistribution of the Program by
all those who receive copies directly or indirectly through you, then
the only way you could satisfy both it and this License would be to
refrain entirely from distribution of the Program.

If any portion of this section is held invalid or unenforceable under
any particular circumstance, the balance of the section is intended to
apply and the section as a whole is intended to apply in other
circumstances.

It is not the purpose of this section to induce you to infringe any
patents or other property right claims or to contest validity of any
such claims; this section has the sole purpose of protecting the
integrity of the free software distribution system, which is
implemented by public license practices.  Many people have made
generous contributions to the wide range of software distributed
through that system in reliance on consistent application of that
system; it is up to the author/donor to decide if he or she is willing
to distribute software through any other system and a licensee cannot
impose that choice.

This section is intended to make thoroughly clear what is believed to
be a consequence of the rest of this License.

  8. If the distribution and/or use of the Program is restricted in
certain countries either by patents or by copyrighted interfaces, the
original copyright holder who places the Program under this License
may add an explicit geographical distribution limitation excluding
those countries, so that distribution is permitted only in or among
countries not thus excluded.  In such case, this License incorporates
the limitation as if written in the body of this License.

  9. The Free Software Foundation may publish revised and/or new versions
of the General Public License from time to time.  Such new versions will
be similar in spirit to the present version, but may differ in detail to
address new problems or concerns.

Each version is given a distinguishing version number.  If the Program
specifies a version number of this License which applies to it and "any
later version", you have the option of following the terms and conditions
either of that version or of any later version published by the Free
Software Foundation.  If the Program does not specify a version number of
this License, you may choose any version ever published by the Free Software
Foundation.

  10. If you wish to incorporate parts of the Program into other free
programs whose distribution conditions are different, write to the author
to ask for permission.  For software which is copyrighted by the Free
Software Foundation, write to the Free Software Foundation; we sometimes
make exceptions for this.  Our decision will be guided by the two goals
of preserving the free status of all derivatives of our free software and
of promoting the sharing and reuse of software generally.

			    NO WARRANTY

  11. BECAUSE THE PROGRAM IS LICENSED FREE OF CHARGE, THERE IS NO WARRANTY
FOR THE PROGRAM, TO THE EXTENT PERMITTED BY APPLICABLE LAW.  EXCEPT WHEN
OTHERWISE STATED IN WRITING THE COPYRIGHT HOLDERS AND/OR OTHER PARTIES
PROVIDE THE PROGRAM "AS IS" WITHOUT WARRANTY OF ANY KIND, EITHER EXPRESSED
OR IMPLIED, INCLUDING, BUT NOT LIMITED TO, THE IMPLIED WARRANTIES OF
MERCHANTABILITY AND FITNESS FOR A PARTICULAR PURPOSE.  THE ENTIRE RISK AS
TO THE QUALITY AND PERFORMANCE OF THE PROGRAM IS WITH YOU.  SHOULD THE
PROGRAM PROVE DEFECTIVE, YOU ASSUME THE COST OF ALL NECESSARY SERVICING,
REPAIR OR CORRECTION.

  12. IN NO EVENT UNLESS REQUIRED BY APPLICABLE LAW OR AGREED TO IN WRITING
WILL ANY COPYRIGHT HOLDER, OR ANY OTHER PARTY WHO MAY MODIFY AND/OR
REDISTRIBUTE THE PROGRAM AS PERMITTED ABOVE, BE LIABLE TO YOU FOR DAMAGES,
INCLUDING ANY GENERAL, SPECIAL, INCIDENTAL OR CONSEQUENTIAL DAMAGES ARISING
OUT OF THE USE OR INABILITY TO USE THE PROGRAM (INCLUDING BUT NOT LIMITED
TO LOSS OF DATA OR DATA BEING RENDERED INACCURATE OR LOSSES SUSTAINED BY
YOU OR THIRD PARTIES OR A FAILURE OF THE PROGRAM TO OPERATE WITH ANY OTHER
PROGRAMS), EVEN IF SUCH HOLDER OR OTHER PARTY HAS BEEN ADVISED OF THE
POSSIBILITY OF SUCH DAMAGES.

		     END OF TERMS AND CONDITIONS

	    How to Apply These Terms to Your New Programs

  If you develop a new program, and you want it to be of the greatest
possible use to the public, the best way to achieve this is to make it
free software which everyone can redistribute and change under these terms.

  To do so, attach the following notices to the program.  It is safest
to attach them to the start of each source file to most effectively
convey the exclusion of warranty; and each file should have at least
the "copyright" line and a pointer to where the full notice is found.

    <one line to give the program's name and a brief idea of what it does.>
    Copyright (C) <year>  <name of author>

    This program is free software; you can redistribute it and/or modify
    it under the terms of the GNU General Public License as published by
    the Free Software Foundation; either version 2 of the License, or
    (at your option) any later version.

    This program is distributed in the hope that it will be useful,
    but WITHOUT ANY WARRANTY; without even the implied warranty of
    MERCHANTABILITY or FITNESS FOR A PARTICULAR PURPOSE.  See the
    GNU General Public License for more details.

    You should have received a copy of the GNU General Public License
    along with this program; if not, write to the Free Software
    Foundation, Inc., 51 Franklin Street, Fifth Floor, Boston, MA  02110-1301  USA


Also add information on how to contact you by electronic and paper mail.

If the program is interactive, make it output a short notice like this
when it starts in an interactive mode:

    Gnomovision version 69, Copyright (C) year  name of author
    Gnomovision comes with ABSOLUTELY NO WARRANTY; for details type `show w'.
    This is free software, and you are welcome to redistribute it
    under certain conditions; type `show c' for details.

The hypothetical commands `show w' and `show c' should show the appropriate
parts of the General Public License.  Of course, the commands you use may
be called something other than `show w' and `show c'; they could even be
mouse-clicks or menu items--whatever suits your program.

You should also get your employer (if you work as a programmer) or your
school, if any, to sign a "copyright disclaimer" for the program, if
necessary.  Here is a sample; alter the names:

  Yoyodyne, Inc., hereby disclaims all copyright interest in the program
  `Gnomovision' (which makes passes at compilers) written by James Hacker.

  <signature of Ty Coon>, 1 April 1989
  Ty Coon, President of Vice

This General Public License does not permit incorporating your program into
proprietary programs.  If your program is a subroutine library, you may
consider it more useful to permit linking proprietary applications with the
library.  If this is what you want to do, use the GNU Library General
Public License instead of this License.
