## Supporting file - scripts for "Protocols for Multi-Scale Molecular Dynamics Simulations: A Comparative Study for Intrinsically Disordered Amyloid Beta in Amber & Gromacs on CPU & GPU": g_top2psf.pl

```
#!/usr/bin/perl

#----------------------------------------------------------------------#
# G_TOP2PSF                                                            #
#----------------------------------------------------------------------#
# Program to convert gromacs topologies to X-PLOR psf files.           #
#                                                                      #
# USAGE:   top2psf.pl -i TOP -o PSF [-c] [-kdpshv]                     #
#                                                                      #
# AUTHOR:  Matias Machado                                              #
# E-MAIL:                                      #
# VERSION: 8.6, Jan 2020                                               #
#                                                                      #
# This program is free software; you can redistribute it and/or modify #
# it under the terms of the GNU General Public License as published by #
# the Free Software Foundation; either version 2 of the License, or    #
# (at your option) any later version.                                  #
#                                                                      #
# This program is distributed in the hope that it will be useful,      #
# but WITHOUT ANY WARRANTY; without even the implied warranty of       #
# MERCHANTABILITY or FITNESS FOR A PARTICULAR PURPOSE. See the         #
# GNU General Public License for more details.                         #
#----------------------------------------------------------------------#

#--------- Import modules --------------------------#

use File::Basename ;
use Getopt::Std    ;

#--------- Program run starts here -----------------#

%opts = ();
getopts("i:o:c:kdpshv",\%opts);

# Checking opts arguments
if ( $opts{"v"} ) { print STDERR "version 8.6 [Jan 2020]\n"; exit 1 ; }
if ( ($opts{"h"} == 1) || (!($opts{"i"})) || (!($opts{"o"})) ) {&Usage;}


#--------- Load data from TOP file -----------------#

$GMXDATA = $ENV{'GMXDATA'} ; # GROMACS data folder

# GROMACS ff directory
if ( $GMXDATA ) {

     if    ( -e $GMXDATA . "/top"         ) { $GMXLIB = $GMXDATA . "/top/"        ; }
     elsif ( -e $GMXDATA . "/gromacs/top" ) { $GMXLIB = $GMXDATA . "/gromacs/top/"; }

} else { $GMXLIB = ''; }

&ReadTop( $opts{"i"}, $GMXLIB, '') ;


#--------- Build conectivities ---------------------#

$atom_ndx =  0; # atom index
$res_ndx  =  0; # residue index
@atoms    = ();
@bonds    = ();
@angles   = ();
@die      = ();
@dist_rst = ();
@segname  = (A..Z,a..z);

if ( $opts{"c"} ) { @segname = split(/,/,$opts{"c"}) };

$molecules = each_array( \@{$MOLECULES[0]}, \@{$MOLECULES[1]} );

while ( my ($mol,$num) = $molecules->() ) {

    if (!($MOL_db{"$mol"})) { print STDERR "Error! Molecule $mol not found in toplogy database.\n"; exit 1;}

    for ( $i=1; $i<=$num; $i++ ) {

        $mol_atoms = (keys %{$MOL_db{"$mol"}{"ATOMS"}}); # Total atoms in the molecule
        
        $prev_res = ''; # previous residue index
        
        # ATOMS
        for ( $at=1; $at<=$mol_atoms; $at++ ) {
            
            # Set residue index
            if ($opts{"k"}) {$res_ndx = $MOL_db{"$mol"}{"ATOMS"}{"$at"}{"RESID"};}
            
            elsif ($prev_res ne $MOL_db{"$mol"}{"ATOMS"}{"$at"}{"RESID"}) {
            
                   $prev_res = $MOL_db{"$mol"}{"ATOMS"}{"$at"}{"RESID"};
            
                   ++$res_ndx; if ($res_ndx > 9999) { $res_ndx = 1; }; # PSF file format limit
            }
            
            push ( @atoms, ( 
                             $at + $atom_ndx,
                             $segname[0],
                             $res_ndx,
                             $MOL_db{"$mol"}{"ATOMS"}{"$at"}{"RESNAME"},
                             $MOL_db{"$mol"}{"ATOMS"}{"$at"}{"NAME"},
                             $MOL_db{"$mol"}{"ATOMS"}{"$at"}{"TYPE"},
                             $MOL_db{"$mol"}{"ATOMS"}{"$at"}{"CHARGE"},
                             $MOL_db{"$mol"}{"ATOMS"}{"$at"}{"MASS"},
                             0
                           )
            );
            
        }
        
        # BONDS
        push (@bonds, map {$_ + $atom_ndx} @{$MOL_db{"$mol"}{"BONDS"}} );
        
        # ANGELS
        push (@angles, map {$_ + $atom_ndx} @{$MOL_db{"$mol"}{"ANGLES"}} );
                
        # DIE
        push (@die, map {$_ + $atom_ndx} @{$MOL_db{"$mol"}{"DIE"}} );
        
        # DIST_RST
        push (@dist_rst, map {$_ + $atom_ndx} @{$MOL_db{"$mol"}{"DIST_RST"}} );
        
        # PAIRS
        push (@pairs, map {$_ + $atom_ndx} @{$MOL_db{"$mol"}{"PAIRS"}} );
        
        $atom_ndx = $atom_ndx + $mol_atoms;
    }

    push(@segname,shift(@segname)) ; # Set circular array
}

#--------- Print PSF file --------------------------#

# Open output file
open ($PSF ,">$opts{o}") or die "Can't open psf file $opts{o} for writting\n";

if ($opts{"d"}) { push (@bonds, @dist_rst); }
if ($opts{"p"}) { push (@bonds, @pairs   ); }

printf $PSF ( "PSF\n\n       3 !NTITLE\n REMARKS PSF file generated by g_top2psf.pl\n REMARKS SIRAH Tools Kit [by M.R. Machado | doi: 10.1093/bioinformatics/btw020]\n REMARKS GROMACS topology: %s\n\n",$opt{"i"} );

$format =   ( "%8s %-4s %-4s %-4s %-4s %-4s %10s %13.2f %11d\n" x (@atoms/9) );
printf $PSF ( "%8d !NATOM\n", (@atoms/9) );
printf $PSF ( $format, @atoms );
printf $PSF   "\n";

$format =   ( "%8d %7d %7d %7d %7d %7d %7d %7d\n" x (@bonds/8) ); if (@bonds % 8) { $format .= ("%8d" x (@bonds % 8)) . "\n" ; }
printf $PSF ( "%8d !NBOND: bonds\n", (@bonds/2) );
printf $PSF ( $format, @bonds );
printf $PSF   "\n";

$format =   ( "%8d %7d %7d %7d %7d %7d %7d %7d %7d\n" x (@angles/9) ); if (@angles % 9) { $format .= ("%8d" x (@angles % 9)) . "\n" ; }
printf $PSF ( "%8d !NTHETA: angles\n", (@angles/3) );
printf $PSF ( $format, @angles );
printf $PSF   "\n";

$format =   ( "%8d %7d %7d %7d %7d %7d %7d %7d\n" x (@die/8) ); if (@die % 8) { $format .= ("%8d" x (@die % 8)) . "\n" ; }
printf $PSF ( "%8d !NPHI: dihedrals\n", (@die/4) );
printf $PSF ( $format, @die );
printf $PSF   "\n";

close $PSF;

#--------- SUBROUTINES -----------------------------#

# Usage
sub Usage {             

print STDERR "
Program description:
 \tG_TOP2PSF converts gromacs (GMX) topologies to X-PLOR PSF files.
 \tGenerated PSF files are only intended for visualization purposes.
 \tThe script does not check the correctness of the topology format.
 \tComment lines are omitted. Files included in topologies are
 \tprocessed by first looking in the local PATH and then in GMX
 \tfolders (provided GMXDATA environment variable is defined). In case
 \tthe file is not found, a warning message is printed. If the file
 \tdoes not contain relevant information on molecule topology then use
 \tflag -s to suppres the message. The PSF is made according to the
 \t[ molecules ] section. To generate a stripped PSF just comment the
 \tlines of the undesired molecules in [ molecules ] section.
 \tBy default, residues are renumbered starting from 1, the index is
 \treset to 1 whenever the residue counter exceeds 9999 to preserve
 \tthe PSF file format. Use option -k to keep the numeration in the
 \tTOP file. Molecules' chains are named from A-Z to a-z, unless
 \toption -c is set.
 
Usage:\tg_top2psf.pl -i TOP -o PSF [-c] [-dpshv]

Options:

 -i\tInput: gromacs topology file.
 -o\tOutput: psf file name.
 -k\tFlag to keep residue indices from TOP file.
 -c\tChain names: String of names separated by colons (e.g. X,Y,Z)
 -d\tAppend distant restraint information as bonds to psf.
 -p\tAppend pairs information as bonds to psf.
 -s\tSuppress warning messages about nonexistent or unreadable files.
 -h\tFlag to print help and exit.
 -v\tFlag to print version and exit.
\n";

exit 1;

}

sub ReadTop { # Usage: ReadTop [file] [defpath] [localdir]

    my $file     = $_[0] ; # Input file
    my $defpath  = $_[1] ; # Default GMX path
    my $localdir = $_[2] ; # Path

    my ($name,$path) = fileparse($file);

    # Open input file
    my $TOP;

    if    (open ($TOP ,"<",($localdir . $file))) { $localdir = ($localdir . $path); }
    elsif (open ($TOP ,"<",($defpath  . $file))) { $localdir = ($defpath  . $path); }
    elsif (!($opts{"s"})) { print STDERR "Warning! Can't open file $name for reading\n" ;}

    while (<$TOP>) {

       $_ =~ s/^\s+//; # remove leading whitespace
       $_ =~ s/\s+$//; # remove trailing whitespace & return

       if ( $_ =~ /^#include[\s]+\"(.+)\"$/) { &ReadTop( $1, $defpath, $localdir); next; }

       # Ignore empty lines or lines starting with ';' or '#'
       if ( $_ =~ /^[;#]|^$/ ) {next;}

       # %MOL_db
       #  MOL_NAME = ( 
       #               ATOMS  = ( # index
       #                            1   =  ( 
       #                                     NAME    = value,
       #                                     TYPE    = value,
       #                                     CHARGE  = value,
       #                                     MASS    = value,
       #                                     RESID   = value,
       #                                     RESNAME = value,
       #                                   )
       #                            ...
       #                        )
       #
       #               BONDS    = (@),
       #               ANGLES   = (@),
       #               DIE      = (@),
       #               DIST_RST = (@),
       #             ) ;

       # @MOLECULES = ((Compound,...), (#mols,...))

       if ( $_ =~ /^\[\s*([a-z_0-9]+)\s*\]/ ) { $section = $1; next; }

       my @data = split (/\s+/, $_);

       if    ($section eq 'moleculetype') { $MOL_NAME = $data[0]; }

       elsif ($section eq 'atoms') {

              $MOL_db{"$MOL_NAME"}{"ATOMS"}{$data[0]}{"TYPE"}    = $data[1];
              $MOL_db{"$MOL_NAME"}{"ATOMS"}{$data[0]}{"RESID"}   = $data[2];
              $MOL_db{"$MOL_NAME"}{"ATOMS"}{$data[0]}{"RESNAME"} = $data[3];
              $MOL_db{"$MOL_NAME"}{"ATOMS"}{$data[0]}{"NAME"}    = $data[4];
              $MOL_db{"$MOL_NAME"}{"ATOMS"}{$data[0]}{"CHARGE"}  = $data[6];
              $MOL_db{"$MOL_NAME"}{"ATOMS"}{$data[0]}{"MASS"}    = $data[7];
       }

       elsif ($section eq 'bonds')               { push ( @{$MOL_db{"$MOL_NAME"}{"BONDS"}}   , @data[0..1] ); }
       elsif ($section eq 'angles')              { push ( @{$MOL_db{"$MOL_NAME"}{"ANGLES"}}  , @data[0..2] ); }
       elsif ($section eq 'dihedrals')           { push ( @{$MOL_db{"$MOL_NAME"}{"DIE"}}     , @data[0..3] ); }
       elsif ($section eq 'distance_restraints') { push ( @{$MOL_db{"$MOL_NAME"}{"DIST_RST"}}, @data[0..1] ); }
       elsif ($section eq 'pairs')               { push ( @{$MOL_db{"$MOL_NAME"}{"PAIRS"}}   , @data[0..1] ); }

       elsif ($section eq 'molecules')           { push(@{$MOLECULES[0]},$data[0]); push(@{$MOLECULES[1]},$data[1]); }

    }

    close $TOP;

}

sub each_array {
# http://stackoverflow.com/questions/3208379/in-perl-how-can-i-iterate-over-multiple-elements-of-an-array
  my @copy = @_;
  my $i;
  my $max;

  for (map scalar @$_, @copy) {
    $max = $_ unless defined $max and $max > $_;
  }

  sub {
    return $i if @_ and shift eq 'index';
    my $new_i = defined $i ? $i + 1 : 0;
    return if $new_i >= $max;
    $i = $new_i;
    return map $_->[$i], @copy;
  }
}

#--------- END OF THE CODE -------------------------#
```
