## Supporting file - scripts for "Protocols for Multi-Scale Molecular Dynamics Simulations: A Comparative Study for Intrinsically Disordered Amyloid Beta in Amber & Gromacs on CPU & GPU": sirah_vmdtk.tcl

```
#----------------------------------------------------------------------#
# SIRAH Tools: mapping, backmapping and visualization of CG models     #
#----------------------------------------------------------------------#
# AUTHOR:    Matias Machado                                            #
# E-MAIL:                                      #
# VERSION:   1.0.3, Dec 2016                                           #
#                                                                      #
# REFERENCE: Machado M, Pantano S. Bioinformatics (2016)               #
#            DOI: 10.1093/bioinformatics/btw020                        #
#                                                                      #
# This program is free software; you can redistribute it and/or modify #
# it under the terms of the GNU General Public License as published by #
# the Free Software Foundation; either version 2 of the License, or    #
# (at your option) any later version.                                  #
#                                                                      #
# This program is distributed in the hope that it will be useful,      #
# but WITHOUT ANY WARRANTY; without even the implied warranty of       #
# MERCHANTABILITY or FITNESS FOR A PARTICULAR PURPOSE. See the         #
# GNU General Public License for more details.                         #
#                                                                      #
#----------------------------------------------------------------------#

proc sirah_help {{tool sirah_help}} {
# SIRAH Tools User Manual

# Manual Entries:

set sirah_docs(sirah_help)    \
"
>>>> sirah_help <<<<

Manual version: 1.1 \[Mar 2019\]

Description:

  Command to access the User Manual pages of SIRAH Tools

Usage: sirah_help CommandName

CommandName       Function
-----------------------------------------------------------------
  sirah_ss        Calculate secondary structure in SIRAH proteins
  sirah_backmap   Backmap coarse-grained to atomistic systems
  sirah_restype   Set SIRAH residue types
  sirah_radii     Set SIRAH vdW radii
  sirah_macros    Set useful selection macros
  sirah_help      Access User Manual pages
-----------------------------------------------------------------
"
#----------------------------
set sirah_docs(sirah_ss)      \
"
>>>> sirah_ss <<<<

version: 1.1 \[Dec 2015\]

Description:

  Command to calculate the secondary structure of SIRAH proteins.
  
  If mol is omitted then all options are applied to top molecule.
  By default the secondary structure data is stored in memory
  as the array sscache_data(mol,frame) and the coloring method
  'Secondary Structure' will display correctly. By default the
  script reads the sscache_data if available in memory and does
  not recalculate the conformation, unless flag ramach is set or
  the sscache_data is cleaned. The outname and noprint keywords
  control the generated output files.

  The current command version does not support PBC options.

Usage: sirah_ss \[options\]

Options: These are the optional arguments

  mol       Set the molecule ID, default top
  
  first     First frame to analyze
  
  last      Last frame to analyze
  
  sel       \"VMD selection\", default \"all\"
  
  outname   List {} of keywords and names for output files.
            Keywords and default names:
            
              byframe   ss_by_frame.xvg
              byres     ss_by_res.xvg
              global    ss_global.xvg
              mtx       ss.mtx
              phi       phi.mtx
              psi       psi.mtx
  
  load      Load secondary structure data from file (e.g ss.mtx) into molecule (mol)
  
  noprint   Flag to avoid printing standard results to out files
  
  ramach    Flag to write phi and psi angles of residue selection (sel) to out files
  
  nocahe    Flag to avoid saving data to sscache_data
  
  now       Flag to calculate the secondary structure at current frame.
            No sscache_data or output file is saved.
  
  clean     Flag to clean the sscache_data of the selected molecule (mol) and exit
  
  help      Display help and exit


Examples:

  1.        Load secondary structure data into molecules 2 and extend the calculation
            sirah_ss mol 2 load ss.mtx
            sirah_ss mol 2

  2.        Set custom output file names
            sirah_ss outname {mtx myss.mtx byframe myss_by_frame.xvg}
"
#----------------------------  
set sirah_docs(sirah_backmap) \
"
>>>> sirah_backmap <<<<

version: 1.0 \[Nov 2015\]

Description:

  Command to recover atomistic information from coarse-grained or multiscale
  systems.
  
  Geometric operations are applied to reconstruct the atomic coordinates then
  a minimization is performed to refine the structure of the system.
  The minimization requires AMBERTOOLS 14 (free at http://ambermd.org/)
  or later properly installed. MPI option requires mpirun and parallel
  compilation of AMBER code. Be aware that small systems may fail to run or
  converge in parallel execution due to decomposition problems. Notice, the
  cuda version requires installing the AMBER licensed suite.
  
  By default the force field ff14SB is used for the atomistic refinement,
  any residue or unit not defined within it will generate an execution error.
  The minimization protocol consists on 100+50 steps of steepest descent and
  conjugate gradient in vacuum conditions with a 0.12 nm cut-off for
  electrostatic.

  The current command version does not support PBC options, so make sure the
  molecules are whole before running the backmap.
  
Usage: sirah_backmap \[options\]

Options: These are the optional arguments

  mol       Set the molecule ID, default top.

  now       Backmap the current frame.
  
  first     First frame to process.
  
  last      Last frame to process.
  
  each      Process frames each number of steps, default 1.

  frames    List of frames to process, e.g. {1 2 10 21 22 23 30}.
  
  outname   Root names for output PDB file.
  
  noload    Flag to avoid loading the atomistic trajectory to the VMD session.
  
  nomin     Flag to avoid minimizing the system.
  
  mpi       MPI processes to use during minimization, default 1.

  cuda      Flag to use pmemd.cuda, sets gbsa on, cutoff to 999 and no MPI.

  gbsa      Flag to use implicit solvation GBSA (igb=1), default off (igb=0).

  cutoff    Set cut-off value (in angstroms) for non-bonded interactions, default 12.

  maxcyc    Set total number of minimization steps, default 150.

  ncyc      Set the initial number of steepest descent steps, default 100.
  
  help      Display help and exit.
" 

#----------------------------
set sirah_docs(sirah_restype) \
"
>>>> sirah_restype <<<<

version: 1.1 \[Feb 2015\]

Description:

  Command to set the residue types of SIRAH forcefield, which enables
  coloring method by \"Res type\".
  
  This command is applied by default when loading SIRAH Tools

"
#----------------------------
set sirah_docs(sirah_radii)   \
"
>>>> sirah_radii <<<<

version: 1.2 \[Dec 2015\]

Description:

  Command to set vdW radii of the SIRAH atom types, which has
  effects on drawing methods VDW, CPK, Surf and QuickSurf.
  Coloring method by \"Element\" is also enabled.
  
  This command is applied by default when loading SIRAH Tools

"

set sirah_docs(sirah_macros)  \
"
>>>> sirah_macros <<<<

version 1.3 \[Dec 2016\]

Description:
  Command to set useful selection macros:

  selection        Description
  ------------------------------
  sirah_membrane   Lipid residues
  sirah_nucleic    Nucleic residues
  sirah_protein    Protein residues
  sirah_basic      Basic protein residues
  sirah_acidic     Acidic protein residues
  sirah_polar      Polar protein residues
  sirah_neutral    Neutral protein residues
  sirah_backbone   Protein backbone atoms
  sirah_water      Water residues
  sirah_ions       Ion residues

  This command is applied by default when loading SIRAH Tools

"

if {[info exists sirah_docs($tool)]} {

     puts $sirah_docs($tool)

} else { puts stderr "No manual entry for $tool" }

}


proc getopts {command_args args} {
   # Usage: array set arrName [ getopts command_args switch {flags} option {opt} ]
   #        $arrName($flag), $arrName($opt), $arrName()

     set option [lsearch -exact $args option]
     set switch [lsearch -exact $args switch]

   # OPTIONS
     if {$option >= 0} {

         foreach opt [lindex $args [expr {$option + 1}]] {

              set pos [lsearch -exact $command_args $opt]
              if {$pos >= 0} {

                  set opts($opt) [lindex $command_args [expr {$pos + 1}]]
                  set command_args [lreplace $command_args $pos [expr {$pos + 1}]]
              }
         }
     }

   # SWITCHES
     if {$switch >= 0} {

         foreach opt [lindex $args [expr {$switch + 1}]] {

              set pos [lsearch -exact $command_args $opt]
              if {$pos >= 0} { 
              
                  set opts($opt) 1
                  set command_args [lreplace $command_args $pos $pos]
              }
         }
     }

   # UNKNOWN
     if {[llength $command_args] > 0} {set opts() $command_args}

     return [array get opts]
}


proc sirah_ss_ramach {phi psi} {
   # Clasify secondary structure elements using a "STRIDE" criteria
   # http://en.wikipedia.org/wiki/Ramachandran_plot#mediaviewer/File:1axc_PCNA_ProCheck_Rama.jpg

     if       {($phi >= -180 && $phi <= 10)               &&  ($psi >= -120 && $psi <=   45)}  { return "H"
     } elseif {($phi >= -180 && $phi <=  0 || $phi > 135) && (($psi >=   45 && $psi <=  180)   ||
                                                              ($psi >= -180 && $psi <= -120))} { return "E"
     } else   { return "C" }
}


proc sirah_ss_load {molid mtx} {

     if {! [file exists $mtx]} { 
         puts stderr "Error! Can't read file $mtx"
         return
     }
     
     if {! [string compare $molid top ]} { set molid [molinfo top] }
      
     global sscache_data
   
     set CA      [atomselect $molid "name GC"]
     set resnum  [expr {[llength [$CA get residue]] + 1}]
   
     set ssmtx   [open $mtx r]
     
     while {1} {
              
             set frame [gets $ssmtx]; if {[eof $ssmtx]} { close $ssmtx; break }
             if {! [string compare [lindex $frame 0] "#"]} {continue}
             if {[llength $frame] != $resnum } { puts stderr "Error! The number of residues in $mtx and mol $molid differs"; return}

             set sscache_data($molid,[lindex $frame 0]) [lrange $frame 1 [llength $frame]]
             
     }

     if {[info exists sscache_data($molid,[molinfo $molid get frame])]} {
         
          $CA set structure $sscache_data($molid,[molinfo $molid get frame])
     }

     start_sscache
}


proc sirah_ss {args} {
   # version 10.8 [Dec 2015]
   # Timeline 2.3 is highly dependent on selection (name CA protein nucleic) ss.tml does not work
   # sirah_ss { mol first {} last {} sel {} outname {byframe byres global mtx phi psi} load {} now noprint nocache clean help ramach}
   
   # Read arguments --------------------------------------------------------------------------------------------
     array set opts [ getopts $args switch { now nocache clean noprint ramach help } option { first last sel outname load mol } ]

     if { [info exists opts()]}            { puts stderr "Error! Bad argument: [join $opts()]"; return }
     
     if { [info exists opts(help)]}        { sirah_help sirah_ss; return }
     
     if { [info exists opts(mol)] && $opts(mol) != "top" } { set molid $opts(mol) } else { set molid [molinfo top] }
     
     if { [info exists opts(clean)]}       { global sscache_data; array unset sscache_data $molid,* ; return }

     if { [info exists opts(load)]}        { sirah_ss_load $molid $opts(load); return }
     
     if {![info exists opts(sel)]}         { set opts(sel) "all" }
     
     if { [info exists opts(now)]}         { sirah_ss mol $molid first [molinfo $molid get frame] last [molinfo $molid get frame] sel $opts(sel) nocache noprint; return }
     
     if { [info exists opts(first)]}       { set first $opts(first) } else { set first 0 }

     if { [info exists opts(last) ]}       { set last  $opts(last)  } else { set last [expr {[molinfo $molid get numframes] -1}] }
     
     if { [info exists opts(outname)]}     { array set outname [ getopts $opts(outname) option {mtx global byframe byres phi psi}]}
     
     if { [info exists outnames()]}        { puts stderr "Error! Bad argument: [join $outnames()]"; return }

     if {![info exists outname(byframe)]}  { set outname(byframe) ss_by_frame.xvg }

     if {![info exists outname(byres)]}    { set outname(byres)   ss_by_res.xvg   }

     if {![info exists outname(global)]}   { set outname(global)  ss_global.xvg   }

     if {![info exists outname(mtx)]}      { set outname(mtx)     ss.mtx          }
     
     if {![info exists outname(phi)]}      { set outname(phi)     phi.mtx         }
     
     if {![info exists outname(psi)]}      { set outname(psi)     psi.mtx         }
   #------------------------------------------------------------------------------------------------------------
   
     global sscache_data

   # Selections  
     set bbone   [[atomselect $molid "name GN GC GO"] get index]
     set GNbond  [[atomselect $molid "name GN"      ] getbonds ]
     set GObond  [[atomselect $molid "name GO"      ] getbonds ]
     set atm2res [[atomselect $molid "all"          ] get {residue fragment}]
     set CA       [atomselect $molid "name GC"      ]
     set PRO     [[atomselect $molid "name GC and resname sP"] get residue]
     set residue  [$CA get residue]
     set outres   [lsort -integer -uniq [[atomselect $molid "$opts(sel) and name GC"] get residue]]
     set curr_fr  [molinfo $molid get frame] ; # Current frame
     set num_fr   [expr $last - $first + 1]  ; # Total number of frames
   
   # Do some check  
     if {[llength $residue]  < 3} {puts stderr "Peptide too short"; return }
     if {[llength $outres ] == 0} {puts stderr "No protein residue selected"; return}

   # Scan chain
     set  outres_ndx 0
     set residue_ndx 0; # initial residue index in the list of residues
    
     foreach n $residue { 
     
          if {$n == [lindex $outres $outres_ndx]} {lappend outres_X 1; incr outres_ndx} else {lappend outres_X 0}
       
        # Build an array of indexes, i(ndx)
          set i($n) $residue_ndx; incr residue_ndx
        
        # NEW: Nt PRO requires spetial attention i(ndx,new)
          if {$n == [lindex $PRO 0]} {set PRO [lreplace $PRO 0 0]; set b 2 } else {set b 1}    ; # b = number of bonds 
          if {[llength [lindex $GNbond $i($n)]] == $b} {set i($n,new) 1} else {set i($n,new) 0}; # boolian 1="new chain", 0="same chain"

        # TER
          if {[llength [lindex $GObond $i($n)]] ==  1} {set i($n,ter) 1} else {set i($n,ter) 0}; # boolian 0="no TER", 1="TER"
     }
                            
   # Output files
     if {![info exists opts(noprint)]} {
     
         set  SSbyFrame  [open $outname(byframe) w]
         set  SSbyResid  [open $outname(byres)   w]
         set  SSGlobal   [open $outname(global)  w]
         set  SSMatrix   [open $outname(mtx)     w]

         puts $SSbyFrame [ format "\# SELECTION \"%s\"\n\#%-[string length $num_fr ]s %5s %5s %5s"           $opts(sel) " Frame" "H(%)" "E(%)" "C(%)" ]
         puts $SSbyResid [ format "\# SELECTION \"%s\"\n\#%-[string length [llength $residue]]s %5s %5s %5s" $opts(sel) " Resid" "H(%)" "E(%)" "C(%)" ]
         puts $SSGlobal  "\# SELECTION \"$opts(sel)\"\n\# Index  Mean(%)  Sd(% by frame)  Conformation"
         puts $SSMatrix  "\# Row = Frame; Columns = Residues"
     }
   
     if { [info exists opts(ramach)]} { 
         
         set  SSPhiMtx   [open $outname(phi) w]
         set  SSPsiMtx   [open $outname(psi) w]
     
         set  PhiMtx     {} ; # list of phi values of a frame
         set  PsiMtx     {} ; # list of psi values of a frame
     
         puts $SSPhiMtx  "\# SELECTION \"$opts(sel)\"\n\# Row = Frame; Columns = Residues"
         puts $SSPsiMtx  "\# SELECTION \"$opts(sel)\"\n\# Row = Frame; Columns = Residues"
     }
          
   
   # Startup counters
     foreach {H_sum H_csum E_sum E_csum C_sum C_csum} {0 0 0 0 0 0} {}
     foreach n $residue {set X($n,"H") 0; set X($n,"E") 0; set X($n,"C") 0}

   # Progress Bar
     set bar    "||||||||||||||||||||"     ;# Progress bar
     set tick    0                         ;# Starting tick mark in bar
     set N      [expr {$last - $first +1}] ;# Total number of frames

     if {$N > 1} { puts -nonewline "         0    25   50   75   100 %\nProgress |" }

   # Loop over frames and residues
     for {set frame $first} {$frame <= $last} {incr frame} {

        # Start counters
          set  H_frame      0 
          set  E_frame      0
          set  C_frame      0
          set  ss          {} ; # list  of conformations 
          set  res_ss("H") {} ; # array of SS containig residues
          set  res_ss("E") {} ;
          set  res_ss("C") {} ;
          
        # By default use sscache if available, unless flag ramach is set
          if { [info exists sscache_data($molid,$frame)] && ![info exists opts(ramach)] } {
          
               set ss $sscache_data($molid,$frame)
               
               foreach n $residue struc $ss { 
                    
                    if {[lindex $outres_X $i($n)]} {
                    
                        switch -exact $struc {
                       
                                H  {incr H_frame; incr X($n,"H")}
                                E  {incr E_frame; incr X($n,"E")}
                                C  {incr C_frame; incr X($n,"C")}
                        }
                    }
               }
          
          } else {

        # Calculate secondary structure
          foreach n $residue {

             # NEW chain
               if {$i($n,new)} {continue}
             
             # TER
               if {$i($n,ter)} {
                   
                   if {$i([expr {$n -1}],new)} {; # Dipeptide within a larger protein (e.g. PDB: 1T6O)
                       
                       set ss          [concat $ss {E E}] ; # give the chance to form B-sheet
                       set res_ss("E") [concat $res_ss("E") "[expr {$n -1}] $n"]
                       continue
                   }
                   
                   lappend ss $conf;
                   lappend res_ss("$conf") $n
                   continue
               }

             # Get index
               set phi_ndx [lrange $bbone [expr { $i($n)      * 3 - 1}] [expr {($i($n) + 1) * 3 - 1}]] ; # PHI: GO(-1)-GN-GC-GO
               set psi_ndx [lrange $bbone [expr {($i($n) + 1) * 3 - 3}] [expr {($i($n) + 2) * 3 - 3}]] ; # PSI:        GN-GC-GO-GN(+1)

             # Measure Phi and Psi and add a correction
               set phi [expr {[measure dihed $phi_ndx molid $molid frame $frame ] - 20}]
               set psi [expr {[measure dihed $psi_ndx molid $molid frame $frame ] + 13}]

               if       {$phi < -180} {set phi [expr {$phi + 360}]
               } elseif {$phi >  180} {set phi [expr {$phi - 360}]}

               if       {$psi < -180} {set psi [expr {$psi + 360}] 
               } elseif {$psi >  180} {set psi [expr {$psi - 360}]}

             # Store phi and psi values
               if {[info exists opts(ramach)] && [lsearch -exact $outres $n] >= 0 } { 
                   
                   lappend PhiMtx [format "%.1f" $phi]
                   lappend PsiMtx [format "%.1f" $psi]
               }
               
             # Set secondary structure Ramachandran ----------------------------
               set conf [sirah_ss_ramach $phi $psi]
               
               if {$i([expr {$n -1}],new)} {lappend ss $conf; lappend res_ss("$conf") [expr {$n - 1}]}
               
               lappend ss $conf; lappend res_ss("$conf") $n
             # -----------------------------------------------------------------
          }

        # Set secondary structure hydrogen bond network ------------------------
          array unset res_hb
          
        # Alpha helix
          foreach n $res_ss("H") {
                   
               foreach {A D} [list [expr {$i($n) - 4}] $i($n) $i($n) [expr {$i($n) + 4}]] { ; # A: acceptor, D: donor
                    
                    if {[lindex $ss $A] != "" && [lindex $ss $D] != "" } {
                    
                         set GO [lindex $bbone [expr {($A + 1) * 3 - 1}]]
                         set GN [lindex $bbone [expr {($D + 1) * 3 - 3}]]
                  
                         if {[measure bond [list $GO $GN] molid $molid frame $frame] <= 4.0} {incr res_hb($n)}
                    }
               }
               
               if {![info exists res_hb($n)]} {
               
                   lset ss $i($n) "C"; 
                   if {[lindex $outres_X $i($n)]} { incr C_frame; incr X($n,"C") } 
               
               } else { 
               
                   if {[lindex $outres_X $i($n)]} { incr H_frame; incr X($n,"H") }
               }
          }
               
        # Beta sheet
          if {[llength $res_ss("E")] > 0} {

              set E_GO     [atomselect $molid "residue $res_ss("E") and name GO"    frame $frame]     ; # H acceptor
              set E_GX     [atomselect $molid "residue $res_ss("E") and name GN GC" frame $frame]     ; # H donors
          
              set contacts [measure contacts 4.001 $E_GX $E_GO] ; # fast algorithm to list neighbors
               
              foreach res1 [lindex $contacts 0] res2 [lindex $contacts 1]  {

                 # skip residues n+1,n+2 of the same chain
                   if {  ( [lindex $atm2res $res1 1] == [lindex $atm2res $res2 1] )  \
                      && ( [expr {abs([lindex $atm2res $res1 0] - [lindex $atm2res $res2 0])}] <= 2 ) } { continue }
 
                   incr res_hb([lindex $atm2res $res1 0])
                   incr res_hb([lindex $atm2res $res2 0])

              }
              
              foreach n $res_ss("E") {
               
                   if {![info exists res_hb($n)]} {
                   
                       lset ss $i($n) "C"
                       if {[lindex $outres_X $i($n)]} { incr C_frame; incr X($n,"C") }
                       
                   } else { 
                   
                       if {[lindex $outres_X $i($n)]} { incr E_frame; incr X($n,"E") }
                   }
              }
          
              $E_GO delete
              $E_GX delete
          }

        # Random Coil
          set outres_ndx 0
          
          foreach n $res_ss("C") {
               
               if {[lindex $outres_X $i($n)]} { incr C_frame; incr X($n,"C") }
          }
          
        # ----------------------------------------------------------------------
          }; # <<<<<< END Calculate secondary structure
          
          if {![info exists opts(nocache)]} { set sscache_data($molid,$frame) $ss }
          
          if {![info exists opts(noprint)]} {

              puts $SSbyFrame [ format "  %-[string length $num_fr]d %5.1f %5.1f %5.1f"\
                                          $frame\
                                          [expr {100.0*$H_frame/[llength $outres]}]\
                                          [expr {100.0*$E_frame/[llength $outres]}]\
                                          [expr {100.0*$C_frame/[llength $outres]}]\
                              ]
                              
              puts $SSMatrix  "$frame $ss"
          }
          
          if { [info exists opts(ramach)]} {
              
              puts $SSPhiMtx "$frame $PhiMtx"
              puts $SSPsiMtx "$frame $PsiMtx"
              
              set  PhiMtx {}
              set  PsiMtx {}
          }
          
        # Global results  
          set H_sum  [expr {$H_sum  + (100.0 * $H_frame / [llength $outres])}]
          set E_sum  [expr {$E_sum  + (100.0 * $E_frame / [llength $outres])}]
          set C_sum  [expr {$C_sum  + (100.0 * $C_frame / [llength $outres])}]
          set H_csum [expr {$H_csum + (100.0 * $H_frame / [llength $outres])**2}]
          set E_csum [expr {$E_csum + (100.0 * $E_frame / [llength $outres])**2}]
          set C_csum [expr {$C_csum + (100.0 * $C_frame / [llength $outres])**2}]

          if { $frame == $curr_fr } { $CA set structure $ss }

       # Print progress ------------
         set pg [expr {100*($frame - $first +1)/$N}] ;# %Progress

         if { ([expr {$pg/5}] > $tick) && ($N > 1)} { puts -nonewline [string range $bar $tick [expr {$pg/5}]]; set tick [expr {$pg/5 +1}]; flush stdout }
       # ---------------------------
     };  if {$N > 1} {puts ""}
   
     if {![info exists opts(nocache)]} { puts -nonewline "Starting sscache... "; start_sscache ; puts "Done!"}
   
   # PRINT TO CONSOLE  
     puts [format "SUMMARY: <H> %3.1f%% <E> %3.1f%% <C> %3.1f%%" [expr {$H_sum/$num_fr}] [expr {$E_sum/$num_fr}] [expr {$C_sum/$num_fr}]]
         
   # PRINT TO FILE
     if {![info exists opts(noprint)]} {
     
         foreach n $outres {
     
              puts $SSbyResid [ format "%-[string length [llength $residue]]d %5.1f %5.1f %5.1f"\
                                        [expr {$n + 1}]\
                                        [expr {100.0*$X($n,"H")/$num_fr}]\
                                        [expr {100.0*$X($n,"E")/$num_fr}]\
                                        [expr {100.0*$X($n,"C")/$num_fr}]\
                              ]
         }

       # Workaround to issues in variance calculation: (E(X^2) - [E(X)]^2)^1/2
       # https://en.wikipedia.org/wiki/Algebraic_formula_for_the_variance
       # https://stackoverflow.com/questions/10212213/round-number-to-2-decimal-places
       #
       # In some unhappy cases (e.g. calculations over few frames withoud SS change)
       # the operation (E(X^2) - [E(X)]^2) on big numbers and round errors may eventualy
       # return a very tiny negetive number (<1e-10) which should be zero but is not,
       # causing an "out of domain" error in the sqrt operation. An empirical way to
       # solve it is using a lower decimal precision before applying the sqrt.
       #
         set   dec 4;   # decimal digit precision (0.0001)
         puts  $SSGlobal  [format "  1%12.1f %6.1f %12s" [expr {$H_sum/$num_fr}] [expr {sqrt(round(10**$dec*(($H_csum/$num_fr) - ($H_sum/$num_fr)**2))/10.0**$dec)} ] "H"]
         puts  $SSGlobal  [format "  2%12.1f %6.1f %12s" [expr {$E_sum/$num_fr}] [expr {sqrt(round(10**$dec*(($E_csum/$num_fr) - ($E_sum/$num_fr)**2))/10.0**$dec)} ] "E"]
         puts  $SSGlobal  [format "  3%12.1f %6.1f %12s" [expr {$C_sum/$num_fr}] [expr {sqrt(round(10**$dec*(($C_csum/$num_fr) - ($C_sum/$num_fr)**2))/10.0**$dec)} ] "C"]

         close $SSbyFrame
         close $SSbyResid
         close $SSGlobal
         close $SSMatrix
     }
     
     if { [info exists opts(ramach)]} {
     
         close $SSPhiMtx
         close $SSPsiMtx
     }
}


proc start_sscache {{molid top}} {
   # Start the cache for a given molecule  
     
     global sscache_data
     
     if {![string compare $molid top]} { set molid [molinfo top] }
     
     global vmd_frame
     
   # Set a trace to detect when an animation frame changes
     trace variable vmd_frame($molid) w sscache
     
     return
}


proc sscache {name index op} {
   # when the frame changes, trace calls this function
   # name  == vmd_frame
   # index == molecule id of the newly changed frame
   # op    == w
    
     global sscache_data

   # Get protein CA atoms
     set CA [atomselect $index "name GC"]

   # Get new frame number
     set frame [molinfo $index get frame]

   # if ss data exists in cache
     if [info exists sscache_data($index,$frame)] {

	$CA set structure $sscache_data($index,$frame)
	return

     } else { $CA set structure C; return }
}


proc seq {from to {each 1}} {
     set out {}

     if {$each <  0} {set each [expr {$each * -1}]}
     if {$each == 0} {set each 1}

     if {$from <= $to} {  for {set i $from} {$i <= $to} {incr i  $each} {lappend out $i}
     } else            {  for {set i $from} {$i >= $to} {incr i -$each} {lappend out $i} }

     return $out
}


proc addatom {at3 at2 at1 r4 a4 d4} {
   # add at4
   # r4 distance  at4-at3
   # a4 angle     at4-at3-at2
   # d4 diehedral at4-at3-at2-at1

   # Computing reference frame
     set v12 [vecsub   $at1 $at2]
     set v32 [vecsub   $at3 $at2]

     set X   [veccross $v32 $v12]
     set Z   [veccross $X   $v32]

     set r23 [vecdist  $at2 $at3]
     set rXO [vecdist  $X {0 0 0}] ;# Ditance to Ori
     set rZO [vecdist  $Z {0 0 0}] ;# Ditance to Ori

     foreach {l1 m1 n1} [vecscale $X   [expr {1.0/$rXO}]] {} ;# faster than lassign
     foreach {l2 m2 n2} [vecscale $v32 [expr {1.0/$r23}]] {}
     foreach {l3 m3 n3} [vecscale $Z   [expr {1.0/$rZO}]] {}

   # Calculating coordinates of at4 in x'y'z' system.
   # get Pi
     global M_PI

     set d4  [expr {$d4*$M_PI/180.0}]
     set a4  [expr {(180.0 - $a4)*$M_PI/180.0}]

     set z   [expr {$r4 * sin($a4) * cos($d4)}]
     set x   [expr {$r4 * sin($a4) * sin($d4)}]
     set y   [expr {$r4 * cos($a4) + $r23}]

     set at4 [list \
                    [expr {$l1*$x + $l2*$y + $l3*$z + [lindex $at2 0]}]\
                    [expr {$m1*$x + $m2*$y + $m3*$z + [lindex $at2 1]}]\
                    [expr {$n1*$x + $n2*$y + $n3*$z + [lindex $at2 2]}]\
             ]

     return $at4
}

proc sirah_mapdb {} {
   # Building keywords {: set} {+ add} | P {: PX ...} N1 {+ {...} {...}} ...
   # Because "dict exists" only works on real dictionaries keyword nomap must have a value
   # https://groups.google.com/forum/#!topic/comp.lang.tcl/KoH-qTvwy0s
     global backmap

   # Amino-Acids
     dict set backmap                                            \
         sG  { name  GLY                                         \
               build {N   {: GN } CA {: GC} O {: GO} N+ {: GN/+} \
                      C   {+ {O   CA  N+  } {1.24  32.0    0.0}} \
                     }                                           \
               last  {build {N+  {: GN}}}                        \
               print {N CA C O}                                  \
             }

     dict set backmap                                            \
         sA  { name  ALA                                         \
               build {N   {: GN } CA {: GC} O {: GO} N+ {: GN/+} \
                                                                 \
                      C   {+ {O   CA  N+  } {1.24  32.0    0.0}} \
                      CB  {+ {CA  N   C   } {1.50 109.0 -120.0}} \
                     }                                           \
               last  {build {N+  {: GN}}}                        \
               print {N CA C O CB}                               \
             }

     dict set backmap                                            \
         sV  { name  VAL                                         \
               build {N   {: GN } CA {: GC} O {: GO} CB {: BCB}  \
                                                                 \
                      C   {+ {CA  N   CB  } {1.50 109.0  120.0}} \
                      CG1 {+ {CB  CA  N   } {1.50 109.0  180.0}} \
                      CG2 {+ {CB  CA  CG1 } {1.50 109.0  120.0}} \
                     }                                           \
               print {N CA C O CB CG1 CG2}                       \
             }

     dict set backmap                                            \
         sL  { name  LEU                                         \
               build {N   {: GN } CA {: GC} O {: GO} N+ {: GN/+} \
                      CG  {: BCG}                                \
                                                                 \
                      C   {+ {O   CA  N+  } {1.24  32.0    0.0}} \
                      CB  {+ {CA  N   C   } {1.50 109.0 -120.0}} \
                      CD1 {+ {CG  CB  CA  } {1.50 109.0  180.0}} \
                      CD2 {+ {CG  CB  CD1 } {1.50 109.0  120.0}} \
                     }                                           \
               last  {build {N+  {: GN}}}                        \
               print {N CA C O CB CG CD1 CD2}                    \
             }

     dict set backmap                                            \
         sI  { name  ILE                                         \
               build {N   {: GN } CA {: GC} O {: GO} N+ {: GN/+} \
                      CG1 {: BCG}                                \
                                                                 \
                      C   {+ {O   CA  N+  } {1.24  32.0    0.0}} \
                      CB  {+ {CA  N   C   } {1.50 109.0 -120.0}} \
                      CG2 {+ {CB  CA  CG1 } {1.50 109.0 -120.0}} \
                      CD1 {+ {CG1 CB  CG2 } {1.50 109.0  -70.0}} \
                     }                                           \
               last  {build {N+  {: GN}}}                        \
               print {N CA C O CB CG1 CG2 CD1}                   \
             } 
             # Note: sometimes CD1 is named CD

     dict set backmap                                            \
         sP  { name  PRO                                         \
               build {N   {: GN } CA {: GC} O {: GO} N+ {: GN/+} \
                      CG  {: BCG}                                \
                                                                 \
                      C   {+ {O   CA  N+  } {1.24  32.0    0.0}} \
                      CB  {+ {CA  N   C   } {1.50 109.0 -120.0}} \
                      CD  {+ {CG  N   CA  } {1.50  38.0  172.0}} \
                     }                                           \
               last  {build {N+  {: GN}}}                        \
               print {N CA C O CB CG CD}                         \
             }                                             

     dict set backmap                                            \
         sW  { name  TRP                                         \
               build {N   {: GN } CA {: GC} O {: GO} N+ {: GN/+} \
                      CG  {: BCG} NE1 {: BNE} CZ2 {: BCZ}        \
                      CE3 {: BCE} HE1 {: BPE}                    \
                                                                 \
                      C   {+ {O   CA  N+  } {1.24  32.0    0.0}} \
                      CB  {+ {CA  N   C   } {1.50 109.0 -120.0}} \
                      CD1 {+ {CG  NE1 CZ2 } {1.40  35.6  180.0}} \
                      CD2 {+ {CG  NE1 CZ2 } {1.40  70.0    0.0}} \
                      CE2 {+ {NE1 CG  CE3 } {1.40  70.0    0.0}} \
                      CZ3 {+ {CE3 CZ2 NE1 } {1.40  60.0  180.0}} \
                      CH2 {+ {CZ2 CE3 CG  } {1.40  60.0  180.0}} \
                     }                                           \
               last  {build {N+  {: GN}}}                        \
               print {N CA C O CB CG CD1 CD2 NE1 CE2 CE3 CZ2 CZ3 \
                      CH2 HE1                                    \
                     }                                           \
             }
             # Note: CB can be build from ring coords

     dict set backmap                                            \
         sF  { name  PHE                                         \
               build {N   {: GN } CA {: GC} O {: GO} N+ {: GN/+} \
                      CG  {: BCG} CE1 {: BCE1} CE2 {: BCE2}      \
                                                                 \
                      C   {+ {O   CA  N+  } {1.24  32.0    0.0}} \
                      CB  {+ {CA  N   C   } {1.50 109.0 -120.0}} \
                      CD1 {+ {CG  CE2 CE1 } {1.40  90.0    0.0}} \
                      CD2 {+ {CG  CE1 CE2 } {1.40  90.0    0.0}} \
                      CZ  {+ {CE1 CG  CE2 } {1.40  90.0    0.0}} \
                     }                                           \
               last  {build {N+  {: GN}}}                        \
               print {N CA C O CB CG CD1 CD2 CE1 CE2 CZ}         \
             }
                                                      
     dict set backmap                                            \
         sY  { name  TYR                                         \
               build {N   {: GN } CA {: GC} O {: GO} N+ {: GN/+} \
                      CG  {: BCG} CE1 {: BCE1} CE2 {: BCE2}      \
                                                                 \
                      C   {+ {O   CA  N+  } {1.24  32.0    0.0}} \
                      CB  {+ {CA  N   C   } {1.50 109.0 -120.0}} \
                      CD1 {+ {CG  CE2 CE1 } {1.40  90.0    0.0}} \
                      CD2 {+ {CG  CE1 CE2 } {1.40  90.0    0.0}} \
                      CZ  {+ {CE1 CG  CE2 } {1.40  90.0    0.0}} \
                      OH  {+ {CZ  CE1 CE2 } {1.40 120.0  180.0}} \
                     }                                           \
               last  {build {N+  {: GN}}}                        \
               print {N CA C O CB CG CD1 CD2 CE1 CE2 CZ OH}      \
             }

     dict set backmap                                            \
         sYp { name  PTR                                         \
               build {N   {: GN } CA {: GC} O {: GO} N+ {: GN/+} \
                      CG  {: BCG} CE1 {: BCE1} CE2 {: BCE2}      \
                      P   {: PX }                                \
                                                                 \
                      C   {+ {O   CA  N+  } {1.24  32.0    0.0}} \
                      CB  {+ {CA  N   C   } {1.50 109.0 -120.0}} \
                      CD1 {+ {CG  CE2 CE1 } {1.40  90.0    0.0}} \
                      CD2 {+ {CG  CE1 CE2 } {1.40  90.0    0.0}} \
                      CZ  {+ {CE1 CG  CE2 } {1.40  90.0    0.0}} \
                      OH  {+ {CZ  CE1 CE2 } {1.40 120.0  180.0}} \
                      O1P {+ {P   OH  CZ  } {1.50 109.0   60.0}} \
                      O2P {+ {P   O1P OH  } {1.50 109.0 -120.0}} \
                      O3P {+ {P   O2P O1P } {1.50 109.0  120.0}} \
                     }                                           \
               last  {build {N+  {: GN}}}                        \
               print {N CA C O CB CG CD1 CD2 CE1 CE2 CZ OH       \
                      P O1P O2P O3P}                             \
             }

     dict set backmap                                            \
         sHe { name  HIE                                         \
               build {N   {: GN } CA {: GC} O {: GO} N+ {: GN/+} \
                      CG  {: BCG} ND1 {: BND} NE2 {: BNE}        \
                                                                 \
                      C   {+ {O   CA  N+  } {1.24  32.0    0.0}} \
                      CB  {+ {CA  N   C   } {1.50 109.0 -120.0}} \
                      CD2 {+ {CG  ND1 NE2 } {1.40 109.0    0.0}} \
                      CE1 {+ {ND1 CG  NE2 } {1.40 109.0    0.0}} \
                     }                                           \
               last  {build {N+  {: GN}}}                        \
               print {N CA C O CB CG CD2 CE1 ND1 NE2}            \
             }

     dict set backmap sHd [dict get $backmap sHe]
     dict set backmap sHd  name HID

     dict set backmap                                            \
         sN  { name  ASN                                         \
               build {N   {: GN } CA {: GC} O {: GO} N+ {: GN/+} \
                      CG  {: BCG} OD1 {: BOD} ND2 {: BND}        \
                                                                 \
                      C   {+ {O   CA  N+  } {1.24  32.0    0.0}} \
                      CB  {+ {CA  N   C   } {1.50 109.0 -120.0}} \
                     }                                           \
               last  {build {N+  {: GN}}}                        \
               print {N CA C O CB CG OD1 ND2}                    \
             }

     dict set backmap                                            \
         sQ  { name  GLN                                         \
               build {N   {: GN } CA {: GC} O {: GO} N+ {: GN/+} \
                      CD  {: BCD} OE1 {: BOD} NE2 {: BND}        \
                                                                 \
                      C   {+ {O   CA  N+  } {1.24  32.0    0.0}} \
                      CB  {+ {CA  N   C   } {1.50 109.0 -120.0}} \
                      CG  {+ {CB  CA  CD  } {1.50 109.0   30.0}} \
                     }                                           \
               last  {build {N+  {: GN}}}                        \
               print {N CA C O CB CG CD OE1 NE2}                 \
             }

     dict set backmap                                            \
         sS  { name  SER                                         \
               build {N   {: GN } CA {: GC} O {: GO} N+ {: GN/+} \
                      OG  {: BOG} HG {: BPG}                     \
                                                                 \
                      C   {+ {O   CA  N+  } {1.24  32.0    0.0}} \
                      CB  {+ {CA  N   C   } {1.50 109.0 -120.0}} \
                     }                                           \
               last  {build {N+  {: GN}}}                        \
               print {N CA C O CB OG HG}                         \
             }

     dict set backmap                                            \
         sSp { name  SEP                                         \
               build {N   {: GN } CA {: GC} O {: GO} N+ {: GN/+} \
                      CB  {: BCB} P {: PX}                       \
                                                                 \
                      C   {+ {O   CA  N+  } {1.24  32.0    0.0}} \
                      OG  {+ {CB  CA  P   } {1.50 109.0    0.0}} \
                      O1P {+ {P   OG  CB  } {1.50 109.0   60.0}} \
                      O2P {+ {P   O1P OG  } {1.50 109.0 -120.0}} \
                      O3P {+ {P   O2P O1P } {1.50 109.0  120.0}} \
                     }                                           \
               last  {build {N+  {: GN}}}                        \
               print {N CA C O CB OG P O1P O2P O3P}              \
             }
             # Note: diedral OG1-CB-CA-P provides the smallest
             # dispertion (+/- 35º) compared to others, e.g.:
             # OG1-P-CA-CB (+/- 75º), when considering steric
             # clashes.

     dict set backmap                                            \
         sT  { name  THR                                         \
               build {N   {: GN } CA {: GC} O {: GO} N+ {: GN/+} \
                      OG1 {: BOG} HG1 {: BPG}                    \
                                                                 \
                      C   {+ {O   CA  N+  } {1.24  32.0    0.0}} \
                      CB  {+ {CA  N   C   } {1.50 109.0 -120.0}} \
                      CG2 {+ {CB  CA  OG1 } {1.50 109.0 -120.0}} \
                     }                                           \
               last  {build {N+  {: GN}}}                        \
               print {N CA C O CB OG1 CG2 HG1}                   \
             }

     dict set backmap                                            \
         sTp { name  TPO                                         \
               build {N   {: GN } CA {: GC} O {: GO} N+ {: GN/+} \
                      CB  {: BCB} P {: PX}                       \
                                                                 \
                      C   {+ {O   CA  N+  } {1.24  32.0    0.0}} \
                      OG1 {+ {CB  CA  P   } {1.50 109.0  -30.0}} \
                      CG2 {+ {CB  CA  OG1 } {1.50 109.0 -120.0}} \
                      O1P {+ {P   OG1 CB  } {1.50 109.0   60.0}} \
                      O2P {+ {P   O1P OG1 } {1.50 109.0 -120.0}} \
                      O3P {+ {P   O2P O1P } {1.50 109.0  120.0}} \
                     }                                           \
               last  {build {N+  {: GN}}}                        \
               print {N CA C O CB OG1 P O1P O2P O3P}             \
             } 
             # Note: diedral OG1-CB-CA-P provides the smallest
             # dispertion (+/- 35º) compared to others, e.g.:
             # OG1-P-CA-CB (+/- 75º), when considering steric
             # clashes. The actual value takes into account the
             # presence of the methyl group.

     dict set backmap                                            \
         sC  { name  CYS                                         \
               build {N   {: GN } CA {: GC} O {: GO} N+ {: GN/+} \
                      SG  {: BSG} HG {: BPG}                     \
                                                                 \
                      C   {+ {O   CA  N+  } {1.24  32.0    0.0}} \
                      CB  {+ {CA  N   C   } {1.50 109.0 -120.0}} \
                     }                                           \
               last  {build {N+  {: GN}}}                        \
               print {N CA C O CB SG HG}                         \
                                                                 \
               bridge {BSG {sX {BSG {SG SG sX}}                  \
                            sC {BSG {SG SG sX}}                  \
                      }}                                         \
             } ;   #   at1  r2  at2  n1 n2 r2
             # Bugfix to CYX residue named CYS (e.g PDB: 1ETM)

     dict set backmap                                            \
         sZ  { name  CYM                                         \
               build {N   {: GN } CA {: GC} O {: GO} N+ {: GN/+} \
                      SG  {: BSG}                                \
                                                                 \
                      C   {+ {O   CA  N+  } {1.24  32.0    0.0}} \
                      CB  {+ {CA  N   C   } {1.50 109.0 -120.0}} \
                     }                                           \
               last  {build {N+  {: GN}}}                        \
               print {N CA C O CB SG}                            \
             }

     dict set backmap                                            \
         sX  { name  CYX                                         \
               build {N   {: GN } CA {: GC} O {: GO} N+ {: GN/+} \
                      SG  {: BSG}                                \
                                                                 \
                      C   {+ {O   CA  N+  } {1.24  32.0    0.0}} \
                      CB  {+ {CA  N   C   } {1.50 109.0 -120.0}} \
                     }                                           \
               last  {build {N+  {: GN}}}                        \
               print {N CA C O CB SG}                            \
                                                                 \
               bridge {BSG {sX {BSG {SG SG sX}}                  \
                            sC {BSG {SG SG sX}}                  \
                      }}                                         \
             }

     dict set backmap sCp [dict get $backmap sC]
     dict set backmap sCp  build HG {: BE1}

     dict set backmap                                            \
         sM  { name  MET                                         \
               build {N   {: GN } CA {: GC} O {: GO} N+ {: GN/+} \
                      SD  {: BSD}                                \
                                                                 \
                      C   {+ {O   CA  N+  } {1.24  32.0    0.0}} \
                      CB  {+ {CA  N   C   } {1.50 109.0 -120.0}} \
                      CG  {+ {SD  CB  CA  } {1.80  32.0 -130.0}} \
                      CE  {+ {SD  CG  CB  } {1.80 109.0   60.0}} \
                     }                                           \
               last  {build {N+  {: GN}}}                        \
               print {N CA C O CB CG SD CE}                      \
             }

     dict set backmap                                            \
         sD  { name  ASP                                         \
               build {N   {: GN } CA {: GC} O {: GO} N+ {: GN/+} \
                      CG  {: BCG} OD1 {: BOE1} OD2 {: BOE2}      \
                                                                 \
                      C   {+ {O   CA  N+  } {1.24  32.0    0.0}} \
                      CB  {+ {CA  N   C   } {1.50 109.0 -120.0}} \
                     }                                           \
               last  {build {N+  {: GN}}}                        \
               print {N CA C O CB CG OD1 OD2}                    \
             }

     dict set backmap sDh [dict get $backmap sD]
     dict set backmap sDh  name ASH

     dict set backmap                                            \
         sE  { name  GLU                                         \
               build {N   {: GN } CA {: GC} O {: GO} N+ {: GN/+} \
                      CD  {: BCD} OE1 {: BOE1} OE2 {: BOE2}      \
                                                                 \
                      C   {+ {O   CA  N+  } {1.24  32.0    0.0}} \
                      CB  {+ {CA  N   C   } {1.50 109.0 -120.0}} \
                      CG  {+ {CB  CA  CD  } {1.50 109.0   30.0}} \
                     }                                           \
               last  {build {N+  {: GN}}}                        \
               print {N CA C O CB CG CD OE1 OE2}                 \
             }

     dict set backmap sEh [dict get $backmap sE]
     dict set backmap sEh  name GLH

     dict set backmap                                            \
         sK  { name  LYS                                         \
               build {N   {: GN } CA {: GC} O {: GO} N+ {: GN/+} \
                      CG  {: BCG} CE {: BCE}                     \
                                                                 \
                      C   {+ {O   CA  N+  } {1.24  32.0    0.0}} \
                      CB  {+ {CA  N   C   } {1.50 109.0 -120.0}} \
                      CD  {+ {CG  CE  CB  } {1.50  32.0    0.0}} \
                      NZ  {+ {CE  CD  CG  } {1.50 109.0  180.0}} \
                     }                                           \
               last  {build {N+  {: GN}}}                        \
               print {N CA C O CB CG CD CE NZ}                   \
             }

     dict set backmap sKm [dict get $backmap sK]

     dict set backmap sKa [dict get $backmap sK]
     dict set backmap sKa  build CE {: GNac}

     dict set backmap                                            \
         sR  { name  ARG                                         \
               build {N   {: GN } CA {: GC} O {: GO} N+ {: GN/+} \
                      CG  {: BCG}                                \
                      CZ  {: BCZ} NH1 {: BNN1} NH2 {: BNN2}      \
                                                                 \
                      C   {+ {O   CA  N+  } {1.24  32.0    0.0}} \
                      CB  {+ {CA  N   C   } {1.50 109.0 -120.0}} \
                      CD  {+ {CG  CB  CZ  } {1.50 109.0   10.0}} \
                      NE  {+ {CD  CG  CZ  } {1.50 109.0   10.0}} \
                     }                                           \
               last  {build {N+  {: GN}}}                        \
               print {N CA C O CB CG CD NE CZ NH1 NH2}           \
             }                                                   \

   # AMBER terminal residues
     foreach aa {G A R N D Dh C Z X Cp Q E Eh He Hd I L K Km Ka M F P S Sp T Tp W Y Yp V} {
          
         dict set backmap  n$aa [dict get $backmap s$aa] ;# Charged Nt
         dict set backmap  c$aa [dict get $backmap s$aa] ;# Charged Ct
         dict set backmap  a$aa [dict get $backmap s$aa] ;# Neutral Nt (~acetylated)
         dict set backmap  m$aa [dict get $backmap s$aa] ;# Neutral Ct (~amidated)
     }

   # Nucleic Acids
   # High presition for A/B/Z DNA forms, PDB: 1BNA 4IZQ 4OCB
     dict set backmap                                            \
         DTX { name  DT                                          \
               build {P   {: PX } C5' {: C5X} C1' {: O3' C1X}    \
                      X3  {: O3'/- C1X/- }                       \
                      O2  {: O2T} N3  {: N3T} O4  {: O4T}        \
                                                                 \
                      N1  {+ {C1' O2  O4  } {1.45  56.0    0.0}} \
                      C2  {+ {O2  N1  N3  } {1.22  30.0    0.0}} \
                      C4  {+ {O4  N3  C2  } {1.22  30.0    0.0}} \
                      C6  {+ {N1  C1' C2  } {1.40 120.0  180.0}} \
                      C5  {+ {C6  N1  C1' } {1.40 120.0  180.0}} \
                      C7  {+ {C5  C4  O4  } {1.50 120.0    0.0}} \
                                                                 \
                      O4' {+ {C1' N1  C5' } {1.40 109.0   30.0}} \
                      C2' {+ {C1' N1  O4' } {1.50 109.0 -115.0}} \
                      C4' {+ {C2' C1' C5' } {2.40  66.0   16.0}} \
                      C3' {+ {C4' O4' C5' } {1.50 109.0 -120.0}} \
                      O3' {+ {C3' C4' C2' } {1.40 109.0 -120.0}} \
                      O5' {+ {P   C5' C4' } {1.50  30.0   16.0}} \
                      OP2 {+ {P   O5' X3  } {1.50 109.0  -90.0}} \
                      OP1 {+ {P   OP2 O5' } {1.50 109.0  120.0}} \
                     }                                           \
               first {apply TX5}                                 \
               last  {apply TX3}                                 \
               print {P OP1 OP2 O5' C5' C4' O4' C3' O3' C2' C1'  \
                      N1 C2 O2 N3 C4 O4 C5 C6 C7}                \
              }

     dict set backmap                                            \
         DCX { name  DC                                          \
               build {P   {: PX } C5' {: C5X} C1' {: O3' C1X}    \
                      X3  {: O3'/- C1X/- }                       \
                      O2  {: O2C} N3  {: N3C} N4  {: N4C}        \
                                                                 \
                      N1  {+ {C1' O2  N4  } {1.45  56.0    0.0}} \
                      C2  {+ {O2  N1  N3  } {1.22  30.0    0.0}} \
                      C4  {+ {N4  N3  C2  } {1.22  30.0    0.0}} \
                      C6  {+ {N1  C1' C2  } {1.40 120.0  180.0}} \
                      C5  {+ {C6  N1  C1' } {1.40 120.0  180.0}} \
                                                                 \
                      O4' {+ {C1' N1  C5' } {1.40 109.0   30.0}} \
                      C2' {+ {C1' N1  O4' } {1.50 109.0 -115.0}} \
                      C4' {+ {C2' C1' C5' } {2.40  66.0   16.0}} \
                      C3' {+ {C4' O4' C5' } {1.50 109.0 -120.0}} \
                      O3' {+ {C3' C4' C2' } {1.40 109.0 -120.0}} \
                      O5' {+ {P   C5' C4' } {1.50  30.0   16.0}} \
                      OP2 {+ {P   O5' X3  } {1.50 109.0  -90.0}} \
                      OP1 {+ {P   OP2 O5' } {1.50 109.0  120.0}} \
                     }                                           \
               first {apply CX5}                                 \
               last  {apply CX3}                                 \
               print {P OP1 OP2 O5' C5' C4' O4' C3' O3' C2' C1'  \
                      N1 C2 O2 N3 C4 N4 C5 C6}                   \
             }

     dict set backmap                                            \
         DGX { name  DG                                          \
               build {P   {: PX } C5' {: C5X} C1' {: O3' C1X}    \
                      X3  {: O3'/- C1X/- }                       \
                      N2  {: N2G} N1  {: N1G} O6  {: O6G}        \
                                                                 \
                      C2  {+ {N1  N2  C1' } {1.40  30.0    0.0}} \
                      N3  {+ {C2  N1  N2  } {1.40 120.0  180.0}} \
                      C4  {+ {N3  C2  N1  } {1.40 120.0    0.0}} \
                      C5  {+ {C4  N3  C2  } {1.40 120.0    0.0}} \
                      C6  {+ {N1  C2  N3  } {1.40 120.0    0.0}} \
                      N7  {+ {C5  C4  N3  } {1.40 109.0  180.0}} \
                      C8  {+ {N7  C5  C4  } {1.40 109.0    0.0}} \
                      N9  {+ {C4  C5  N7  } {1.40 109.0    0.0}} \
                                                                 \
                      O4' {+ {C1' N9  C5' } {1.40 109.0   30.0}} \
                      C2' {+ {C1' N9  O4' } {1.50 109.0 -115.0}} \
                      C4' {+ {C2' C1' C5' } {2.40  66.0   16.0}} \
                      C3' {+ {C4' O4' C5' } {1.50 109.0 -120.0}} \
                      O3' {+ {C3' C4' C2' } {1.40 109.0 -120.0}} \
                      O5' {+ {P   C5' C4' } {1.50  30.0   16.0}} \
                      OP2 {+ {P   O5' X3  } {1.50 109.0  -90.0}} \
                      OP1 {+ {P   OP2 O5' } {1.50 109.0  120.0}} \
                     }                                           \
               first {apply GX5}                                 \
               last  {apply GX3}                                 \
               print {P OP1 OP2 O5' C5' C4' O4' C3' O3' C2' C1'  \
                      N9 C8 N7 C5 C6 O6 N1 C2 N2 N3 C4}          \
             }
  
     dict set backmap                                            \
         DAX { name  DA                                          \
               build {P   {: PX } C5' {: C5X} C1' {: O3' C1X}    \
                      X3  {: O3'/- C1X/- }                       \
                      H2  {: C2A} N1  {: N1A} N6 {: N6A}         \
                                                                 \
                      C2  {+ {N1  N6  C1' } {1.40 150.0    0.0}} \
                      N3  {+ {C2  N1  N6  } {1.40 120.0    0.0}} \
                      C4  {+ {N3  C2  N1  } {1.40 120.0    0.0}} \
                      C5  {+ {C4  N3  C2  } {1.40 120.0    0.0}} \
                      C6  {+ {N1  C2  N3  } {1.40 120.0    0.0}} \
                      N7  {+ {C5  C4  N3  } {1.40 109.0  180.0}} \
                      C8  {+ {N7  C5  C4  } {1.40 109.0    0.0}} \
                      N9  {+ {C4  C5  N7  } {1.40 109.0    0.0}} \
                                                                 \
                      O4' {+ {C1' N9  C5' } {1.40 109.0   30.0}} \
                      C2' {+ {C1' N9  O4' } {1.50 109.0 -115.0}} \
                      C4' {+ {C2' C1' C5' } {2.40  66.0   16.0}} \
                      C3' {+ {C4' O4' C5' } {1.50 109.0 -120.0}} \
                      O3' {+ {C3' C4' C2' } {1.40 109.0 -120.0}} \
                      O5' {+ {P   C5' C4' } {1.50  30.0   16.0}} \
                      OP2 {+ {P   O5' X3  } {1.50 109.0  -90.0}} \
                      OP1 {+ {P   OP2 O5' } {1.50 109.0  120.0}} \
                     }                                           \
               first {apply AX5}                                 \
               last  {apply AX3}                                 \
               print {P OP1 OP2 O5' C5' C4' O4' C3' O3' C2' C1'  \
                      N9 C8 N7 C5 C6 N6 N1 C2 H2 N3 C4}          \
             }

   # 5' END
     dict set backmap                                            \
         TX5 { name  DT5                                         \
               build {C5' {: C5X} C1' {: O3' C1X}                \
                      O2  {: O2T} N3  {: N3T} O4 {: O4T}         \
                                                                 \
                      N1  {+ {C1' O2  O4  } {1.45  56.0    0.0}} \
                      C2  {+ {O2  N1  N3  } {1.22  30.0    0.0}} \
                      C4  {+ {O4  N3  C2  } {1.22  30.0    0.0}} \
                      C6  {+ {N1  C1' C2  } {1.40 120.0  180.0}} \
                      C5  {+ {C6  N1  C1' } {1.40 120.0  180.0}} \
                      C7  {+ {C5  C4  O4  } {1.50 120.0    0.0}} \
                                                                 \
                      O4' {+ {C1' N1  C5' } {1.40 109.0   30.0}} \
                      C2' {+ {C1' N1  O4' } {1.50 109.0 -115.0}} \
                      C4' {+ {C2' C1' C5' } {2.40  66.0   16.0}} \
                      C3' {+ {C4' O4' C5' } {1.50 109.0 -120.0}} \
                      O3' {+ {C3' C4' C2' } {1.40 109.0 -120.0}} \
                      O5' {+ {C5' C4' O4' } {1.40 109.0  -80.0}} \
                     }                                           \
               print {O5' C5' C4' O4' C3' O3' C2' C1'            \
                      N1 C2 O2 N3 C4 O4 C5 C6 C7}                \
             }

     dict set backmap                                            \
         CX5 { name  DC5                                         \
               build {C5' {: C5X} C1' {: O3' C1X}                \
                      O2  {: O2C} N3  {: N3C} N4 {: N4C}         \
                                                                 \
                      N1  {+ {C1' O2  N4  } {1.45  56.0    0.0}} \
                      C2  {+ {O2  N1  N3  } {1.22  30.0    0.0}} \
                      C4  {+ {N4  N3  C2  } {1.22  30.0    0.0}} \
                      C6  {+ {N1  C1' C2  } {1.40 120.0  180.0}} \
                      C5  {+ {C6  N1  C1' } {1.40 120.0  180.0}} \
                                                                 \
                      O4' {+ {C1' N1  C5' } {1.40 109.0   30.0}} \
                      C2' {+ {C1' N1  O4' } {1.50 109.0 -115.0}} \
                      C4' {+ {C2' C1' C5' } {2.40  66.0   16.0}} \
                      C3' {+ {C4' O4' C5' } {1.50 109.0 -120.0}} \
                      O3' {+ {C3' C4' C2' } {1.40 109.0 -120.0}} \
                      O5' {+ {C5' C4' O4' } {1.40 109.0  -80.0}} \
                     }                                           \
               print {O5' C5' C4' O4' C3' O3' C2' C1'            \
                      N1 C2 O2 N3 C4 N4 C5 C6}                   \
             }

     dict set backmap                                            \
         GX5 { name  DG5                                         \
               build {C5' {: C5X} C1' {: O3' C1X}                \
                      N2  {: N2G} N1  {: N1G} O6 {: O6G}         \
                                                                 \
                      C2  {+ {N1  N2  C1' } {1.40  30.0    0.0}} \
                      N3  {+ {C2  N1  N2  } {1.40 120.0  180.0}} \
                      C4  {+ {N3  C2  N1  } {1.40 120.0    0.0}} \
                      C5  {+ {C4  N3  C2  } {1.40 120.0    0.0}} \
                      C6  {+ {N1  C2  N3  } {1.40 120.0    0.0}} \
                      N7  {+ {C5  C4  N3  } {1.40 109.0  180.0}} \
                      C8  {+ {N7  C5  C4  } {1.40 109.0    0.0}} \
                      N9  {+ {C4  C5  N7  } {1.40 109.0    0.0}} \
                                                                 \
                      O4' {+ {C1' N9  C5' } {1.40 109.0   30.0}} \
                      C2' {+ {C1' N9  O4' } {1.50 109.0 -115.0}} \
                      C4' {+ {C2' C1' C5' } {2.40  66.0   16.0}} \
                      C3' {+ {C4' O4' C5' } {1.50 109.0 -120.0}} \
                      O3' {+ {C3' C4' C2' } {1.40 109.0 -120.0}} \
                      O5' {+ {C5' C4' O4' } {1.40 109.0  -80.0}} \
                     }                                           \
               print {O5' C5' C4' O4' C3' O3' C2' C1'            \
                      N9 C8 N7 C5 C6 O6 N1 C2 N2 N3 C4}          \
             }

     dict set backmap                                            \
         AX5 { name  DA5                                         \
               build {C5' {: C5X} C1' {: O3' C1X}                \
                      H2  {: C2A} N1  {: N1A} N6 {: N6A}         \
                                                                 \
                      C2  {+ {N1  N6  C1' } {1.40 150.0    0.0}} \
                      N3  {+ {C2  N1  N6  } {1.40 120.0    0.0}} \
                      C4  {+ {N3  C2  N1  } {1.40 120.0    0.0}} \
                      C5  {+ {C4  N3  C2  } {1.40 120.0    0.0}} \
                      C6  {+ {N1  C2  N3  } {1.40 120.0    0.0}} \
                      N7  {+ {C5  C4  N3  } {1.40 109.0  180.0}} \
                      C8  {+ {N7  C5  C4  } {1.40 109.0    0.0}} \
                      N9  {+ {C4  C5  N7  } {1.40 109.0    0.0}} \
                                                                 \
                      O4' {+ {C1' N9  C5' } {1.40 109.0   30.0}} \
                      C2' {+ {C1' N9  O4' } {1.50 109.0 -115.0}} \
                      C4' {+ {C2' C1' C5' } {2.40  66.0   16.0}} \
                      C3' {+ {C4' O4' C5' } {1.50 109.0 -120.0}} \
                      O3' {+ {C3' C4' C2' } {1.40 109.0 -120.0}} \
                      O5' {+ {C5' C4' O4' } {1.40 109.0  -80.0}} \
                     }                                           \
               print {O5' C5' C4' O4' C3' O3' C2' C1'            \
                      N9 C8 N7 C5 C6 N6 N1 C2 H2 N3 C4}          \
             }

   # Alternative implementation
   # dict   set backmap TX5 [dict get $backmap DTX]
   # dict   set backmap TX5 build O5' {+ {C5' C4' O4'} {1.40 109.0 -80.0}}
   # dict unset backmap TX5 build P
   # dict unset backmap TX5 build X3
   # dict unset backmap TX5 build OP1
   # dict unset backmap TX5 build OP2
   # dict   set backmap TX5 print {O5' C5' C4' O4' C3' O3' C2' C1' N1 C2 O2 N3 C4 O4 C5 C6 C7}

   # 5' END closed circular DNA
     dict set backmap TW5 [dict get $backmap DTX]

     dict set backmap CW5 [dict get $backmap DCX]

     dict set backmap GW5 [dict get $backmap DGX]

     dict set backmap AW5 [dict get $backmap DAX]

     dict set backmap                                            \
         TW5 bridge {PX { DGX {C1X {P O3' TW5 w}}                \
                          DCX {C1X {P O3' TW5 w}}                \
                          DAX {C1X {P O3' TW5 w}}                \
                          DTX {C1X {P O3' TW5 w}}                \
                          GW3 {C1X {P O3' TW5 w}}                \
                          CW3 {C1X {P O3' TW5 w}}                \
                          AW3 {C1X {P O3' TW5 w}}                \
                          TW3 {C1X {P O3' TW5 w}}                \
                    }}

     dict set backmap                                            \
         CW5 bridge {PX { DGX {C1X {P O3' CW5 w}}                \
                          DCX {C1X {P O3' CW5 w}}                \
                          DAX {C1X {P O3' CW5 w}}                \
                          DTX {C1X {P O3' CW5 w}}                \
                          GW3 {C1X {P O3' CW5 w}}                \
                          CW3 {C1X {P O3' CW5 w}}                \
                          AW3 {C1X {P O3' CW5 w}}                \
                          TW3 {C1X {P O3' CW5 w}}                \
                    }}

     dict set backmap                                            \
         GW5 bridge {PX { DGX {C1X {P O3' GW5 w}}                \
                          DCX {C1X {P O3' GW5 w}}                \
                          DAX {C1X {P O3' GW5 w}}                \
                          DTX {C1X {P O3' GW5 w}}                \
                          GW3 {C1X {P O3' GW5 w}}                \
                          CW3 {C1X {P O3' GW5 w}}                \
                          AW3 {C1X {P O3' GW5 w}}                \
                          TW3 {C1X {P O3' GW5 w}}                \
                    }}

     dict set backmap                                            \
         AW5 bridge {PX { DGX {C1X {P O3' AW5 w}}                \
                          DCX {C1X {P O3' AW5 w}}                \
                          DAX {C1X {P O3' AW5 w}}                \
                          DTX {C1X {P O3' AW5 w}}                \
                          GW3 {C1X {P O3' AW5 w}}                \
                          CW3 {C1X {P O3' AW5 w}}                \
                          AW3 {C1X {P O3' AW5 w}}                \
                          TW3 {C1X {P O3' AW5 w}}                \
                    }}

   # 3' END
     dict set backmap TX3 [dict get $backmap DTX]
     dict set backmap TX3 name DT3
     
     dict set backmap CX3 [dict get $backmap DCX]
     dict set backmap CX3 name DC3
     
     dict set backmap GX3 [dict get $backmap DGX]
     dict set backmap GX3 name DG3

     dict set backmap AX3 [dict get $backmap DAX]
     dict set backmap AX3 name DA3

   # 3' END closed circular DNA
     dict set backmap TW3 [dict get $backmap DTX]

     dict set backmap CW3 [dict get $backmap DCX]

     dict set backmap GW3 [dict get $backmap DGX]

     dict set backmap AW3 [dict get $backmap DAX]

   # Membrane
     dict set backmap                                            \
         xPC { nomap * }

     dict set backmap                                            \
         xPE { nomap * }

     dict set backmap                                            \
         xPS { nomap * }

     dict set backmap                                            \
         xMY { nomap * }

     dict set backmap                                            \
         xPA { nomap * }

     dict set backmap                                            \
         xOL { nomap * }

     dict set backmap                                            \
         CMM { nomap * }

     dict set backmap                                            \
         CPP { nomap * }

     dict set backmap                                            \
         CPO { nomap * }

     dict set backmap                                            \
         EPO { nomap * }

     dict set backmap                                            \
         SPO { nomap * }

   # Solvent + Ions
     dict set backmap                                            \
         WT4 { nomap * }
     
     dict set backmap                                            \
         WLS { nomap * }
         
     dict set backmap                                            \
         ClW { nomap * }
     
     dict set backmap                                            \
         NaW { nomap * }
     
     dict set backmap                                            \
         KW  { nomap * }

     dict set backmap                                            \
         MgX { name   MG                                         \
               build {MG {: MgX}}                                \
               print {MG}                                        \
             }

     dict set backmap                                            \
         CaX { name   CA                                         \
               build {CA {: CaX}}                                \
               print {CA}                                        \
             }

     dict set backmap                                            \
         ZnX { name   ZN                                         \
               build {ZN {: ZnX}}                                \
               print {ZN}                                        \
             }

   # Atomistic residue exceptions
     dict set backmap                                            \
         CYS { bridge {SG {CYS {SG {SG SG CYX}}                  \
                           CYX {SG {SG SG CYX}}                  \
                      }}                                         \
             }

     dict set backmap                                            \
         CYX { bridge {SG {CYS {SG {SG SG CYX}}                  \
                           CYX {SG {SG SG CYX}}                  \
                      }}                                         \
             }

     dict set backmap                                            \
         WAT { nomap * }

     dict set backmap                                            \
         SOL { nomap * }

     dict set backmap                                            \
         KW  { nomap * }

     dict set backmap                                            \
         NA  { nomap * }

     dict set backmap                                            \
         CL  { nomap * }

   # Done!
     return ;# Avoids printing actions to interactive console
}


proc sirah_build {res resdb xyz} {
   # Load database
     global backmap

     set name  [dict get $resdb $res name] ;# residue name
     set build {}                          ;# building instructions
     set atoms {}                          ;# output list

   # Apply new residue mapping?
     if {![dict exists $resdb $res res-] && [dict exists $backmap $name first apply]} { set name  [dict get $backmap $name first apply] }
     if {![dict exists $resdb $res res+] && [dict exists $backmap $name last  apply]} { set name  [dict get $backmap $name last  apply] }

   # Setup building instructions
     if {                                   [dict exists $backmap $name       build]} { set build                    [dict get $backmap $name       build]  }     
     if {![dict exists $resdb $res res-] && [dict exists $backmap $name first build]} { set build [dict merge $build [dict get $backmap $name first build]] }
     if {![dict exists $resdb $res res+] && [dict exists $backmap $name last  build]} { set build [dict merge $build [dict get $backmap $name last  build]] }

   # No backmapping procedure
     if {![llength $build]} {

         foreach {atom index} [dict get $resdb $res atoms] { 

              lappend atoms [list $atom $name [lindex $xyz $index]]
         }

         return $atoms
     }

   # Build!
     foreach map [dict keys $build] {

        if { [info exists new($map)] } { continue } ;# If atom already exists then use it!

        set info [dict get $build $map] ;# Building instructions
        
      # Function set <:>
        if { [lindex $info 0] == ":" } {
            
            foreach atom [lrange $info 1 end] {

                  # catch ATOM/[+-]
                    if     { [string match {?*/-} $atom] } { set use [dict get $resdb $res res-] }\
                    elseif { [string match {?*/+} $atom] } { set use [dict get $resdb $res res+] }\
                    else   { set use $res }

                    set atom [string trim $atom {*/[+-]}]

                  # Create new atom
                    if {[dict exists $resdb $use atoms $atom]} {
                    
                       set new($map) [lindex $xyz [dict get $resdb $use atoms $atom]]
                       
                       break
                    }
            }
            
            if {![info exists new($map)]} { puts stderr "Error! Can't set atom $map in $name"; return }

      # Function add <+>
        } elseif { [lindex $info 0] == "+" } {
            
            foreach atom [lindex $info 1] { if {![info exists new($atom)]} { puts stderr "Error! First set atom $atom in $name"; return } }

            set new($map) [addatom \
                                   $new([lindex $info 1 0]) \
                                   $new([lindex $info 1 1]) \
                                   $new([lindex $info 1 2]) \
                                        [lindex $info 2 0]  \
                                        [lindex $info 2 1]  \
                                        [lindex $info 2 2]  \
                          ]
            
      # Function unknown
        } else { puts stderr "Error! Unknown building function for atom $map in $name"; return }
     }

   # Printable atoms >>>> Add functionality split??? e.g.: print { HEM{} Fe{} }
     foreach atom [dict get $backmap $name print] { lappend atoms [list $atom [dict get $backmap $name name] $new($atom)] }

     return $atoms
}


proc sirah_backmap {args} {
   # Read arguments --------------------------------------------------------------------------------------------
     array set opts [ getopts $args switch { now nomin cuda gbsa noload help } option { mol first last each frames outname mpi maxcyc ncyc cutoff} ]

     if { [info exists opts()]}            { puts stderr "Error! Bad argument: [join $opts()]"; return }
     
     if { [info exists opts(help)]}        { sirah_help sirah_backmap; return }
     
     if {![info exists opts(nomin)]}       { set opts(nomin)  0 }
     
     if {![info exists opts(noload)]}      { set opts(noload) 0 }

     if { [info exists opts(mol)] && $opts(mol) != "top" } { set molid $opts(mol) } else { set molid [molinfo top] }

     if {![info exists opts(outname)]}     { set opts(outname) "backmap" }

     if {![info exists opts(maxcyc)]}      { set opts(maxcyc) 150 }

     if {![info exists opts(ncyc)]}        { set opts(ncyc)   100 }
     
     if {![info exists opts(cutoff)]}      { set opts(cutoff) 12 }

     if {![info exists opts(gbsa)]}        { set opts(gbsa)   0 }

     if { [info exists opts(mpi)] && $opts(mpi) > 1} { set exe "sander.MPI"; set MPI "mpirun -np $opts(mpi)"  } else\
                                                     { set exe "sander"    ; set MPI "" ;    set  opts(mpi) 0 }
     
     if { [info exists opts(cuda)]}        { set exe "pmemd.cuda"; set opts(gbsa) 1; set opts(cutoff) 999; set MPI "" ; set opts(mpi) 0 } 
                                           # pmemd does not explicitly support vacuum simulations [http://archive.ambermd.org/201206/0575.html]

     if { [info exists opts(now)]}         { set opts(frames) [molinfo $molid get frame] }

     if {![info exists opts(first)]}       { set opts(first)  0 }
     
     if {![info exists opts(last)]}        { set opts(last)   [expr [molinfo $molid get numframes] -1] }

     if {![info exists opts(each)]}        { set opts(each)   1 }

     if {![info exists opts(frames)]}      { set opts(frames) [seq $opts(first) $opts(last) $opts(each)] }
   #------------------------------------------------------------------------------------------------------------

     if {! $opts(nomin) } {

      # Check required packages
        puts -nonewline "Checking for external packages... "
        global env
        if {![info exists  env(AMBERHOME)]}                                                             { puts stderr "AMBERTOOL is not available, check \$AMBERHOME path" ; return }
        if { [file exists $env(AMBERHOME)/bin/tleap]}   { set tleap  $env(AMBERHOME)/bin/tleap   } else { puts stderr "LEAP is not available, check \$AMBERHOME path"      ; return }
        if { [file exists $env(AMBERHOME)/bin/cpptraj]} { set ptraj  $env(AMBERHOME)/bin/cpptraj } else { puts stderr "CPPTRAJ is not available, check \$AMBERHOME path"   ; return }
        if { [file exists $env(AMBERHOME)/bin/$exe]}    { set sander $env(AMBERHOME)/bin/$exe    } else { puts stderr "$exe is not available, check \$AMBERHOME path"      ; return }
   
      # MPI
        if { $opts(mpi) && ![file exists [auto_execok mpirun]] } { puts stderr "mpirun is not available in your PATH" ; return }
    

      # Check force field AMBERTOOLS 14-18
      # Protein | DNA
        if     { [file exists $env(AMBERHOME)/dat/leap/cmd/leaprc.ff14SB]       } { lappend forcefield(cmd)       leaprc.ff14SB }\
        elseif { [file exists $env(AMBERHOME)/dat/leap/cmd/oldff/leaprc.ff14SB] } { lappend forcefield(cmd) oldff/leaprc.ff14SB }\
        else   {  puts stderr "Can't find leaprc.ff14SB" ; return }

      # Mg2+/Ca2+
        if     { [file exists $env(AMBERHOME)/dat/leap/parm/frcmod.ionslrcm_cm_tip3p]   } { lappend forcefield(parm) frcmod.ionslrcm_cm_tip3p   }\
        elseif { [file exists $env(AMBERHOME)/dat/leap/parm/frcmod.ions234lm_126_tip3p] } { lappend forcefield(parm) frcmod.ions234lm_126_tip3p }\
        else   {  puts stderr "Can't find ion parameters" ; return }

      # PTM: phosphorylated amino acid
        lappend forcefield(cmd) leaprc.phosaa10

      # Done!
        puts "OK"
     }

   # Selections
     set sys      [atomselect $molid "all"]
     set atm2res  [$sys get residue] ;# atom info
     set atm2resn [$sys get resname]
     set atm2name [$sys get name   ]
     
     set noh      [atomselect $molid "noh"] ;# skip H atoms to avoid nomenclature problems
     
   # AMBER patch for GROMACS proteins | I need to improve this
     set CD       [atomselect $molid "resname ILE and name CD"]
     set OC1      [atomselect $molid "name OC1"]
     set OC2      [atomselect $molid "name OC2"]
     
     $CD  set name CD1
     $OC1 set name OXT
     $OC2 set name O

   # Databases
     global backmap             ;# Maps
     set resdb    [dict create] ;# Residues
     
   # Set new residue index
     set res(prev) -1     ;# Previous residue index
     set res(new)   0     ;# New residue index
     
     puts -nonewline "Building residue database... "
     foreach atom [$noh get {residue resname chain resid name index}] bonds [$noh getbonds] {
     
           # Check Backmapping exists
             if {[dict exists $backmap [lindex $atom 1] nomap]} { set nomap([lindex $atom 1]) 1; continue }

             set r1 [lindex $atom 0]; # residue index

           # Build residue database
           # resdb --> index --> name:     ()
           #                 --> chain:    () A,B...
           #                 --> resid:    ()
           #                 --> reindex:  ()
           #                 --> frag:     () 1,2...
           #                 --> atoms --> GN --> index
           #                           --> *  --> ...  
           #                 -->[res-:     ()]
           #                 -->[res+:     ()]
           #       --> ...
             
           # Add residue info
             if {$r1 != $res(prev)} {
             
             #  New fragment
                if {![dict exists $resdb $res(prev) res+] || [dict get $resdb $res(prev) res+] != $r1 } {incr frag}
                
             #  Update info on new residue
                set  res(prev) $r1; incr res(new)
             
                dict set resdb $r1 name     [lindex $atom 1]
                dict set resdb $r1 chain    [lindex $atom 2]
                dict set resdb $r1 resid    [string range [lindex $atom 3] end-3 end] ;# fix resid > 9999 to 0000, 0001 ...
                dict set resdb $r1 reindex  $res(new)
                dict set resdb $r1 frag     $frag
             }

           # Add atom info
             dict set resdb $r1 atoms [lindex $atom 4] [lindex $atom 5]
             
           # Search residue connectivity
             foreach atom2 $bonds {
                  
                  set r2 [lindex $atm2res $atom2]

                # Bridges | CYX terminal residues (e.g. 3HJ2)
                  if {[dict exists $backmap [lindex $atom 1] bridge [lindex $atom 4] [lindex $atm2resn $atom2] [lindex $atm2name $atom2]]} {

                      set b [dict get $backmap [lindex $atom 1] bridge [lindex $atom 4] [lindex $atm2resn $atom2] [lindex $atm2name $atom2]]

                      if {![info exists SS($r2,$r1)]} { set SS($r1,$r2) [list $r1 $r2 [lindex $b 0] [lindex $b 1]] }

                      dict set resdb $r1 name [lindex $b 2]
                      
                    # Relevant bonds in chain connectivity
                      if {[lindex $b 3] == "w"} {
                      
                          set dist [expr {$r2 - $r1}]
                          
                      #      close residue || circualar chain  
                          if { $dist ==  1 || $dist < -1 } { dict set resdb $r1 res+ $r2 }
                          if { $dist == -1 || $dist >  1 } { dict set resdb $r1 res- $r2 }
                      }
                      
                # Others
                  } else {

                      set dist [expr {$r2 - $r1}]
                   
                    #    close residue || circualar chain
                      if { $dist ==  1 || $dist < -1 } { dict set resdb $r1 res+ $r2 }
                      if { $dist == -1 || $dist >  1 } { dict set resdb $r1 res- $r2 }
                  }
             }
     }
     puts "Done \[[array size SS] bridge(s) found\]"
     array unset res

   # No backmapped residues
     if {[info exists nomap]} {
     
         puts "Skipping following residues:"
         foreach resname [array name nomap] { puts -nonewline "$resname " } ; puts ""
     }

     if {[dict size $resdb] < 1} {puts stderr "No residue to backmap"; return}

   # BACKMAP --------------------
   # Clean previous calculations
     if { [file exists .backmap]}            { puts -nonewline "Removing folder .backmap/... "    ; file delete -force -- .backmap          ; puts "Done!" }
     if { [file exists $opts(outname).pdb] } { puts -nonewline "Removing file $opts(outname).pdb "; file delete -force -- $opts(outname).pdb; puts "Done!" }

     puts "Runing backmapping procedure..."

   # Working dir
     file mkdir .backmap

   # MIN input file
   # NOTE:pmemd does not explicitly support vacuum simulations
   # http://archive.ambermd.org/201206/0552.html
     if {! $opts(nomin) } {
         
         set min [open ".backmap/min.in" w]

         puts  $min "Minimization\n&cntrl\nimin=1,ntmin=1,maxcyc=${opts(maxcyc)},ncyc=${opts(ncyc)},ntpr=50,ntxo=2,"
         puts  $min "ntb=0,igb=${opts(gbsa)},cut=${opts(cutoff)}"
         puts  $min "/"
         
         close $min
     }

   # OUTPUT
     set OUT [open "$opts(outname).pdb" a]
     
   # PDB format
     set PDBf "ATOM%7d%5s%4s%2s%4s%12.3f%8.3f%8.3f"

     puts "Processing [llength $opts(frames)] frame(s)..."
     
   # Progress
     puts -nonewline "         0    25   50   75   100 %\nProgress |"

     set bar    "||||||||||||||||||||" ;# Progress bar
     set tick    0                     ;# Starting tick mark in bar
     set num_fr  0                     ;# Frame number

   # Save trajectory to bkPDB
     foreach frame $opts(frames) {
     
       # Update frame
         $sys frame $frame
         incr num_fr

       # Backmap names and residues
         set PDB  [open ".backmap/backmap.pdb" w]

         set xyz  [$sys get {x y z}]
         set ndx  0                  ;# atom index
         set frag 1                  ;# fragment index

         foreach res [dict keys $resdb] {
           # Check new chain
             if {[dict get $resdb $res frag] != $frag} { puts $PDB "TER"; incr frag }
         
           # Build residue
             foreach atom [sirah_build $res $resdb $xyz] { ;# return {{ATOMNAME RESNAME {xyz}}...}
                 
                 incr ndx ; if {$ndx > 99999} {set ndx 0}

                 puts $PDB [ format $PDBf                        \
                                    $ndx                         \
                                    [lindex $atom 0]             \
                                    [lindex $atom 1]             \
                                    [dict get $resdb $res chain] \
                                    [dict get $resdb $res resid] \
                                    [lindex $atom 2 0]           \
                                    [lindex $atom 2 1]           \
                                    [lindex $atom 2 2]           \
                           ]
             }
         }

         puts  $PDB "TER\nEND"
         close $PDB

       # Minimaze structure
         if {! $opts(nomin)} {
         
          # AMBER input files
            set    leap [open ".backmap/backmap.leap" w]

            puts  $leap "# Load FF -------------"
            
            foreach cmd  ${forcefield(cmd)}  { puts $leap "source $cmd" }
            foreach parm ${forcefield(parm)} { puts $leap "loadAmberParams $parm" } ; # Mg2+/Ca2+
          
          # Workaround for circular DNA: no auto-terminal
            puts  $leap "addPdbResMap {
                         { 0 \"DG\" \"DG\" } { 1 \"DG\" \"DG\" }
                         { 0 \"DA\" \"DA\" } { 1 \"DA\" \"DA\" }
                         { 0 \"DC\" \"DC\" } { 1 \"DC\" \"DC\" }
                         { 0 \"DT\" \"DT\" } { 1 \"DT\" \"DT\" }
                        }"
            
            puts  $leap "m = loadpdb .backmap/backmap.pdb"
       
          # S-S Brigdes <<<<<<<<<<<<!
            foreach bond [array names SS] {

                  set  r1 [dict get $resdb [lindex $SS($bond) 0] reindex]
                  set  r2 [dict get $resdb [lindex $SS($bond) 1] reindex]

                  puts $leap "bond m.$r1.[lindex $SS($bond) 2] m.$r2.[lindex $SS($bond) 3]"
            }

         #  Save topology
            puts  $leap "saveAmberParmNetcdf m .backmap/backmap.prmtop .backmap/backmap.ncrst"
            puts  $leap "quit"
            close $leap
       
            exec $tleap -f .backmap/backmap.leap
            file rename -force -- leap.log .backmap/leap.log

          # Run Mimimization
          # In case TCL complains about "Error: Unable to create the sub-directory (/usr/tmp)"
          # mkdir /usr/tmp; chmod 777 /usr/tmp
            set run [concat $MPI $sander -O\
                           -i   .backmap/min.in\
                           -inf .backmap/min.inf\
                           -p   .backmap/backmap.prmtop\
                           -c   .backmap/backmap.ncrst\
                           -o   .backmap/backmap_min.out\
                           -r   .backmap/backmap_min.ncrst
                    ]
  
            eval exec $run
          
          # Workaround to avoid cpptraj Error: Atom ??? was assigned a lower molecule index than previous atom
          # http://archive.ambermd.org/201306/0453.html
          # The PDB is generated but the error exit status makes TCL execution to stop
            catch {exec cpptraj -p .backmap/backmap.prmtop -y .backmap/backmap_min.ncrst -x .backmap/backmap_min.pdb}
            
            file rename -force .backmap/backmap_min.pdb .backmap/backmap.pdb
         }
         
       # Trajectory
         puts  $OUT [format "MODEL %8d" $frame]
         set    BKM [open ".backmap/backmap.pdb" r]
         fcopy $BKM $OUT
         close $BKM
       
       # Print progress ------------
         set pg [expr {100*$num_fr/[llength $opts(frames)]}] ;# %Progress

         if { [expr {$pg/5}] > $tick } { puts -nonewline [string range $bar $tick [expr {$pg/5}]]; set tick [expr {$pg/5 +1}]; flush stdout }
       # ---------------------------
     };  puts ""

     close $OUT
     
   # Clean temporal folder
     file delete -force -- .backmap

   # Load backmap trajectory into VMD
     if {! $opts(noload) } { 
         puts -nonewline "Loading atomistic trajectory... "
         mol new  $opts(outname).pdb waitfor all
         puts "Done!"
     }
}


proc sirah_restype {} {

   # SIRAH DNA
     color restype DAX Nucleic_Acid
     color restype DTX Nucleic_Acid
     color restype DGX Nucleic_Acid
     color restype DCX Nucleic_Acid
     color restype AX5 Nucleic_Acid
     color restype TX5 Nucleic_Acid
     color restype GX5 Nucleic_Acid
     color restype CX5 Nucleic_Acid
     color restype AW5 Nucleic_Acid
     color restype TW5 Nucleic_Acid
     color restype GW5 Nucleic_Acid
     color restype CW5 Nucleic_Acid
     color restype AX3 Nucleic_Acid
     color restype TX3 Nucleic_Acid
     color restype GX3 Nucleic_Acid
     color restype CX3 Nucleic_Acid

   # SIRAH Proteins
     foreach aa {K Km R Hd He}             { foreach x {s n c a m} { color restype ${x}${aa} Basic    } }
     foreach aa {E D Z Sp Tp Yp}           { foreach x {s n c a m} { color restype ${x}${aa} Acidic   } }
     foreach aa {N Q S T Y Dh Eh Ka}       { foreach x {s n c a m} { color restype ${x}${aa} Polar    } }
     foreach aa {A C Cp X G I L M F P W V} { foreach x {s n c a m} { color restype ${x}${aa} Nonpolar } }

   # SIRAH Membrane
     color restype xPC Nonpolar
     color restype xPE Nonpolar
     color restype xPS Nonpolar
     color restype xMY Nonpolar
     color restype xPA Nonpolar
     color restype xOL Nonpolar

     color restype CMM Nonpolar
     color restype CPP Nonpolar
     color restype CPO Nonpolar
     color restype EPO Nonpolar
     color restype SPO Nonpolar

   # SIRAH Ions
     color restype NaW Ion
     color restype KW  Ion
     color restype MgX Ion
     color restype CaX Ion
     color restype ZnX Ion
     color restype ClW Ion

   # SIRAH Solvent
     color restype WT4 Solvent
     color restype WLS Solvent
     
     puts "SIRAH coloring methods were set"
}


proc sirah_macros {} {
   # Set some selection macros
     atomselect macro sirah_membrane {resname "xP[CESA]" xMY xOL CMM CPP CPO EPO SPO}

     atomselect macro sirah_nucleic  {resname "D[ATGC]X" "[ATGC][WX]5" "[ATGC][WX]3"}
     
     atomselect macro sirah_protein  {resname "[sncam][KREDNQSTYACZXGILMFPWV]" "[sncam][ED]h" "[sncam]H[de]" "[sncam][CSTY]p" "[sncam]K[am]"}
     atomselect macro sirah_basic    {resname "[sncam][KR]" "[sncam]H[de]" "[sncam]Km"}
     atomselect macro sirah_acidic   {resname "[sncam][EDZ]" "[sncam][STY]p"}
     atomselect macro sirah_polar    {resname "[sncam][NQSTY]" "[sncam][ED]h" "[sncam]Ka"}
     atomselect macro sirah_neutral  {resname "[sncam][ACXGILMFPWV]" "[sncam]Cp"}
     
     atomselect macro sirah_backbone {(name GN GC GO and sirah_protein) or (type PX KX KN and sirah_nucleic)}
     
     atomselect macro sirah_water    {resname WT4 WLS}
     atomselect macro sirah_ions     {resname KW NaW ClW MgX CaX ZnX}
     
     puts "SIRAH selection macros were set"
}


proc sirah_radii {args} {
   #
   # TCL script to set VdW radii of SIRAH atom types in VMD
   # Definitions will be applied to existing molecules and
   # to new loaded onces. Use in combination with an X-PLOR
   # PSF or AMBER topology file.
   #
   # SIRAH version [Dec 2019]
   #

     if {$args == "" } { sirah_help sirah_radii; return }
     
     lassign $args fname molid 
     
                         # atom-type
                         # GMX   AMB  sigma(nm)  element
     set sirah_atomtypes {
                           GNz   T1   0.55000    N
                           GOz   T2   0.55000    O
                           GNn   T3   0.40000    N
                           GOn   T4   0.40000    O
                           GN    ""   0.42000    N
                           GO    ""   0.42000    O
                           GC    ""   0.42000    C
                           Y1C   Y1   0.42000    C
                           Y2Ca  YA   0.42000    C
                           Y3Sm  Y3   0.42000    S
                           Y4Cv  Y4   0.42000    C
                           Y5Sx  Y5   0.42000    S
                           Y6Cp  Y6   0.42000    C
                           A1C   W1   0.35000    C
                           A1Cw  WC   0.35000    C
                           A2C   W2   0.35000    C
                           A3P   W3   0.35000    H
                           A4O   W4   0.35000    O
                           A5D   WD   0.35000    N
                           A5E   WE   0.35000    N
                           A7N   WN   0.35000    N
                           A8P   WH   0.35000    H
                           P1O   QO   0.40000    O
                           P1S   QS   0.41000    S
                           P2P   QP   0.40000    H
                           P3Cn  QC   0.40000    C
                           P3Cq  QK   0.40000    C
                           P4O   QX   0.40000    O
                           P5N   QN   0.40000    N
                           C1Ck  X1   0.40000    C
                           C2Cr  X2   0.42000    C
                           C3Cr  X3   0.40000    C
                           C4Cd  XD   0.40000    C
                           C4Ce  XE   0.40000    C
                           C5N   X5   0.45000    N
                           C6O   X6   0.45000    O
                           C7Nk  X7   0.55000    N
                           C7Nm  XM   0.55000    N
                           Y5Sz  YZ   0.42000    S

                           PL    ""   0.46327    P
                           PS    ""   0.46327    P
                           ETA   ET   0.35000    N
                           ""    WX   0.35000    N
                           YGL   GL   0.42000    C
                           P1E   E1   0.41000    O
                           P2E   E2   0.41000    O
                           Y2C   Y2   0.42000    C
                           Y3C   YT   0.42000    C
                           Y4C   YF   0.42000    C

                           PX    ""   0.46327    P
                           KX    ""   0.42906    C
                           KN    ""   0.33997    C
                           D2    ""   0.26698    H
                           D1    ""   0.32500    N
                           D6    ""   0.32500    N
                           J1    ""   0.32500    N
                           J2    ""   0.32500    N
                           M3    ""   0.32500    N
                           S4    ""   0.32500    N
                           S3    ""   0.32500    N
                           S2    ""   0.29599    O
                           M2    ""   0.29599    O
                           M4    ""   0.29599    O
                           J6    ""   0.29599    O

                           WT    ""   0.42000    X
                           WL    ""   0.65600    X

                           NaW   ""   0.58000    Na
                           KW    ""   0.64500    K
                           ClW   ""   0.68000    Cl
                           MgX   ""   0.40000    Mg
                           CaX   ""   0.40000    Ca
                           ZnX   ZX   0.40000    Zn
                         }

     foreach {GMX AMB sigma element} $sirah_atomtypes {
             
             set my_sel [atomselect $molid "type $GMX $AMB"]
             
             #       vdw radius = rm/2 = 2^(1/6) * sigma / 2
             $my_sel set radius [ expr {pow(2.0,(1.0/6))*$sigma*10.0/2.0} ]
             
             $my_sel set element $element
            
             $my_sel delete
     }

}

# ------------------------------------------------------------------------------------------

# Set SIRAH radii for existing molecules
  foreach mol [array name vmd_initialize_structure] {

        if ($vmd_initialize_structure($mol)) { sirah_radii vmd_initialize_structure $mol w }
  }

# Set SIRAH radii for new molecules
  trace variable vmd_initialize_structure w sirah_radii

  puts "SIRAH radii were set"

# Set SIRAH macros and residue types
  sirah_macros
  sirah_restype

# Build SIRAH backmapping library
  sirah_mapdb 

  puts "SIRAH Tool kit for VMD was loaded. Use sirah_help to access the User Manual pages"

# ------------------------------------------------------------------------------------------
```
