## Supporting file - scripts for "Protocols for Multi-Scale Molecular Dynamics Simulations: A Comparative Study for Intrinsically Disordered Amyloid Beta in Amber & Gromacs on CPU & GPU": ssmtx2png.R

```
#!/usr/bin/Rscript

#----------------------------------------------------------------------#
# SSMTX2PSF                                                            #
#----------------------------------------------------------------------#
# Script to create a PNG image from file ss.mtx generated by sirah_ss  #
#                                                                      #
# USAGE:   ./ssmtx2png.R [--args] [--help]                             #
#                                                                      #
# AUTHOR:  Matias Machado                                              #
# E-MAIL:                                      #
#                                                                      #
# This program is free software; you can redistribute it and/or modify #
# it under the terms of the GNU General Public License as published by #
# the Free Software Foundation; either version 2 of the License, or    #
# (at your option) any later version.                                  #
#                                                                      #
# This program is distributed in the hope that it will be useful,      #
# but WITHOUT ANY WARRANTY; without even the implied warranty of       #
# MERCHANTABILITY or FITNESS FOR A PARTICULAR PURPOSE. See the         #
# GNU General Public License for more details.                         #
#----------------------------------------------------------------------#

  Version = "version 1.0 [Nov 2015]"

# Functions ---------------------------------------------------------------
# Manual
  man = function(){
       cat("
       Script to create a PNG image from file ss.mtx generated by sirah_ss.
       The secondary structure information of each residue is plotted along
       time as a grid using colors from the R palette.

       Usage: ssmtx2png.R [--args] [--help]

       Arguments:
       --mtx=ss.mtx     - Secondary structure matrix
       --tu=us          - Time unit (ps,ns,us,ms,s,...)
       --dt=1e-04       - Time between consecutive frames
       --ttick=0.1      - Time tick marks
       --rtick=10       - Residue tick marks

       --H=purple       - Alpha-helix color
       --E=yellow       - Beta-sheet color
       --C=cyan         - Coil color

       --png=ssmtx.png  - Image output name
       --res=100        - PNG resolution (dpi)
       --w=10           - Image width (inches)
       --h=8            - Image height (inches)

       --version        - Print version and exit

       --help           - Print help and exit\n\n"
     )

    q(save="no")
  }

# Parse arguments
  parseArgs = function(args) {
            # http://www.r-bloggers.com/parse-arguments-of-an-r-script/
              args.S  = strsplit(sub("^--", "", grep("^--.*=.*",args,value=T)), "=")
              args.DF = as.data.frame(do.call("rbind",args.S))
              args.L  = as.list(as.character(args.DF$V2))
              names(args.L) = args.DF$V1
              return(args.L)
  }
# -------------------------------------------------------------------------

# Collect arguments
  args <- commandArgs(TRUE)

# Parse arguments (we expect the form --arg=value)
  if ("--help"    %in% args) {man()}
  if ("--version" %in% args) {cat(Version,"\n");q(save="no")}

  argsL = parseArgs(args)         ;# (L)ist of arguments

# Default Parameters ------------------------------------------------------
  if(is.null(argsL$mtx))  {matrix = "ss.mtx"   } else {matrix = argsL$mtx}
  if(is.null(argsL$tu))   {tu     = "us"       } else {tu     = argsL$tu }
  if(is.null(argsL$dt))   {dt     = 1e-04      } else {dt     = as.numeric(argsL$dt)}
  if(is.null(argsL$ttick)){ttick  = 0.1        } else {ttick  = as.numeric(argsL$ttick)}
  if(is.null(argsL$rtick)){rtick  = 10         } else {rtick  = as.numeric(argsL$rtick)}

  if(is.null(argsL$H))    {H.col  = "purple"   } else {H.col  = argsL$H}
  if(is.null(argsL$E))    {E.col  = "yellow"   } else {E.col  = argsL$E}
  if(is.null(argsL$C))    {C.col  = "cyan"     } else {C.col  = argsL$C}

  if(is.null(argsL$png))  {image  = "ssmtx.png"} else {image  = argsL$png}
  if(is.null(argsL$res))  {res    = 100        } else {res    = as.numeric(argsL$res)}
  if(is.null(argsL$w))    {width  =  10        } else {width  = as.numeric(argsL$w)}
  if(is.null(argsL$h))    {height =   8        } else {height = as.numeric(argsL$h)}

# Do some checks
  if (!file.exists(matrix)) {cat("Can't read input file", matrix,"use flag --help for access usage\n"); q(save="no")}
  if (tu == "us") {Time = expression( paste("Time (",mu,"s)") )} else {Time = paste("Time (",tu,")",sep="")}
  if (rtick <= 0) {rtick = 1}
# -------------------------------------------------------------------------

# Fast matrix reading
# Assign colClasses
  tab1row = read.table(matrix, header=FALSE, nrows = 1)
  classes = c("numeric",rep("character",dim(tab1row)[2] - 1))

# Import data
  SSMtx = as.matrix(read.table(file=matrix,header=FALSE,colClasses=classes))

  SSMtx[SSMtx == "H"] = 1
  SSMtx[SSMtx == "E"] = 2
  SSMtx[SSMtx == "C"] = 3

  class(SSMtx) = "numeric"

# Account for time and residue selection in sirah_ss
  t.max = dim(SSMtx)[1]
  r.max = dim(SSMtx)[2]
  t1    = SSMtx[1,1]         ;# Start time
  t2    = SSMtx[t.max,1]     ;# End time
  r1    = 1                  ;# Start residue
  r2    = r.max - 1          ;# End residue

# Start PNG device driver to save output to figure.png
  png(filename=image, width=width, height=height, bg="white", units="in", res=res)

  par(mar=c(4,4,1,1))

# Draw Color Matrix
  image( x=c(t1:t2)*dt            , xlab= Time    ,
         y=c(r1:r2)               , ylab="Residue",
         z=SSMtx[1:t.max,2:r.max] ,
         col=c(H.col,E.col,C.col) ,
         axes=FALSE
  )

# x axe
  axis(1,at=seq(0,t2*dt,by=ttick),las=1)

# y axe
  axis(2,at=seq(0,r2,by=rtick),las=1)
  axis(2,at=c(r1,r2),las=1,lwd=2)

# Turn off device driver (to flush output to png)
  dev.off()

# -------------------------------------------------------------------------
# EXIT
  q(save='no')
```
